# Tropical Origin, Global Diversification and Dispersal in the Pond Damselflies (Coenagrionoidea) Revealed by a New Molecular Phylogeny

**DOI:** 10.1101/2022.01.21.477207

**Authors:** B. Willink, J. Ware, E. I. Svensson

## Abstract

The processes responsible for the formation of Earth’s most conspicuous diversity pattern, the latitudinal diversity gradient (LDG), remain unexplored for many clades in the Tree of Life. Here, we present a densely-sampled and dated molecular phylogeny for the most speciose clade of damselflies worldwide (Odonata: Coenagrionoidea), and investigate the role of time, macroevolutionary processes and biome-shift dynamics in shaping the LDG in this ancient insect superfamily. We used process-based biogeographic models to jointly infer ancestral ranges and speciation times, and to characterise within-biome dispersal and biome-shift dynamics across the cosmopolitan distribution of Coenagrionoidea. We also investigated temporal and biome-dependent variation in diversification rates. Our results uncover a tropical origin of pond damselflies and featherlegs ∼ 105 Ma, while highligthing uncertainty of ancestral ranges within the tropics in deep time. Even though diversification rates have declined since the origin of this clade, global climate change and biome-shifts have slowly increased diversity in warm- and cold-temperate areas, where lineage turnover rates have been relatively higher. This study underscores the importance of biogeographic origin and time to diversify as important drivers of the LDG in pond damselflies and their relatives, while diversification dynamics have instead resulted in the formation of ephemeral species in temperate regions. Biome-shifts, although limited by tropical niche conservatism, have been the main factor reducing the steepness of the LDG in the last 30 Myr. With ongoing climate change and increasing northward range expansions of many damselfly taxa, the LDG may become less pronounced. Our results support recent calls to unify biogeographic and macroevolutionary approaches to increase our understanding of how latitudinal diversity gradients are formed and why they vary across time and among taxa.

---

One of the most striking features of species diversity is the non-uniformity of its global distribution. Tropical regions of the world generally have higher species richness compared to temperate regions, a pattern known as the Latitudinal Diversity Gradient (LDG) (Hillebrand 2004). The LDG has fascinated naturalists from centuries, including both Darwin (1859) and Wallace (1876, 1878), who saw the climatic stability of the tropics as a factor hindering extinction, and believed that stronger biotic interactions in the tropics could accelerate speciation. Wallace also suggested that climatic fluctuations, such as Pleistocene glaciations, were more pronounced in temperate regions, providing a limited opportunity for diversity to accumulate in these areas compared to the tropics (Wallace 1878). These early biogeographic hypotheses were further developed by ecologists and evolutionary biologists throughout the twentieth century (Dobzhansky 1950; Fischer 1960; MacArthur 1969), and today, the mechanisms responsible for the LDG remain a highly active area of research (e.g. Hanly et al. 2017; Rabosky et al. 2018; Gaboriau et al. 2019; Saupe et al. 2019; Meseguer and Condamine 2020).

Current hypotheses to explain the causes of the LDG can be broadly categorized into those that emphasize ecological effects on diversification rates vs. those that emphasize time for speciation in different regions and the contingencies of evolutionary history (Pianka 1966; Mittelbach et al. 2007; Wiens 2011). At one end of this dichotomy, the tropics are portrayed as ecology-fuelled species pumps, due to higher speciation rates (Wiens 2007), lower extinction rates (Weir and Schluter 2007; Pyron 2014) or both (Jablonski et al. 2006; Pyron and Wiens 2013; Rolland et al. 2014). The tropics have also been suggested to be the sources of most evolutionary novel lineages due to their greater geographical area (Rosenzweig and others 1995; Belmaker and Jetz 2015), greater niche heterogeneity (Moritz et al. 2000; Salisbury et al. 2012), stronger biotic interactions (Dobzhansky 1950; Schemske et al. 2009), or higher mean temperature (Wright et al. 2006; Brown 2014). As Darwin and Wallace already anticipated, tropical diversity might also be buffered from the extinction that occurs in temperate regions because of greater climatic instability (Dynesius and Jansson 2000; Jablonski et al. 2006; Hawkins et al. 2007). In contrast, studies at the other end of the dichotomy have proposed that the amount of time available for diversification might be the predominant cause of variation in regional species richness, whether aligned or contrary to the LDG (Stephens and Wiens 2003; Stevens 2006; Economo et al. 2018; Miller and Román-Palacios 2021).

Understanding how ecological effects and historical contingencies shape regional diversification dynamics is therefore fundamental for a full understanding of the LDG, and it is crucial to critically evaluate the empirical evidence for these contrasting alternatives (Wiens 2011). However, regional biotas are not only shaped by varying speciation and extinction rates, but also by dispersal (Salisbury et al. 2012; Smith et al. 2012; Antonelli et al. 2015; Jablonski et al. 2017). A lineage may disperse to a distant region with a similar biome, maintaining the LDG, or move to an ecologically novel area (i.e. a colder or warmer biome), thus altering the steepness of the LDG. Moreover, diversification and dispersal can act interdependently (Goldberg et al. 2011). For example, ecological limits to diversity, if present, could potentially reduce both speciation rates and incoming dispersal (Etienne et al. 2019). Similarly, ecological specialization and niche partitioning may result in fast speciation locally, but trade-off with the evolution of dispersal ability (Jocque et al. 2010; Salisbury et al. 2012; Polato et al. 2018). Despite a potential key role of dispersal in molding the LDG, a prevailing focus on diversification dynamics may have curbed our understanding of the relative roles of each of these processes shaping regional biotas (Roy and Goldberg 2007).

To date, only a few studies have examined the combined effects of speciation, extinction and dispersal in shaping diversity gradients (Antonelli et al. 2015; Rolland et al. 2015; Spano et al. 2016; Jablonski et al. 2017; Owens et al. 2017). The majority of these studies target clades with a relatively rich fossil record, which also enables process-based methods for inferring time-calibrated phylogenies, such as the fossilized birth-death process and total evidence approaches (Ronquist et al. 2012; Heath et al. 2014). However, most organisms, including insects, do not fossilize well (Wills 2001; Kidwell and Holland 2002), and therefore other process-based methods are necessary to calibrate molecular clocks and infer ancestral geographic ranges (e.g. Tiley et al. 2020). Biogeographic dating approaches have been long used as an alternative to fossil dating, but until recently, most of these methods relied on node calibrations (Kodandaramaiah 2011; De Baets et al. 2016) and have consequently failed to consider the joint probability of alternative diversification and dispersal scenarios (Ho et al. 2015).

Landis (2017) introduced a data-dependent generative model, in which speciation times and ancestral geographic states are jointly estimated over discretely defined epochs. Crucially, this approach used paleogeographical data to explicitly consider how the Earth’s dynamic past, through tectonic movements, has influenced the availability of dispersal routes for terrestrial animals. More recently, Landis et al. (2021) implemented a similar time-stratified approach to model within-biome dispersal and biome shifts, building on paleoecological literature and thus examining the role of phylogenetic niche conservatism in shaping biogeographic histories. These studies illustrate how empirical models of dispersal barriers can be used in lieu of fossils to implement process-based calibrations. They also show how time-dependent process-based models can be used to reconstruct ancestral diversity patterns, such as the LDG, while accounting for paleoclimatic changes. Here, we capitalised on these developments to investigate the phylogenetic and biogeographic history of a globally-distributed group of insects with a scant insect fossil, the damselfly superfamily Coenagrionoidea (Carle et al. 2008; Bybee et al. 2021).

Our target clade includes two families: the pond damselflies (family Coenagrionidae) and their close relatives the featherlegs (family Platycnemididae). Coenagrionidae is a large family, distributed across all continents except Antarctica, and with species that are at extremes of trait and niche diversity. For instance, the family includes the tropical-rainforest phytotelmata-breeding helicopter damselflies, previously considered a separate family due to their morphological distinctiveness (Dijkstra et al. 2014; Toussaint et al. 2019). At the other extreme, Coenagrionidae also includes species are restricted to Subartic fens and marshes in the type genus *Coenagrion* (Paulson 2009, 2011). In the Nearctic alone, where most systematic niche data has been assembled, heterogeneity in habitat preference is striking. Species vary vastly in their affinities to forested areas, range sizes and breeding habitats (Abbott et al. 2022). Platycnemididae, the other family in Coenagrionoidea, is composed by primarily lotic (i.e. inhabiting fast-moving water bodies) and paleotropical taxa, with some species reaching Northern Europe and East Asia (Boudot and Kalkman 2015; Dijkstra 2017).

Combined, the two families in Coeangrionoidea account for over a fourth of all Odonata species. Their ancient history (Kohli et al. 2021; Suvorov et al. 2021), rich ecological diversity (Abbott et al. 2022) and high dispersal potential as relatively large volant insects (May 2019), makes pond damselflies and featherlegs an excellent group to study the processes responsible for global diversity patterns, such as the LDG. However, few molecular studies have examined the phylogenetic relationships within Coenagrionoidea (Carle et al. 2008; Dijkstra et al. 2014; Waller and Svensson 2017; Toussaint et al. 2019; Bybee et al. 2021; Kohli et al. 2021), and a single dated and species-level phylogeny currently includes this clade (Waller and Svensson 2017). This species-level phylogeny nonetheless used phylogenetically-unsupported taxonomy as backbone. There is clearly a need for an updated phylogeny of Coenagrionoidea with denser taxon sampling, and for modern phylogenetic comparative analyses to uncover large-scale diversity patterns and their drivers in this ancient group of insects.

Here, we present the most extensively sampled phylogeny of pond damselflies and their relatives to date. We include over 35% of the approximately 1,805 species, and thereby substantially increase the phylogenetic coverage of sampled taxa compared to previous studies (approximately 7% of all species in Dijkstra et al. 2014; and 23% in Waller and Svensson 2017). Recent phylogenomic studies have produced a robust and dated backbone for the order Odonata, which we used to inform the origin time, over 100 Ma, of the most recent common ancestor of pond damselflies and featherlegs (Kohli et al. 2021; Suvorov et al. 2021). We combine this root-age prior with an empirical paleogeographic model (Landis 2017), genetic data, and distribution range data of extant taxa, to simultaneously infer the divergence times and geographic distributions of ancestral lineages. We also use this time-calibrated phylogeny to quantify temporal and geographic patterns of diversification, and to jointly model how within-biome dispersal and biome shifts have shaped regional diversity throughout the history of the clade. Our results provide a comprehensive overview of the macroevolutionary history of this insect group and shed new light on the question of how speciation, extinction and lineage movements influence global diversity patterns.

## Materials and Methods

Here, we summarize the main features of the phylogenetic analyses performed in this study. For a detailed description of data collection, sequence alignment and the statistical approach used for biogeographic dating, diversification, and biome-shift analyses, see the Supplementary Material. We mention here the most relevant prior distributions. For complete information on priors and parameter proposals, see the Extended Methods and accompanying scripts. All novel sequence data used in this study will be uploaded to NCBI Genbank. The sequence alignment, distribution data and code necessary to reproduce our results will be uploaded on Dryad (https://datadryad.org/stash) and are currently available on Github (https://github.com/bwillink/Damsel_Phylo). Phylogenetic analyses were partly conducted on the computer clusters at the LUNARC Center for Scientific and Technical Computing, Lund University, Sweden, and the Uppsala Multidisciplinary Center for Advanced Computational Science (UPPMAX).

### Sequence Data and Tree Topology Inference

We sampled a total of 669 taxa in the Coenagrionoidea superfamily, 556 of which are currently classified as Coenagrionidae and 113 as Platycnemididae (Table S1, S2). In a recent phylogenomic study, the small family Isostictidae was recovered as sister to the clade including both Coeangrionidae and Platycnemididae (Bybee et al. 2021). As we did not sample Isostictidae, we hereafter use the superfamily name Coenagrionoidea to refer to Coenagrionidae (pond damselflies) and Platycnemididae (featherlegs) only. Samples were collected in the field (n = 142 species), or obtained from specimens in museum and private collections (n = 362 species), and publicly available data on NCBI (https://www.ncbi.nlm.nih.gov/genbank/) (n = 194 species). For each species, we obtained sequence data for up to five loci (cytochrome oxidase subunit I, 16S ribosomal DNA, histone 3, phenyl-methyltransferase and 28S ribosomal DNA). Our molecular data consists of 1,695 new sequences, 564 previously published sequences, and 105 unpublished sequences from the Naturalis Biodiversity Center, The Netherlands. We conducted all phylogenetic analyses using probabilistic graphical models and Bayesian inference for parameter estimation, using RevBayes v. 1.0.7, v. 1.0.12, and v. 1.1.1 (Höhna et al. 2014, 2016).

Our first analysis aimed to infer the topology of the Coenagrionoidea phylogeny. We used a rooted uniform tree prior and a partitioned data scheme accounting for substitution rate variation among loci and among codon positions of coding sequences (Table S3). We specified a General Time-Reversible (GTR) model of molecular evolution for each partition, and accounted for rate heterogeneity among sites by assuming gamma-distributed rates. Finally, we enforced the monophyly of each of the two families within Coenagrionoidea (Coenagrionidae and Platycnemididae), as supported by recent large-scale studies (Kohli et al. 2021; Bybee et al. 2021; Suvorov et al. 2021). To do this, we imposed a single topological constraint of the root node of the tree, which was necessary as preliminary analyses did not adequately resolve this well-established ancestral relationship. The maximum *a posteriori* probability (MAP) tree was used to summarize phylogenetic inference.

### Biogeographic Dating with Empirical Paleogeography

Next, we simultaneously inferred speciation times and ancestral distributions across the Coenagrionoidea tree using the empirical paleogeographic model and statistical approach developed by Landis (2017). Instead of relying on internal node calibrations, this approach jointly models ancestral dispersal and speciation events as conditional on historical shifts in oceanic barriers and the distribution ranges of extant taxa. To use biogeography in node-age estimation, we recorded species distributions from the literature, museum collections, IUCN Red List assessments and reputable web sources for all extant taxa included in this study (Table S4). Due to the low spatial resolution and coarse-grained nature of available data, species distributions were registered as presence or absence from administrative geographical regions (countries and states/provinces/territories for countries spanning multiple biogeographic areas; see below).

In the model developed by Landis (2017), dispersal can occur by three modes (short-medium- and long-distance), which are differentially limited by oceanic barriers. The empirical paleogeographic model captures historical changes in these water barriers due to continental drift (Landis 2017). Thus, paleogeography determines the probability of transitions between biogeographic states, for each dispersal mode and for each discrete time interval (epoch) since the maximum plausible root age of the tree. This time-heterogeneous process describing the probability of dispersal events in turn informs branch length estimation in absolute time (Landis 2017). For example, ancestral dispersal leading to a disjunct distribution in the present is inferred to occur with higher probability during time intervals (epochs) in which the currently disjunct land masses were adjacent, and hence connected by all three dispersal modes, than when they were separated by small or large water barriers, which would have then required medium- and long-distance dispersal respectively.

Damselflies are aquatic insects with a terrestrial adult phase. They require fresh water habitats to breed and for larvae to develop (Corbet 1999). As damselflies cannot survive in or on salt water, it is highly likely that their dispersal has been historically constrained by oceanic barriers. Yet, Coenagrionoidea has a cosmopolitan distribution at present, ranging across all continents except for Antarctica and Greenland (Table S4, Fig. 1). Unlike the larger dragonflies (suborder Anisoptera), in which long-distance and sometimes trans-oceanic migration has been described (Wikelski et al. 2006; Anderson 2009; Troast et al. 2016), the majority of damselflies are small, weak fliers and tend to be more sedentary. For these biological reasons, we considered that a global biogeographic model of changes in dispersal routes due to continental drift would be a suitable approach to explore historical biogeography in these damselflies.

**Figure 1.**
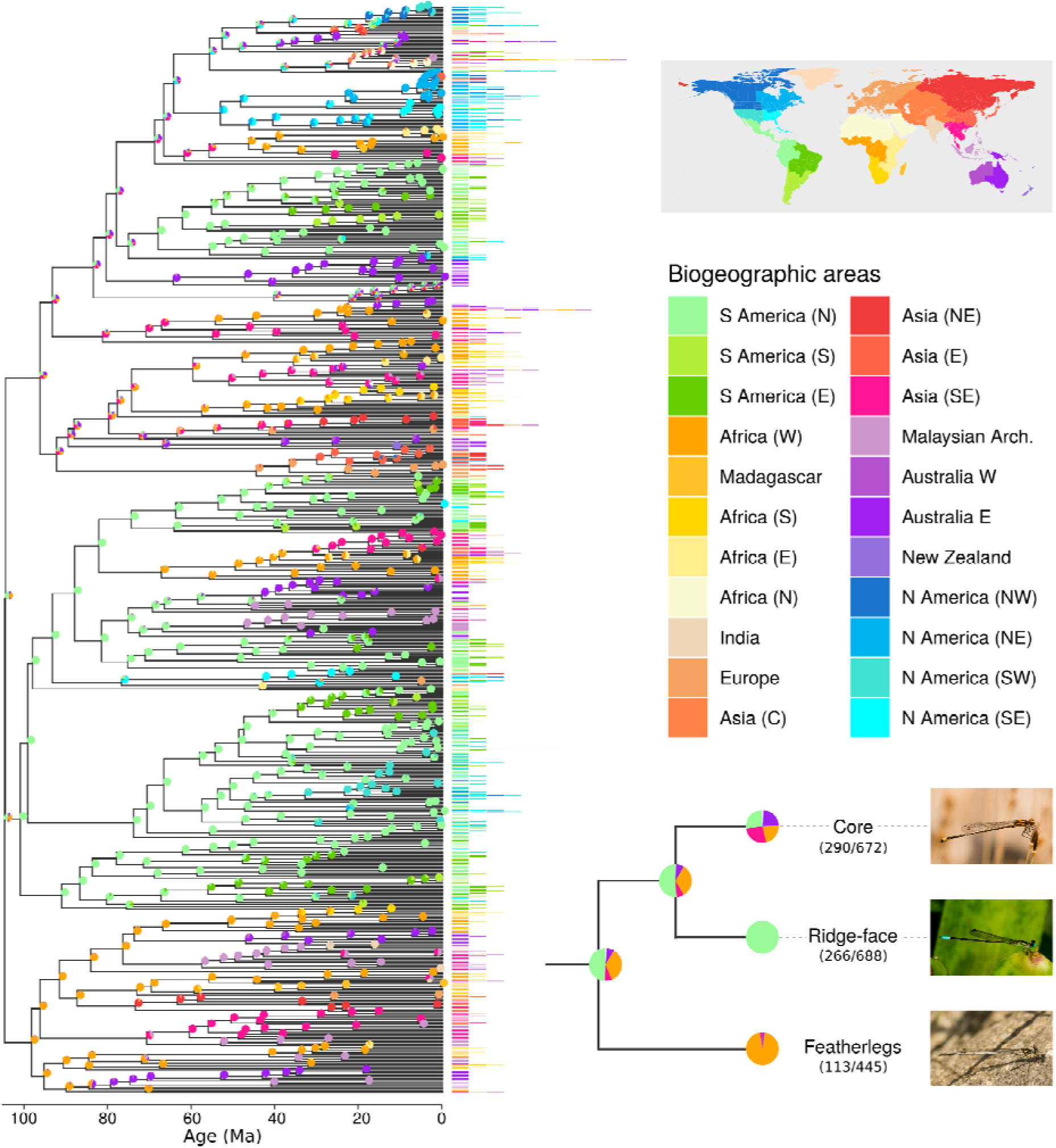
Inferring the biogeographic history of pond damselflies and their relatives the featherlegs (superfamily Coenagrionoidea). Speciation times and ancestral ranges were jointly estimated using a time-heterogeneous continuous-time Markov model, based on empirical paleogeography, an informed root-age prior and molecular and distribution data of 669 extant taxa. The empirical paleogeographic model in Landis (2017) was used to determine dispersal edges between 25 geographic areas across the globe (Greenland, Antarctica (E) and Antarctica (W) not shown in the colour legend). Phylogenetic inference was summarized using a maximum *a posteriori* (MAP) tree. The distribution areas occupied by each of the extant taxa are shown at the tips of the tree. Pies show the proportion of each ancestral state in a random sample of 1000 posterior trees. The cladogram in the lower right shows the inferred ancestral ranges for the most recent common ancestor of Coenagrionoidea, and for each of the three major clades in the superfamily. The fraction of taxa sampled is shown in parenthesis. One representative species is illustrated for each clade: *Acanthagrion adustum* (‘core’), *Leptagrion elongatum* (ridge-face’), *Platycnemis pennipes* (featherlegs). Photos: EIS.

The empirical paleogeographic model in Landis (2017) consists of 25 biogeographic areas (Fig. 1, Table S5) and 21 epochs between 240 Ma and the present (see Table S2 and Fig. S8 in Landis 2017). We combined this model with a root-age prior based on recent large-scale phylogenomic studies of major families in the whole order Odonata (Kohli et al. 2021; Suvorov et al. 2021). Suvorov et al. (2021) inferred a backbone phylogeny for Odonata using a supermatrix of 1,603 gene orthologs and fossil calibrations with 20 crown fossil constraints. Kohli et al. (2021) used the sequences from nearly 3,000 protein-coding genes and 17 fossil calibrations to reconstruct and date the phylogeny of Odonata. These studies date the origin of Coenagrionoidea to ∼107 and ∼116 Ma respectively. We placed a normally distributed root-age prior with a mean of 120 and standard deviation of 20 to reflect this prior knowledge on the origin time of Coenagrionoidea. The normal distribution was truncated between 240 Ma, the approximate age of the most recent common ancestor (MRCA) of all Odonata (Misof et al. 2014; Suvorov et al. 2021) and 40 Ma, the minimum age according to the study with the youngest age estimate for Coenagrionoidea (Waller and Svensson 2017).

To assess the impact of our strongly informed root-age prior and the empirical paelogeographic model, we conducted two additional analyses to contrast our results. First, we ran an identical model but with a weakly informed root-age prior, in which the MRCA of Coenagrionoidea occurred with equal probability between 240 and 40 Ma. Second, we conducted an analysis with a uniform prior on the root age, in this case bounded between 240 and 0 Ma, and ignoring the empirical paleogeographic data. This latter analysis hence excluded all empirical data (except the inferred origin time of Odonata) that could inform absolute divergence times and time-heterogeneous dispersal rates between biogeographic areas, and was used to rule out the possibility that divergence times were influenced by artefacts arising from model construction. For a comparison of this paleogeographic dating approach to a dating analysis informed by fossil constraints and a pure-birth model see the Supplementary Material.

### Diversification through Time and Space

We quantified the temporal and geographic dynamics of diversification in Coenagrionoidea using the MAP tree from the dating analyses above. We used an episodic birth-death (EBD) model (Stadler 2011; Höhna 2015) to test for time-dependent diversification rate shifts since the MRCA of Coeangrionoidea. EBD analyses assume that diversification rates change through time in a piece-wise constant fashion, following a process of Brownian motion. Thus for each epoch, rates are centred around the estimate in the previous epoch going backwards in time (Höhna 2015; Magee et al. 2020). We used an EBD model with 20 equal-length time intervals, and accounted for missing taxa in each of the main clades of Coenagrionoidea (Fig. S4).

We used a Hidden State-Dependent Speciation and Extinction (HiSSE) model to estimate the diversification rates in three latitude-delimited biomes (tropical, warm-temperate and cold-temperate) while accommodating background diversification-rate heterogeneity across different branches of the tree (Beaulieu and O’Meara 2016). Heterogeneity in background diversification was taken into account by including a hidden trait with two states, which differed in their effects on diversification. We first compared the relative fit of this biome-dependent HiSSE model against a biome-independent model, which also accounted for diversification rate heterogeneity using a hidden trait with two states. We then conducted three biome-dependent diversification analyses under different data assumptions.

Of the 669 taxa in our phylogeny 31 had an unknown biome state, either because their species identity was uncertain, or because their distribution range was not covered by the paleobiome model used in the biome shift and within-biome dispersal analysis (see below). An additional 99 species occupy ranges that extend across tropical and warm-temperate biomes (n = 46), across warm-temperate and cold-temperate biomes (n = 47) or across all three biome types (n = 6). We treated these taxa in two ways in separated analyses. In the main text, we present the results of our HiSSE model assigning taxa the coldest biome they inhabit. Because the common ancestor of Coenagrionoidea was tropical (see Results), biome states here indicate the extent to which a lineage has become established in a novel temperature niche. In the Supplementary Material, we show these results are qualitatively unchanged if we treat the widely distributed taxa as ambiguous between the biomes they currently occupy. Finally, we also conducted the biome-dependent HiSSE analysis using the Coenagrionoidea tree from the weakly informed dating analysis as input, and show these results are also qualitatively similar.

For all HiSSE analyses, biomes were assigned to each taxa based on present-day geography, using distribution data at the level of administrative areas (Table S4), and biome reconstructions for the last 5 Ma (Jolly et al. 1998; Salzmann et al. 2008; Otto-Bliesner et al. 2020). We note that the HiSSE model is time-constant and does not consider how paleoclimatic change influence the geographic distribution of tropical and temperate biomes.

### Biome shifts and within-biome dispersal

To investigate the role of lineage movements on the formation and dynamics of the LDG, we jointly modelled dispersal events within biomes and biome shifts in Coenagrionoidea using the approach developed in Landis et al. (2021). Similarly to our dating analysis, the likelihood of a dispersal event or biome shift is informed by the availability of suitable routes, but here, routes are potentially shaped by both land and biome continuity. For instance, if dispersal is limited by biome affinities, a lineage adapted to tropical rainforests in South East Asia should have a better chance of reaching Europe during periods of the Earth’s climatic history in which tropical rainforests were widespread in both regions, compared to more recent icehouse periods, when tropical rainforests had withdrawn from Europe. In contrast, if dispersal is unconstrained by ecological features, the same lineage movement would only depend on geographic continuity, which has been largely maintained between Europe and South East Asia throughout the last 100 Myr. Crucially, because the extent to which these geographic and ecological structures influence ancestral range shifts is unknown, Landis et al. (2021) treated them as free parameters that are estimated from the data as part of the biome shift and dispersal process (see Extended Methods).

As in our dating analysis, the dynamics of biome availability and connectivity are represented in this model by time-dependent graphs, informed by literature data. Here, we build on the paleobiome graphs used in Landis et al. (2021), which include three broad biome categories (tropical, warm-temperate, and cold-temperate) across six geographic regions (Southeast Asia, East Asia, Europe, South America, Central America, and North America), and eight time intervals over the last 106 Myr. We focus on the last 106 Myr, based on the inferred origin time of Coenagrionoidea in our dating analysis (see Results). We nonetheless expand the empirical model of Landis et al. (2021) to accommodate more of the global distribution of Coenagrionoidea, by adding three geographic regions (Africa, Australia, and India) that harbour a large fraction of pond damselfly and featherleg diversity (Fig. S2-S3). Species with distributions outside these main geographic regions (e.g. Pacific islands and Central Asia endemics, n = 32) were treated as missing data (i.e. included in the tree but lacking a biome-region state). Similarly, extant species with ranges that span two or more geographic regions or biomes (n = 151), were treated as ambiguous between their current states (e.g. a tropical rainforest species ranging from Central America to South America was coded as ambiguous between the Tropical+SAm and Tropica+CAm states, see also Table S7).

We used stochastic character mapping to sample lineage proportions through time and evolutionary histories in the dispersal and biome shift model. Following Landis et al. (2021), we also used these sampled histories to quantify the frequency of events (biome shifts and region shifts) and event series (see below) throughout the evolution of Coenagrionoidea. *Event series* represent different patterns of consecutive transitions between biomes and regions. First, we quantified instances in which both biomes and regions shifted in a lineage and recorded whether the biome shift preceded the dispersal event (*biome-first* series) or *vice versa* (*region-first* series). Then, we examined consecutive changes in biomes and regions separately. A *reversal* corresponds to a series of events with the same starting and ending state. For instance, a *biome reversal* occurs when a tropical species shifts to a warm-temperate biome and then back to the tropics. A *flight* is a series of two events across three different states. For example, a lineage that originates in Africa and then disperses to India and South East Asia would represent a *region flight*. We compute and report the proportion of each event and *event series* type across our posterior sample of evolutionary histories (see Extended Methods).

In this and previous analyses, we report the posterior mean (PM) and 95% highest posterior density (HPD) intervals for parameter estimates. For proportions of events and *event series* we also report 80% HPD intervals. Phylogenetic inferences and parameter estimates were summarized using R (R Core Team 2021). Plots we produced with the R packages *ggplot2* (Wickham 2016) and *ggtree* (Yu et al. 2017).

## Results

### Topology inference

The relationships reconstructed here are largely congruent with previous works (Dijkstra et al. 2014; Toussaint et al. 2019; Bybee et al. 2021). We fixed the relationship between the two families of Coenagrionoidea, Platycnemididae (featherlegs) and Coenagrionidae (pond damselflies) as resolved in previous genomic-scale backbone phylogenies (Suvorov et al. 2021; Bybee et al. 2021; Kohli et al. 2021). Within Platycnemididae, our tree recovers the previously well-supported relationships between the subfamilies Platycnemidinae, Disparoneurinae and Calicnemiinae, while the phylogenetic placement of Allocnemidinae, Idiocnemidinae and Onychargiinae (*sensu* Dijkstra et al. 2014) remains elusive across studies (Fig. S5a). Nonetheless, the generic composition of each of these subfamilies (Fig. S6) is consistent with Dijkstra et al. (2014), who sampled a relatively large fraction (∼10%) of all fetherleg species.

Within Coenagrionidae, the relationships between four main clades, the ‘core’ pond damselflies (*sensu* Dijkstra et al. 2014), the genus *Argia* (Rambur, 1842), the subfamily Protoneurinae (*sensu* Dijkstra et al. 2014) and the strict-sense ‘ridge-face’ clade (Dijkstra et al. 2014), have remained contentious among recent studies (Fig S5b). Our results are congruent with the most recent backbone phylogeny for Odonata, which used targeted genomics to resolve ancestral relationships in the order (Bybee et al. 2021) (Fig S5b). In both phylogenies, *Argia* and the strict-sense ‘ridge-face’ pond damselflies form a clade, in turn sister to Protoneurinae (Fig S5b). We hereafter refer to this more inclusive clade as ‘ridge-face’ pond damselflies to distinguish it from the the ‘core’ clade identified in Dijkstra et al. (2014).

Three large and well-sampled genera of ‘core’ pond damselflies (*Ischnura*, *Enallagma* and *Acanthagrion*) form a clade that is sister to *Agriocnemis* (incl. *Mortonagrion* and *Argiocnemis*, see Extended Results). We recovered these previously supported relationships in our phylogenetic inference (Fig. S5c). We also recovered a close relationship between *Coenagrion* and *Pseudagrion* (and allied genera) relatively to the clade described above and consistently with previous studies (Fig. S5c). However, our results differ from previous phylogenies in that it places other relatively small South Pacific genera (*Xanthocnemis*, *Austrocoenagrion*, and *Stenagrion*) within the same clade as *Coenagrion* and *Pseudagrion*, albeit with low support (Fig. S5c).

Finally, in the strict-sense ‘ridge-face’ pond damselflies (i.e. excluding *Argia* and Protoneurinae), relationships between the most distinct genera are also largely congruent among studies. The Neotropical *Telebasis* and the Paleotropical *Ceriagrion* form a clade separately from the strict rainforest dwellers *Mecistogaster*, *Metaleptobasis* and *Teinobasis* (Fig. S5d). In both our study and in Toussaint et al. (2019), who focused on ridge-face pond damselflies, the Neotropical genus *Metaleptobasis* and the Neotropical helicopter damselflies (including *Mecistogaster*) show closer affinity, while in Dijkstra et al. (2014), *Mecistogaster* and the Paleotropical *Teinobasis* appear more closely related (Fig. S5d).

Out of 115 genera included in this study, 45 are represented by a single species, either because the genus is monotypic (n = 22, i.e. having only one described species) or because material was available for only one species (n = 23). Of the remaining 70 genera, 50 were recovered as monophyletic, in most cases with high posterior probability (Table S6, median = 1.00, min = 0.55, max = 1.00). A total of 20 genera were thus recovered as paraphyletic. Of these, five pose phylogenetic relationships that challenge current taxonomy supported by morphology (*Acanthagrion*, *Cyanallagma*, and *Oxyagrion*) or earlier molecular studies (*Erythromma*, *Platycnemis*), 10 are supported by recent studies that also inferred paraphyly (*Agriocnemis*, *Coeliccia*, *Elattoneura*, *Indocnemis*, *Ischnura*, *Mecistogaster*, *Nesobasis*, *Pseudagrion*, *Prodasineura*, and *Teinobasis*), and five lack previous studies with taxonomic sampling that would enable detection of paraphyly (*Aciagrion*, *Enallagma*, *Mortonagrion*, *Psaironeura*, *Telebasis*). We discuss these potentially paraphyletic genera in the Extended Results.

### Dated Phylogeny and Ancestral Biogeography

Our biogeographic dating analysis using a strongly informed root calibration returned a mean age estimate for the MRCA of Coenagrionidae and Platycnemididae of 105 Ma (95% HPD interval = 64 – 145 Ma; Fig. S26). In contrast, applying a broad uniform prior on the root age resulted in a much younger estimate of 67 Ma (95% HPD interval = 40 – 118 Ma; Fig. S26). An analysis excluding empirical paleogeographic data produced, as expected, a flat posterior root age distribution between 240 and 0 Ma (Fig. S26). Finally, an analysis excluding paleogeographic data and relying on fossil constraints for node-dating returned a similar root-age estimate as the strongly informed biogeographic dating analysis (Table S10). Relatively shallow nodes (i.e. at the level of genera) had older age estimates in the strongly informed biogeographic dating analysis than in previous studies of comparable clades (Table S10). In contrast, our fossil-based approach generally resulted in young age estimates for shallow nodes compared to other large-scale studies, and in agreement with genus-level studies (Table S10). Despite large dating differences between the two biogeographic analyses in this study, biogeographic trends were qualitatively similar between strongly informed and weakly informed models (Fig. S27-S33). Here, we focus on results under the strongly informed prior, and present the full results under the weakly informed prior in the Supplementary Material.

Extant pond damselflies and featherlegs are globally distributed (Fig. 1). Our ancestral biogeographic inference indicated that Coenagrionoidea most likely originated in either Northern South America or Western Africa (Fig. 1, S27), as these two ancestral ranges combined accounted for 88% of the posterior samples of the root state. The two extant lineages descending from this common ancestor had contrasting biogeographic histories. The featherlegs (family Platycnemididae) originated in Western Africa (PS = 0.959, Fig. 1, S27) and likely dispersed throughout the Old World after multiple departures from Africa (Fig. 1, S28-S29). The pond damselflies (family Coenagrionidae) are in turn composed of two clades with distinct dispersal dynamics. The ‘ridge-face’ clade of pond damselflies originated with high probability in Northern South America (PS = 0.999; Fig. 1, S27), and thereafter remained largely neotropical, with a few successful dispersal events to the Nearctic (in *Argia* and *Nehalennia*), tropical Africa and Asia (in *Ceriagrion*) and presumably also across the Pacific (*Melanesobasis*, *Teinobasis*, *Amphicnemis* and allied genera) (Fig. 1, S30-S31). Finally, the ‘core’ pond damselflies have a much more dynamic dispersal history, particularly in the genus *Ischnura* (Fig. 1, S32-S33), and currently occupy all available biogeographic areas and many oceanic islands not considered in this analysis (e.g. *Megalagrion* in Hawaii; Fig. S32-S33). While the origin of ‘core’ pond damselflies is also tropical, the geographic range of their MRCA remained elusive (Fig. 1, S27).

### Temporal Dynamics of Diversification

Our EBD analysis showed a decline in the rate of speciation since the origin of Coenagrionoidea, particularly between 85 - 55 Ma (Fig. 2a). This analysis also detected a recent but modest increase in extinction (Fig. 2b). The combined effects of speciation and extinction dynamics resulted in a relatively constant net diversification rate in the early history of Coenagrionoidea, followed by a period of declining diversification (∼ 85 - 55 Ma) and another period of constant to slowly declining diversification (∼55 Ma to the present) (Fig. 2c). As a result of increasing extinction over the last 5 Myr, the 95% HPD interval of diversification over this period includes the possibility of net diversity loss near the present (Fig. 2c). EBD dynamics modelled on the dated phylogeny under a weakly informed root-age prior are qualitatively similar to the patterns described above (Fig. S34).

**Figure 2.**
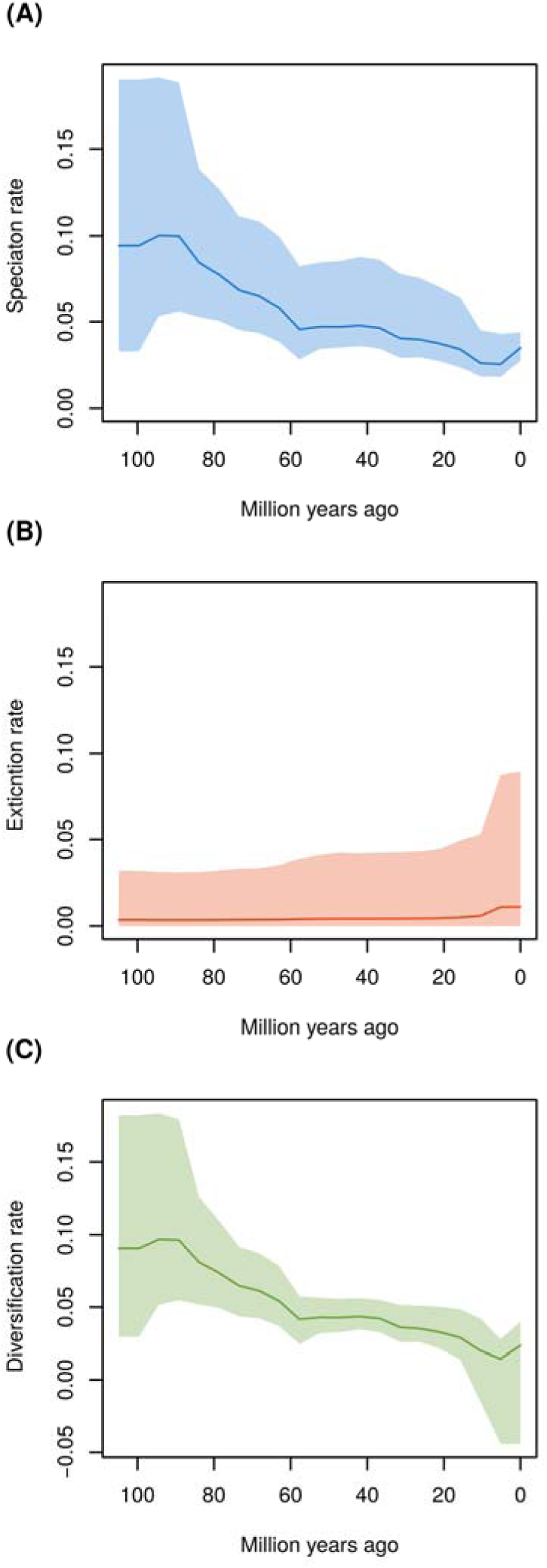
Diversification rate changes since the MRCA of Coenagrionoidea: a) speciation, b) extinction and c) net diversification. Diversification rates were estimated on the maximum *a posteriori* (MAP) tree from the biogeographic phylogenetic dating analysis using a root age calibration based of previous phylogenomic studies (Kohli et al. 2021; Suvorov et al. 2021). Changes in diversification rates were modelled under an episodic birth-death (EBD) process, over 20 equal-length time intervals and assuming autocorrelation among consecutive time intervals (see Extended Methods).

### Biome-dependent Diversification

The biome-dependent HiSSE model was very strongly supported over an alternative model with only biome-independent rate heterogeneity (Table S11). As in the paleogeographic dating analysis, the HiSSE model inferred a tropical distribution with high certainty for the MRCA of pond damselflies and featherlegs (Fig. S35). Following dispersal from the tropics, both speciation and extinction were accelerated in cold-temperate biomes, while warm-temperate biomes prompted relatively low speciation and high extinction (Table S12; Fig. 3a,b). As a result, net diversification rates were similar across biomes (Table S12; Fig. 3a,b). In contrast, lineage turnover occurred at a faster pace in both temperate biomes compared to the tropics (Table S12; Fig. 3d).

**Figure 3.**
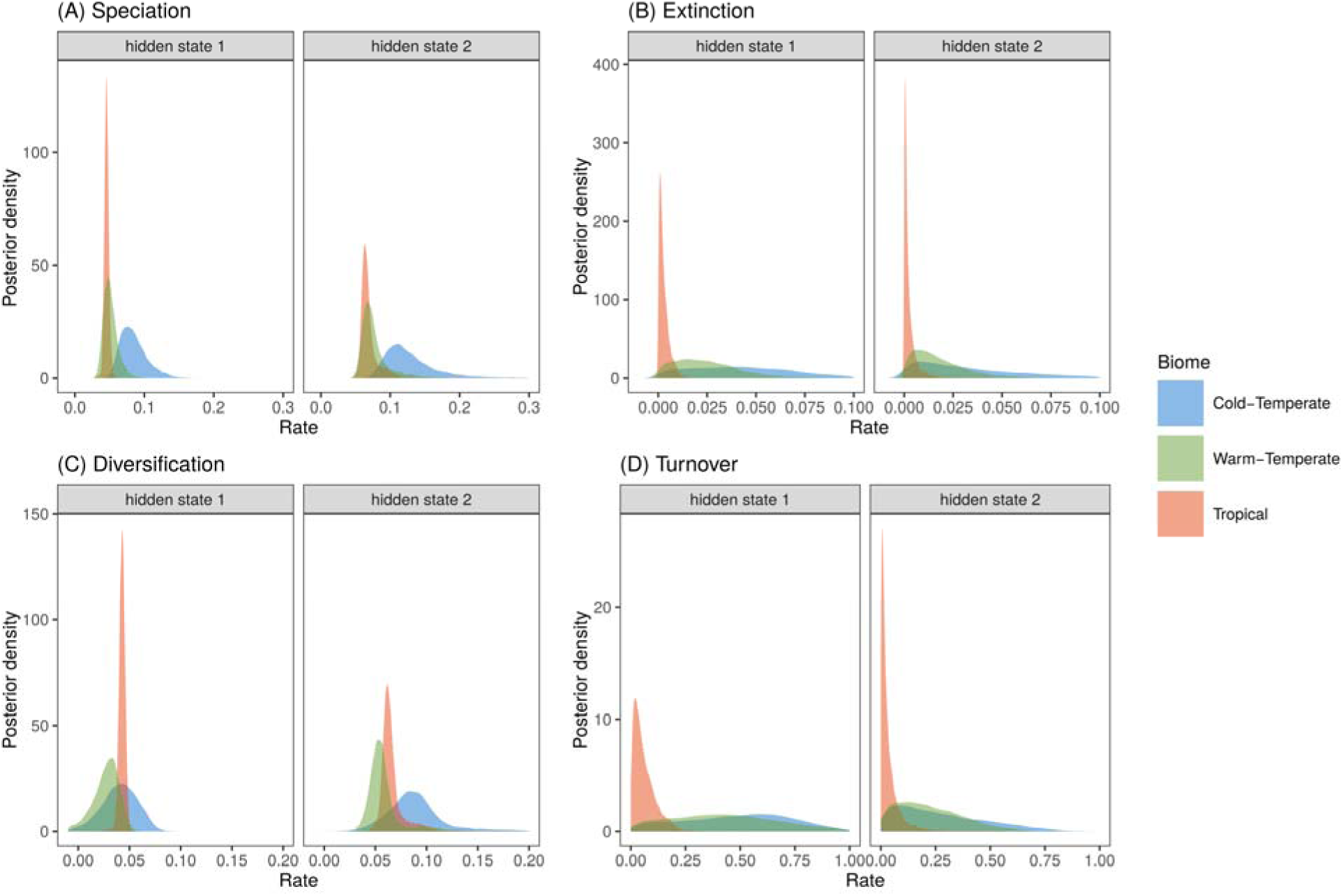
Latitudinal effects on diversification of pond damselflies and their relatives the featherlegs (superfamily Coenagrionoidea). (a) Speciation and (b) extinction rates at tropical and temperate latitudes were estimated using a Hidden-State Dependent Speciation and Extinction (HiSSE) model that accounts for background heterogeneity in diversification rates across the phylogeny, by including a hidden trait with two states. (c) Diversification was calculated as the net difference between speciation and extiction and (d) turnover was calculated as the ratio between extinction and speciation. The histograms show the posterior distribution of parameter estimates.

Background diversification rates distinguished the ‘core’ pond damselflies and the ‘ridge-face’ genera *Argia* and *Ceriagrion*, all with relatively high diversification, from the remaining ‘ridge-face’ pond damselflies and the featherlegs, with with relatively low diversification (Fig. S35). Results of the HiSSE analysis using the MAP tree under a weakly informed root-age prior and using a character matrix that included ambiguous states for wide ranging taxa were qualitatively similar (Table S13-S14; Fig. S36-S39). A model run without data recovered prior distributions, as expected (Fig. S40).

### Dispersal and biome shift dynamics

Estimated weighing parameters for geographical (w_G_= 0.003) and biome features (w_B_= 0.975), indicated that lineage movements in Coenagrionoidea are strongly influenced by biome affinities, while land connectivity independently of ecological features has a marginal effect on shaping ancestral ranges. It is thus not surprising that ancestral state reconstructions differed somewhat from those in our biogeographic dating model, which considered paleogeographic but not paleoecological history. In both models, we obtained strong support for a tropical ancestor of pond damselflies and featherlegs (Fig. 1, 4, S41). However, unlike the biogeographic dating model, the ancestral region with highest posterior probability in the biome-shift model was South East Asia (Fig. 4, S41). Featherlegs and ‘core’ pond damselflies were also inferred as most likely originating in tropical South East Asia, in contrast to the biogeographic dating analysis, while ‘ridge-face’ pond damselflies originated with highest posterior probability in tropical South America, consistent with the biogeographic dating analysis (Fig. S41).

**Figure 4.**
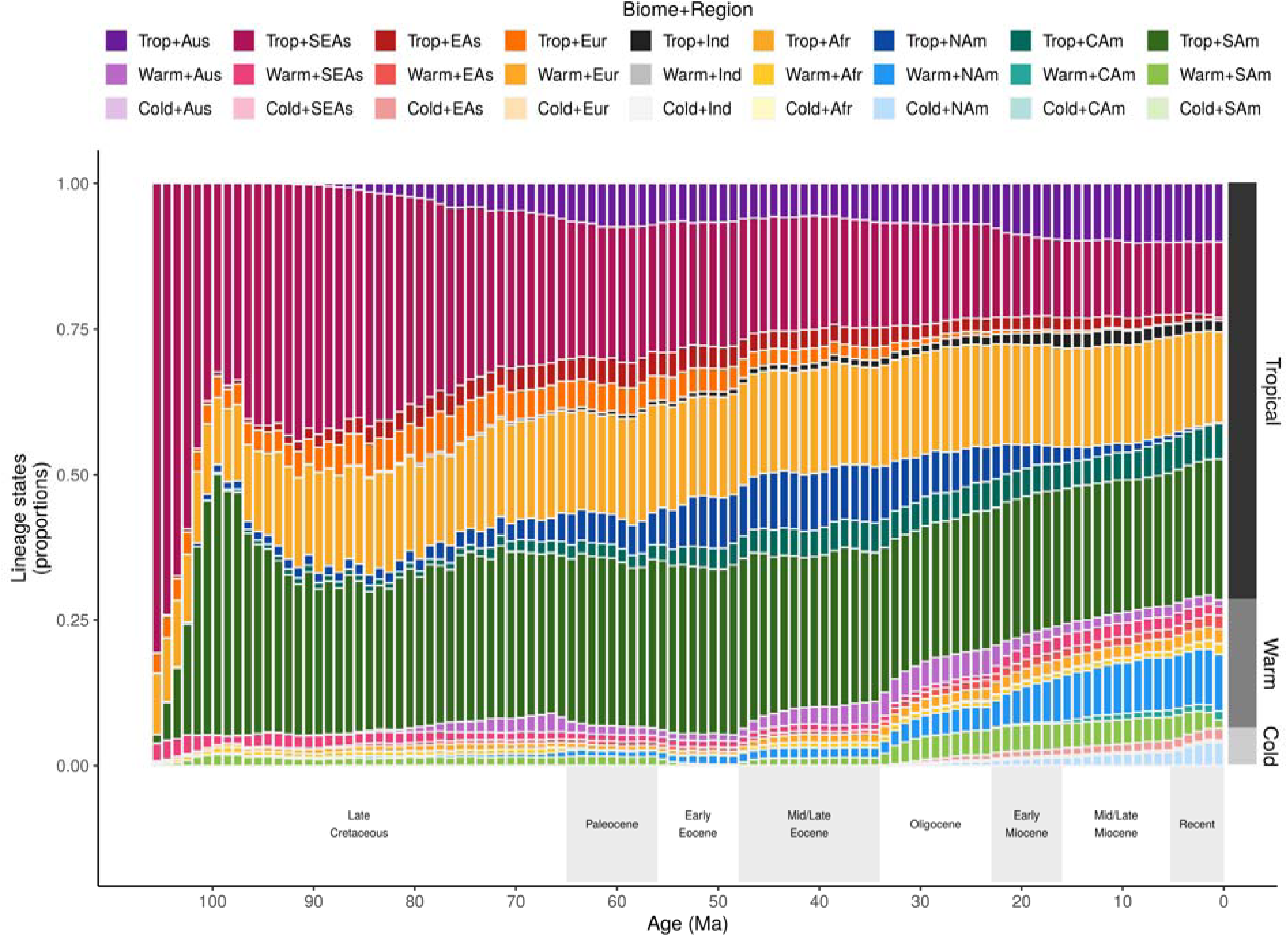
Proportion of Coenagrionoidea lineages in each biome-region state and through time. Lineage proportions were computed from stochastic histories sampled in a time-heterogeneous model of biome-region shifts (Landis et al. 2021). The dark-grey to light-grey bars on the right show the proportions of the three biome states in extant taxa sampled in this study (669 out of ∼1,800 species).

Clade-wide biome-region dynamics over time point to three major themes in the ancestral distribution of Coenagrionoidea. First, lineage diversity has been overwhelmingly tropical throughout history, with a rapid increase of taxa in South America shortly after the origin of the clade (Fig. 4). Second, a cooling trend during the Oligocene coincided with a slow and steady increase in temperate diversity, which continues to the present and is most notable in warm-temperate North America (Fig. 4). Finally, inference of relative biome-shift rates suggested a dynamic diversity exchange between tropical and warm-temperate biomes, indicated by a moderate rate of dispersal from tropical to warm-temperate biomes and fast dispersal back to the tropics (Fig. S42). In contrast, cold-temperate biomes gain diversity and lose diversity at comparable and slow rates (Fig. S42).

The dynamic diversity exchange between tropical and warm-temperate biomes also emerged within regions, but the timing of these events seems to have varied across the planet. In South America, increased shifts between tropical and warm-temperate biomes followed the Oligocene cooling trend (Fig. 5a), as warm-temperate biomes became more available in the region (Fig. S2-S3). In North America and Northern Eurasia (Fig. 5a,b), the exchange occurred mainly before the end of the Eocene, after which tropical biomes began to decline in these regions (Fig. S2-S3). Finally, in the rest of Afroeurasia (Africa, India and South East Asia) and in Australia, shifts between tropical and warm-temperate biomes were comparatively stable over time (Fig. 5c).

**Figure 5.**
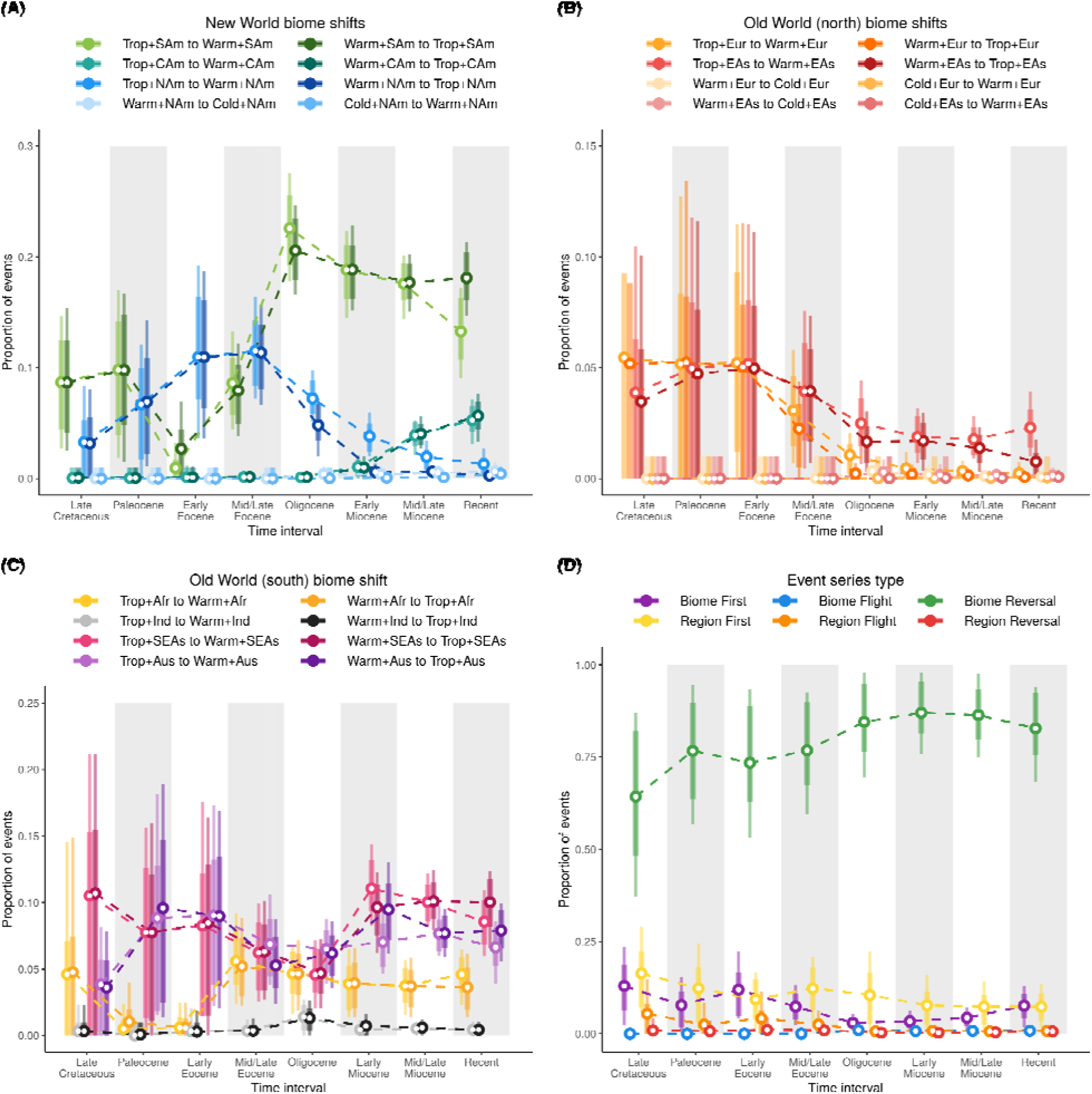
Proportion of biome-shift events (a-c) and event series (d) through time. Event proportions were estimated from stochastic histories sampled in a time-heterogeneous model of biome-region shifts. Circles represent posterior means, bars represent 80% HPD intervals, and lines represent 95% HPD intervals. See Methods for descriptions of the event series types.

We note that stochastically mapped character histories were largely congruent with regionally available biomes (Fig. S43, see also Extended Methods). The largest proportion of lineages with mismatched states (i.e. lineages with biome states that are regionally unavailable) occurred at the start of the Miocene, when nearly 6% of Coenagrionoidea lineages were inferred to occupy tropical biomes in North America (Fig. 4), but tropical biomes in this region became marginal during the transition between the Oligocene and Miocene, in the empirical paleobiome model (Fig. S2-S3). As the proportion of tropical lineages in North America decreased during the Miocene (Fig. 4), congruence between inferred and available states increased (Fig. S43).

Consistent with estimates of biome-shift rates (Fig. S42) and relatively frequent shifts between tropical and warm-temperate biomes within regions (Fig. 5a-c), we found that *biome reversals* (e.g. from tropical to warm-temperate and back to tropical) are by large the most frequent *event series* across Coenagrionoidea (Fig. 5d). In contrast, *region reversals* and *biome flights* are extremely rare, while *region flights* and series involving both biome shifts and dispersal events occur at intermediate frequencies (Fig. 5d). Interestingly, the proportion of *region flights* dropped to almost zero during the Oligocene (Fig. 5d), when our empirical paleobiome model encodes a sharp decline in the continuity of tropical biomes across the Earth (Fig S2-S3). Together, *event series* reconstructions further indicate that movements within relatively warm areas of the globe (i.e. tropical and warm-temperate biomes) are the most prominent features in the historical biogeography of Coenagrionoidea.

## Discussion

In this study, we used a combination of biogeographic and multi-locus sequence data from extant taxa, empirical paleogeographic and paleobiome models (Landis 2017; Landis et al. 2021), and dating priors from recent phylogenomic analyses (Kohli et al. 2021; Suvorov et al. 2021). Our goal was to understand how biome-dependent diversification and dispersal shaped the current latitudinal diversity gradient in the most speciose superfamily of damselflies. While inferences about the age of Coenagrionoidea are notably sensitive to prior information (Fig. S26), our ancestral biogeographic reconstructions based on empirical paleogeography are robust to node-age uncertainty (Fig. S27-S33). Biogeographic inferences that additionally consider ecological niche conservatism, by modelling the effects of biome continuity on lineage movements, were partly in conflict with the paleogeographical-only model, especially in deeper nodes of the tree. Nonetheless, all of our analyses concurred on a tropical origin of Coenagrionoidea, followed by rapid dispersal throughout the world’s tropics, and more recently into high latitudes.

Despite their early global presence, diversification rates in Coenagrionoidea have declined since their evolutionary origin (Fig. 2). Diversification dynamics also vary between biomes, with cold-temperate lineages undergoing the fastest speciation and extinction, but with limited consequences for net diversification (Fig. 3). In contrast, biome-shifts, via lineage movements and paleoclimatic change, have resulted in a growing representation of temperate lineages, particularly as the Earth’s climate became cooler during the Oligocene (Fig. 4). Taken together, our results point to an early start of tropical diversification as the main driver of the LDG in Coenagrionoidea. Diversity gains due to biome shifts and accelerated speciation in temperate regions have counteracted the LDG, yet these effects may be at least partially swamped by a tendency of damselfly lineages to stay in relatively warm areas of the planet and a relatively fast pace of extinctions outside the tropics.

Our topology inference of Coenagrionoidea recovered phylogenetic relationships that are largely congruent with other recent molecular phylogenetic studies (Dijkstra et al. 2014; Toussaint et al. 2019; Bybee et al. 2021). However, our study also calls for the revision of multiple generic classifications based exclusively on morphological evidence, and for further phylogenetic studies of a few particularly understudied clades (see Extended Results). We note that the root node of the tree, which marks the divergence between the pond damselfly and featherleg families was not resolved in our preliminary analysis, despite being strongly supported in multiple phylogenomic studies (Kohli et al. 2021; Suvorov et al. 2021; Bybee et al. 2021). In our preliminary analysis, the featherlegs were recovered as a monophyletic group, but their relationships to the main clades within the pond damselflies were uncertain. We therefore imposed a topological constraint to enforce the monophyly of each family in our final phylogenetic analysis. This resulted in strongly supported relationships between pond damselfly clades, in agreement with the most comprehensive backbone phylogeny of Odonata to date (Fig. S5c, Bybee et al. 2021). Nonetheless, enforcing the monophyly of pond damselflies may have also contributed to underestimating branch lengths at the base of the family (see Table S10). Ancestral introgression and incomplete lineage sorting in the early history of Coenagrionoidea are likely causes for phylogenetic discordance, and their effects tend to be aggravated in analyses based on concatenated loci (Kubatko and Degnan 2007). Moreover, both processes have been pervasive in the deep history of Odonata, leaving a genetic footprint within and between damselfly superfamilies (Suvorov et al. 2021).

Our biogeographic dating analysis, combining empirical paleogeography with prior information on the root age of Coenagrionoidea, resulted in an estimated time for the MRCA of 105 Ma (Fig. S26). This estimate is only slightly younger, yet more uncertain, than the age of Coenagrionoidea in the phylogenomic studies used for calibration (Kohli et al. 2021; Suvorov et al. 2021). Our estimates of internal node ages under the strongly informed prior are also consistent with most fossil records, except for specimens attributed to Platycnemididae, which have been dated to ∼ 99 Ma (Zheng et al. 2017), before the inferred origin of crown Platycnemididae in this analysis (∼ 80 Ma) and in previous studies (Table S10). Compared to shallow (genus-level) nodes dated in previous studies, inferred ages in this analysis are relatively older (Table S10). In contrast, the dating analysis with a weakly informed root-age prior, returned a younger origin time for Coenagrionoidea (Fig. S26), which was also younger than estimates in previous studies (Table S10). However, this latter analysis produced age estimates for shallow nodes that were usually closer to estimates in previous studies (see for example *Ischnura*, *Enallagma*, *Nesobasis*, *Nehalennia* and *Platycnemis* in Table S10).

These results underscore the challenges in dating phylogenies of groups of organisms with a scant fossil record, and the limitations of simple models of dispersal and paleogeography. The statistical approach developed by Landis (2017) is appealing for its ability to jointly sample dispersal histories and diversification events. However, the computational challenges to the likelihood calculation over a large space of states currently prohibits incorporating critical complexity into the paleogeographic model (Ree and Sanmartín 2009). A terrestrial graph at a global scale seems suitable for widely distributed damselflies, which require fresh water habitats to breed (Corbet 1999). Nonetheless, many genera of pond damselflies are endemic to specific archipelagos, while some taxa are widely distributed across continents (Fig. 1; Table S4). In the present analysis, island endemics are subsumed into larger biogeographic areas or coded as missing data (see Supplementary Material), and distribution ranges spanning two or more areas are treated as character-state uncertainty in extant taxa. A model that incorporates branch heterogeneity in dispersal rates and captures the more fine grained paleogeographic events that shaped the distribution of island endemics would likely improve biogeographic dating in this and other diverse groups of insects that vary in their ease of dispersal (e.g. Jones et al. 2016; Landis et al. 2018) and tend to leave a relatively sparse fossil record (Wills 2001). Yet, such extensions, as well as the use of large genomic data sets, under a fully Bayesian approach are currently constrained by computational limitations.

Despite these challenges to node dating, both biogeographic dating analyses under alternative root-age priors indicate that the MRCA of pond damselflies and featherlegs originated in either Northern South America or Western Africa (Fig. 1; S27). These areas were separated by narrower water barriers at the presumed origin of Coenagrionoidea (particularly under the strongly informed root-age prior) and have preserved tropical rainforests throughout the Cenozoic, notwithstanding climatic change (Morley 2007; Couvreur et al. 2011). In contrast, the biome-shift and dispersal analysis inferred the ancestral range of Coenagrionoidea in tropical South East Asia with relatively high (> 80%) posterior probability (Fig. S41). Two other deep nodes in the Coenagrionoidea phylogeny, the MRCAs of the featherlegs and ‘core’ pond damselflies, were also incongruent between models, while shallower nodes were generally consistent. An important distinction between the analyses is that our dating model assumed that all land habitats are equally suitable for a dispersing lineage, whereas the biome-shift model allowed ecological affinities to influence lineage movements. Another feature that differentiated the models is that the paleogeography of the Malaysian Archipelago was treated separately from South East Asia in the dating analysis, while both areas were treated as a single region in the biome-shift analysis. Our results therefore highlight the sensitivity of ancestral range reconstructions to assumptions embedded in dispersal graphs. While more complex models (e.g. with spatial and temporal resolution of dispersal graphs) could improve these inferences, we think that improving our mechanistic understanding of range dynamics to inform these models will be crucial to achieve robust inferences of ancestral ranges in ancient and widely distributed clades such as Coenagrionoidea (Jønsson et al. 2016).

The early evolution of Coenagrionoidea was characterised by fast diversification, followed by a decline in the rate of accumulation of new species (Fig. 2). This is a common feature of dated phylogenies (Morlon et al. 2010), which has been subject to considerable controversy and which can be attributed to several possible causes including statistical artefacts (reviewed in Moen and Morlon 2014). One possibility is that regional diversification slows down over time because niche space is reduced as species accumulate, eventually reaching some ecological limit (Rabosky 2009; Etienne et al. 2011; Etienne and Haegeman 2012). While plausible if contemporaneous taxa co-occur and compete for finite resources (Herrera-Alsina et al. 2018), this explanation for a decline in diversification with time seems less likely for non-adaptive radiations (Rundell and Price 2009; Czekanski-Moir and Rundell 2019). Some relatively young radiations in pond damselflies and other odonates have resulted from non-ecological speciation mechanisms, such as sexual selection or sexual conflict, and many closely related taxa are therefore only weakly ecologically differentiated from each other (McPeek and Brown 2000; Siepielski et al. 2010; Svensson 2012; Svensson et al. 2018). These features of damselfly communities suggest that niche partitioning does not strongly constrain speciation, although such weak niche differentiation might of course impact the longevity of newly formed lineages (see below). Moreover, our finding of abundant range reversals from warm-temperate to ancestral tropical biomes (Fig. 5) further argues against ecological diversity limits at a regional scale.

An apparent slowdown in diversification may also arise from statistical artefacts or sampling biases. For example, underparametrized models of molecular evolution can lead to underestimation of branch lengths, particularly in older branches of the tree (Revell et al. 2005). As diversification rate estimation ultimately relies on the distribution of branch lengths across the tree, this type of model misspecification can result in an artificial slowdown of diversification over time. An artificial slowdown can also be a product of incomplete lineage sampling, especially when taxon sampling is non-random (Cusimano and Renner 2010; Brock et al. 2011). However, in this study we have used methods that explicitly incorporate sampling fractions of specific clades as a measure to counter this bias (Höhna 2015). Nonetheless, even when models are correctly specified and sampling of extant taxa is accounted for, the “push of the past” can produce an apparent slowdown of diversification over time (Budd and Mann 2018). This survivorship bias occurs because we can only sample extant descendants from lineages which had high enough speciation rates as to have survived to the present. Deep branches in the tree that diversified more slowly, and would have contributed to a more constant estimate of diversification rates over time, are more likely to have gone extinct over the 105 Myr of Coenagrionoidea evolution, thereby leaving no descendants from which to infer their lower diversification rates. The “push of the past” remains a challenge to estimating temporal diversification dynamics when we lack information of whether and how many entire clades have gone extinct (Budd and Mann 2018).

The present distribution of pond damselflies, as many other clades, is characterised by higher species richness near the equator compared to temperate areas (Fig. 4) (Hillebrand 2004; Kinlock et al. 2018). This LDG in extant Coenagrionoidea is striking, considering that tropical lineages, particularly from India, South East Asia, and the Malaysian Archipelago, are probably underrepresented in the largest collections that contributed to this study. With this potential caveat in mind, our results indicated that none of the hypotheses based on accelerated speciation or reduced extinction in the tropics (see Introduction) is a likely explanation for why most extant damselfly species occur at low latitudes. While taking background diversification heterogeneity into account (Beaulieu and O’Meara 2016), we found similar rates of net diversification between the tropical and temperate regions of the globe (Fig. 3c). Nonetheless, speciation and extinction dynamics do tend to differ across latitudes, with both rates being highest in cold-temperate biomes, and thus cancelling each other out. As a consequence, both warm-temperate and cold-temperate lineages tend to be relatively short lived and replace one another at a faster rate than tropical lineages do, resulting in higher turnover rates outside of the tropics (Fig. 3d). Several studies have uncovered a similar pattern of increased lineage turnover in the temperate region or in harsher non-tropical environments (Weir and Schluter 2007; Botero et al. 2014; Pyron 2014; Harvey et al. 2020).

Faster lineage turnover at higher latitudes may come about in at least two ways. First, paleoclimatic fluctuations (e.g. DeConto et al. 2008) can cause the fragmentation of distribution ranges, a phenomenon that is particularly pronounced at higher latitudes (Jansson and Dynesius 2002). By repeatedly creating isolated refugia, climate fluctuations may have episodically fostered allopatric speciation (Weir and Schluter 2004; Milá et al. 2007; Sánchez-Ramírez et al. 2015; Morales-Barbero et al. 2018). However, the same recurrent climatic events can also increase extinction rates, either because range fragmentation reduces population size, or because hybridization and competitive exclusion drive extinction upon secondary contact (Dynesius and Jansson 2000; Barnosky 2005; Botero et al. 2014). Furthermore, climate fluctuations at higher latitudes might open up novel ecological opportunities that prompt speciation, by constantly renewing the availability of unoccupied niches (Schluter 2016). High environmental harshness in these novel environments at high latitudes might also result in the predominance of viability selection driven by abiotic environmental conditions, potentially increasing extinction risk and thereby accelerating lineage turnover (Haldane 1937; Gomulkiewicz and Holt 1995; Cutter and Gray 2016).

Whether it is driven by ecological factors (Cutter and Gray 2016; Schluter 2016), or non-ecological mechanisms (McPeek and Gavrilets 2006; Svensson et al. 2006; Siepielski et al. 2018), ephemeral speciation in temperate regions does not seem to have a net impact on diversification. Our results therefore highlight the role of the historical origin of Coenagrionoidea in the tropics as one of the main causes of the present-day LDG. This historical early start of diversification in the tropics has maintained the LDG despite an overall decrease in speciation rate (Fig. 2), and a net gain of lineages through biome shifts into temperate regions, particularly over the last 30 Myr (Fig. 4). Our findings that biome affinities strongly influenced ancestral distribution ranges, and that temperate lineages tend to return to the tropics (Fig. 5, S42), suggest that dispersal out of the tropics has been limited by tropical niche conservatism, even in periods when warm and cold-temperate biomes were widely available. These results support previous studies finding stronger niche conservatism in tropical lineages (Smith et al. 2012; Pyron and Wiens 2013), and underscore how the potential for adaptive evolution into colder biomes is likely a primary factor governing the steepness of the gradient (Rangel et al. 2018).

As a result of an early start of diversification in the tropics, followed by similar net diversification rates across latitudinal regions, the Coenagrioniodea LDG is expected to persist, unless an external input of diversity to the temperate areas reverses this general pattern. Continuing the trend of the last 30 Myr, a trickle of tropical lineages reaching cold biomes could slowly contribute to a shallower LDG (Fig. 4). However, in a much shorter term, the smoothing of environmental gradients due of global warming (Loarie et al. 2009), might release adaptive constraints for lineages dispersing into higher latitudes. Examples of such rapid northward expansions have already been documented in Odonata (Paulson 2001; Grewe et al. 2013; Lancaster et al. 2015). The role of dispersal shaping the future steepness of the LDG thus warrants further investigation.

## Conclusion

The damselfly superfamily Coenagrionoidea originated in tropical areas, where most of its diversity is currently found (Fig. 1). An increasing number of studies have revealed the importance of both ecology and historical biogeography in producing regional differences in diversity across distinct clades of plants and animals (e.g. Rolland et al. 2014; Antonelli et al. 2015; Jablonski et al. 2017). Here, we have shown that in pond damselflies and their relatives, early tropical diversification followed by relatively slow shifts into warm-temperate and later cold-temperate biomes established the latitudinal diversity gradient observed today. Faster ecological speciation in temperate regions is unlikely to compensate for this time lag in the future, as speciation might be more ephemeral at higher latitudes (Fig. 3). However, climate change and lineage movements have also contributed to an increase of Coenagrionoidea diversity in cold biomes, currently at high latitudes (Fig. 4). Our results reveal a complex interplay between history, macroevolutionary processes and dispersal shaping global diversity patterns.

## Acknowledgements

We are grateful to Hanna Bensch, Robin Pranter, Tammy Ho, Chiara de Pasqual and Yassin Tschinda for their ceaseless efforts to sample specimens in field and obtain sequence data in the lab. This study would have been impossible without the generosity and time of all the avid odonatologists and museum curators who made hundreds of specimens available to us and helped us with taxonomic identifications. We are particularly grateful to Bill Mauffray and Paul Skelly at the Florida State Collection of Arthropods, Oliver Flint and Floyd Shockley at the Smithsonian Institute, Darren Ward at the New Zealand Arthropod Collection, Darío Lijtmaer at the Argentinian Museum of Natural Sciences, and KD Dijkstra at the Naturalis Biodiversity Center. Many others contributed with samples and/or taxonomic expertise (Table S2). Specimens were collected with permits from the Guyana Environmental Protection Agency (Reference No. 010715 BR001 and 012615 SP: 002), the Secretaría de Ambiente y Desarrollo Sustentable, Argentina (Ref No. 11084/16) and from the Cameroon Ministry of Forestry and Wildlife (Ref. No. 0000034). We acknowledge the logistic support from the Iwokrama International Centre for Rain Forest Conservation and Development, Guyana, the Karanambu Trust, Guyana, the National Park Administration, Argentina, and the Congo Basin Institute, Cameroon. Funding for this study was provided by research grants from The Swedish Research Council (VR: grant no. 2016-03356), the Swedish Foundation for International Cooperation in Research and Higher Education, and Erik Philip Sörensens Stiftelse to E.I.S and from The Royal Physiographic Society of Lund, the Jörgen Lindströms Fund and “Lunds Djurskyddsfond” to B.W. We thank Stephen De Lisle and Miguel Gómez-Llano for helpful comments on the manuscript and Brian P. Hanotte for computational support.

## Literature Cited

Abbott J.C., Bota-Sierra C.A., Guralnick R., Kalkman V., González-Soriano E., Novelo-Gutiérrez R., Bybee S., Ware J., Belitz M.W. 2022. Diversity of nearctic dragonflies and damselflies (Odonata). Diversity. 14:575.

Anderson R.C. 2009. Do dragonflies migrate across the western Indian Ocean? J. Trop. Ecology. 25:347–358.

Antonelli A., Zizka A., Silvestro D., Scharn R., Cascales-Miñana B., Bacon C.D. 2015. An engine for global plant diversity: Highest evolutionary turnover and emigration in the American tropics. Front. Genet. 6:130.

Barnosky A.D. 2005. Effects of Quaternary climatic change on speciation in mammals. J. Mamm. Evol. 12:247–264.

Beaulieu J.M., O’Meara B.C. 2016. Detecting hidden diversification shifts in models of trait-dependent speciation and extinction. Syst. Biol. 65:583–601.

Belmaker J., Jetz W. 2015. Relative roles of ecological and energetic constraints, diversification rates and region history on global species richness gradients. Ecol. Lett. 18:563–571.

Botero C.A., Dor R., McCain C.M., Safran R.J. 2014. Environmental harshness is positively correlated with intraspecific divergence in mammals and birds. Mol. Ecol. 23:259–268.

Boudot J.-P., Kalkman V.J.(eds). 2015. Atlas of the European dragonflies and damselflies. The Netherlands: KNNV Publishing.

Brock C.D., Harmon L.J., Alfaro M.E. 2011. Testing for temporal variation in diversification rates when sampling is incomplete and nonrandom. Syst. Biol. 60:410–419.

Brown J.H. 2014. Why are there so many species in the tropics? J. Biogeogr. 41:8–22.

Budd G.E., Mann R.P. 2018. History is written by the victors: The effect of the push of the past on the fossil record. Evolution. 72:2276–2291.

Bybee S.M., Kalkman V.J., Erickson R.J., Frandsen P.B., Breinholt J.W., Suvorov A., Dijkstra K.-D.B., Cordero-Rivera A., Skevington J.H., Abbott J.C., Sánchez-Herrera M., Lemmon A.L., Lemmon E.M., Ware J.L. 2021. Phylogeny and classification of Odonata using targeted genomics. Mol. Phylogenet. Evol. 160:107115.

Carle F.L., Kjer K., May M. 2008. Evolution of Odonata, with special reference to Coenagrionoidea (Zygoptera). Arthropod Syst. 66:37–44.

Corbet P.S. 1999. Dragonflies: Behavior and ecology of Odonata. Colchester, UK: Harley.

Couvreur T.L.P., Forest F., Baker W.J. 2011. Origin and global diversification patterns of tropical rain forests: Inferences from a complete genus-level phylogeny of palms. BMC Biol. 9:44.

Cusimano N., Renner S.S. 2010. Slowdowns in diversification rates from real phylogenies may not be real. Syst. Biol. 59:458–464.

Cutter A.D., Gray J.C. 2016. Ephemeral ecological speciation and the latitudinal biodiversity gradient. Evolution. 70:2171–2185.

Czekanski-Moir J.E., Rundell R.J. 2019. The ecology of nonecological speciation and nonadaptive radiations. Trends Ecol. Evol. 34:400–415.

Darwin C.R. 1859. The origin of species by means of natural selection, or the preservation of favoured races in the struggle for life. London, UK: John Murray.

De Baets K., Antonelli A., Donoghue P.C. 2016. Tectonic blocks and molecular clocks. Philos. Trans. R. Soc. Lond., B. 371:20160098.

DeConto R.M., Pollard D., Wilson P.A., Pälike H., Lear C.H., Pagani M. 2008. Thresholds for Cenozoic bipolar glaciation. Nature. 455:652–657.

Dijkstra K.-D.B. 2017. African Dragonflies and Damselflies. Online: http://addo.adu.org.za/

Dijkstra K.D.B., Kalkman V.J., Dow R.A., Stokvis F.R., Van Tol J.A.N. 2014. Redefining the damselfly families: A comprehensive molecular phylogeny of Zygoptera (Odonata). Syst. Entomol. 39:68–96.

Dobzhansky T. 1950. Evolution in the tropics. Amer. Sci. 38:209–221.

Dynesius M., Jansson R. 2000. Evolutionary consequences of changes in species’ geographical distributions driven by Milankovitch climate oscillations. Proc. Natl. Acad. Sci. U.S.A. 97:9115–9120.

Economo E.P., Narula N., Friedman N.R., Weiser M.D., Guénard B. 2018. Macroecology and macroevolution of the latitudinal diversity gradient in ants. Nat. Commun. 9:1778.

Etienne R.S., Cabral J.S., Hagen O., Hartig F., Hurlbert A.H., Pellissier L., Pontarp M., Storch D. 2019. A minimal model for the latitudinal diversity gradient suggests a dominant role for ecological limits. Am. Nat. 194:E122–E133.

Etienne R.S., Haegeman B. 2012. A conceptual and statistical framework for adaptive radiations with a key role for diversity dependence. Am. Nat. 180:E75–E89.

Etienne R.S., Haegeman B., Stadler T., Aze T., Pearson P.N., Purvis A., Phillimore A.B. 2011. Diversity-dependence brings molecular phylogenies closer to agreement with the fossil record. Proc. R. Soc. B. 279:1300–1309.

Fischer A.G. 1960. Latitudinal variations in organic diversity. Evolution. 14:64–81.

Gaboriau T., Albouy C., Descombes P., Mouillot D., Pellissier L., Leprieur F. 2019. Ecological constraints coupled with deep-time habitat dynamics predict the latitudinal diversity gradient in reef fishes. Proc. R. Soc. B. 286:20191506.

Goldberg E.E., Lancaster L.T., Ree R.H. 2011. Phylogenetic inference of reciprocal effects between geographic range evolution and diversification. Syst. Biol. 60:451–465.

Gomulkiewicz R., Holt R.D. 1995. When does evolution by natural selection prevent extinction? Evolution. 49:201–207.

Grewe Y., Hof C., Dehling D.M., Brandl R., Brändle M. 2013. Recent range shifts of European dragonflies provide support for an inverse relationship between habitat predictability and dispersal. Global Ecol. Biogeogr. 22:403–409.

Haldane J.B.S. 1937. The effect of variation of fitness. Am. Nat. 71:337–349.

Hanly P.J., Mittelbach G.G., Schemske D.W. 2017. Speciation and the latitudinal diversity gradient: Insights from the global distribution of endemic fish. Am. Nat. 189:604–615.

Harvey M.G., Bravo G.A., Claramunt S., Cuervo A.M., Derryberry G.E., Battilana J., Seeholzer G.F., McKay J.S., O’Meara B.C., Faircloth B.C., Edwards S.V., Pérez-Emán J., Moyle R.G., Sheldon F.H., Aleixo A., Smith B.T., Chesser R.T., Silveira L.F., Cracraft J., Brumfield R.T., Derryberry E.P. 2020. The evolution of a tropical biodiversity hotspot. Science. 370:1343–1348.

Hawkins B.A., Diniz-Filho J.A.F., Jaramillo C.A., Soeller S.A. 2007. Climate, niche conservatism, and the global bird diversity gradient. Am. Nat. 170:S16–S27.

Heath T.A., Huelsenbeck J.P., Stadler T. 2014. The fossilized birth–death process for coherent calibration of divergence-time estimates. Proc. Natl. Acad. Sci. U.S.A. 111:E2957– E2966.

Herrera-Alsina L., Pigot A.L., Hildenbrandt H., Etienne R.S. 2018. The influence of ecological and geographic limits on the evolution of species distributions and diversity. Evolution. 72:1978–1991.

Hillebrand H. 2004. On the generality of the latitudinal diversity gradient. Am. Nat. 163:192– 211.

Ho S.Y.W., Tong K.J., Foster C.S.P., Ritchie A.M., Lo N., Crisp M.D. 2015. Biogeographic calibrations for the molecular clock. Biol. Lett. 11:20150194.

Höhna S. 2015. The time-dependent reconstructed evolutionary process with a key-role for mass-extinction events. J. Theor. Biol. 380:321–331.

Höhna S., Heath T.A., Boussau B., Landis M.J., Ronquist F., Huelsenbeck J.P. 2014. Probabilistic graphical model representation in phylogenetics. Syst. Biol. 63:753–771.

Höhna S., Landis M.J., Heath T.A., Boussau B., Lartillot N., Moore B.R., Huelsenbeck J.P., Ronquist F. 2016. RevBayes: Bayesian phylogenetic inference using graphical models and an interactive model-specification language. Syst. Biol. 65:726–736.

Jablonski D., Huang S., Roy K., Valentine J.W. 2017. Shaping the latitudinal diversity gradient: New perspectives from a synthesis of paleobiology and biogeography. Am. Nat. 189:1– 12.

Jablonski D., Roy K., Valentine J.W. 2006. Out of the tropics: Evolutionary dynamics of the latitudinal diversity gradient. Science. 314:102–106.

Jansson R., Dynesius M. 2002. The fate of clades in a world of recurrent climatic change: Milankovitch oscillations and evolution. Annu. Rev. Ecol. Evol. Syst. 33:741–777.

Jocque M., Field R., Brendonck L., De Meester L. 2010. Climatic control of dispersal–ecological specialization trade-offs: A metacommunity process at the heart of the latitudinal diversity gradient? Global Ecol. Biogeogr. 19:244–252.

Jolly D., Harrison S.P., Damnati B., Bonnefille R. 1998. Simulated climate and biomes of Africa during the Late Quaternary: Comparison with pollen and lake status data. Quaternary Science Reviews. 17:629–657.

Jones H.B.C., Lim K.S., Bell J.R., Hill J.K., Chapman J.W. 2016. Quantifying interspecific variation in dispersal ability of noctuid moths using an advanced tethered flight technique. Ecol. Evol. 6:181–190.

Jønsson K.A., Tøttrup A.P., Borregaard M.K., Keith S.A., Rahbek C., Thorup K. 2016. Tracking animal dispersal: From individual movement to community assembly and global range dynamics. Trends in Ecology & Evolution. 31:204–214.

Kidwell S.M., Holland S.M. 2002. The quality of the fossil record: Implications for evolutionary analyses. Annu. Rev. Ecol. Evol. Syst. 33:561–588.

Kinlock N.L., Prowant L., Herstoff E.M., Foley C.M., Akin-Fajiye M., Bender N., Umarani M., Ryu H.Y., Şen B., Gurevitch J. 2018. Explaining global variation in the latitudinal diversity gradient: Meta-analysis confirms known patterns and uncovers new ones. Global Ecol. Biogeogr. 27:125–141.

Kodandaramaiah U. 2011. Tectonic calibrations in molecular dating. Curr. Zool. 57:116–124.

Kohli M., Letsch H., Greve C., Béthoux O., Deregnaucourt I., Liu S., Zhou X., Donath A., Mayer C., Podsiadlowski L., Gunkel S., Machida R., Niehuis O., Rust, J., Wappler T., Yu X., Misof B., Ware J.L. 2021. Evolutionary history and divergence times of Odonata (dragonflies and damselflies) revealed through transcriptomics. Iscience. 24:103324.

Kubatko L.S., Degnan J.H. 2007. Inconsistency of phylogenetic estimates from concatenated data under coalescence. Syst. Biol. 56:17–24.

Lancaster L.T., Dudaniec R.Y., Hansson B., Svensson E.I. 2015. Latitudinal shift in thermal niche breadth results from thermal release during a climate-mediated range expansion. J. Biogeogr. 42:1953–1963.

Landis M., Edwards E.J., Donoghue M.J. 2021. Modeling phylogenetic biome shifts on a planet with a past. Syst. Biol. 70:86–107.

Landis M.J. 2017. Biogeographic dating of speciation times using paleogeographically informed processes. Syst. Biol. 66:128–144.

Landis M.J., Freyman W.A., Baldwin B.G. 2018. Retracing the Hawaiian silversword radiation despite phylogenetic, biogeographic, and paleogeographic uncertainty. Evolution. 72:2343–2359.

Loarie S.R., Duffy P.B., Hamilton H., Asner G.P., Field C.B., Ackerly D.D. 2009. The velocity of climate change. Nature. 462:1052–1055.

MacArthur R.H. 1969. Patterns of communities in the tropics. Biol. J. Linn. Soc. 1:19–30.

Magee A.F., Höhna S., Vasylyeva T.I., Leaché A.D., Minin V.N. 2020. Locally adaptive Bayesian birth-death model successfully detects slow and rapid rate shifts. PLoS Comput. Biol. 16:e1007999.

May M.L. 2019. Aquatic insects: Behavior and ecology. Aquatic Insects: Behavior and Ecology. Springer. p. 35–73.

McPeek M.A., Brown J.M. 2000. Building a regional species pool: Diversification of the *Enallagma* damselflies in eastern North America. Ecology. 81:904–920.

McPeek M.A., Gavrilets S. 2006. The evolution of female mating preferences: Differentiation from species with promiscuous males can promote speciation. Evolution. 60:1967–1980.

Meseguer A.L., Condamine F.L. 2020. Ancient tropical extinctions at high latitudes contributed to the latitudinal diversity gradient. Evolution. 74:1966–1987.

Milá B., McCormack J.E., Castañeda G., Wayne R.K., Smith T.B. 2007. Recent postglacial range expansion drives the rapid diversification of a songbird lineage in the genus *Junco*. Proc. R. Soc. B. 274:2653–2660.

Miller E.C., Román-Palacios C. 2021. Evolutionary time best explains the latitudinal diversity gradient of living freshwater fish diversity. Global Ecol. Biogeogr. 30:749–763.

Misof B., Liu S., Meusemann K., Peters R.S., Donath A., Mayer C., Frandsen P.B., Ware J., Flouri T., Beutel R.G., Niehuis O., Petersen M., Izquierdo-Carrasco F., Wappler T., Rust J., Aberer A.J., Aspöck U., Aspöck H., Bartel D., Blanke A., Berger S., Böhm A., Buckley T.R., Calcott B., Chen J., Friedrich F., Fukui M., Fujita M., Greve C., Grobe P., Gu S., Huang Y., Jermiin L.S., Kawahara A.Y., Krogmann L., Kubiak M., Lanfear R., Letsch H., Li Y., Li Z., Li J., Lu H., Machida R., Mashimo Y., Kapli P., McKenna D.D., Meng G., Nakagaki Y., Navarrete-Heredia J.L., Ott M., Ou Y., Pass G., Podsiadlowski L., Pohl H., Reumont von B.M., Schütte K., Sekiya K., Shimizu S., Slipinski A., Stamatakis A., Song W., Su X., Szucsich N.U., Tan M., Tan X., Tang M., Tang J., Timelthaler G., Tomizuka S., Trautwein M., Tong X., Uchifune T., Walzl M.G., Wiegmann B.M., Wilbrandt J., Wipfler B., Wong T.K.F., Wu Q., Wu G., Xie Y., Yang S., Yang Q., Yeates D.K., Yoshizawa K., Zhang Q., Zhang R., Zhang W., Zhang Y., Zhao J., Zhou C., Zhou L., Ziesmann T., Zou S., Li Y., Xu X., Zhang Y., Yang H., Wang J., Wang J., Kjer K.M., Zhou X. 2014. Phylogenomics resolves the timing and pattern of insect evolution. Science. 346:763–767.

Mittelbach G.G., Schemske D.W., Cornell H.V., Allen A.P., Brown J.M., Bush M.B., Harrison S.P., Hurlbert A.H., Knowlton N., Lessios H.A. McCain C., McCune A.R., McDade L.A., McPeek M.A., Near T.J., Price T.D., Ricklefs R.E., Roy K. Sax D.F., Schluter D., Sobel J.M., Turelli M. 2007. Evolution and the latitudinal diversity gradient: Speciation, extinction and biogeography. Ecol. Lett. 10:315–331.

Moen D., Morlon H. 2014. Why does diversification slow down? Trends Ecol. Evol. 29:190– 197.

Morales-Barbero J., Martinez P.A., Ferrer-Castán D., Olalla-Tárraga M.Á. 2018. Quaternary refugia are associated with higher speciation rates in mammalian faunas of the Western Palaearctic. Ecography. 41:607–621.

Moritz C., Patton J.L., Schneider C.J., Smith T.B. 2000. Diversification of rainforest faunas: An integrated molecular approach. Annu. Rev. Ecol. Evol. Syst. 31:533–563.

Morley R.J. 2007. Tropical rainforest responses to climatic change. In: Bush M., Flenley J., Gosling W., editors. Tropical rainforest responses to climatic change. Berlin, Germany: Springer. p. 1–31.

Morlon H., Potts M.D., Plotkin J.B. 2010. Inferring the dynamics of diversification: A coalescent approach. PLoS Biol. 8:e1000493.

Otto-Bliesner B.L., Brady E.C., Tomas R.A., Albani S., Bartlein P.J., Mahowald N.M., Shafer S.L., Kluzek E., Lawrence P.J., Leguy G., others. 2020. A comparison of the CMIP6 midHolocene and lig127k simulations in CESM2. Paleoceanogr. Paleoclimatol. 35:e2020PA003957.

Owens H.L., Lewis D.S., Dupuis J.R., Clamens A.L., Sperling F.A., Kawahara A.Y., Guralnick R.P., Condamine F.L. 2017. Latitudinal diversity gradient in New World swallowtail butterflies is caused by contrasting patterns of out-of- and into-the-tropics dispersal. Global Ecol. Biogeogr.

Paulson D. 2009. Dragonflies and Damselflies of the West. Princeton University Press.

Paulson D. 2011. Dragonflies and Damselflies of the East. Princeton University Press.

Paulson D.R. 2001. Recent Odonata records from southern Florida-effects of global warming? Int. J. Odonatol. 4:57–69.

Pianka E.R. 1966. Latitudinal gradients in species diversity: A review of concepts. Am. Nat. 100:33–46.

Polato N.R., Gill B.A., Shah A.A., Gray M.M., Casner K.L., Barthelet A., Messer P.W., Simmons M.P., Guayasamin J.M., Encalada A.C., Kondratieff B.C., Flecker A.S., Thomas S.A., Ghalambor C.K., Poff L., Funk W.C., Zamudio K. 2018. Narrow thermal tolerance and low dispersal drive higher speciation in tropical mountains. Proc. Natl. Acad. Sci. U.S.A.. 115:12471–12476.

Pyron R.A. 2014. Temperate extinction in squamate reptiles and the roots of latitudinal diversity gradients. Global Ecol. Biogeogr. 23:1126–1134.

Pyron R.A., Wiens J.J. 2013. Large-scale phylogenetic analyses reveal the causes of high tropical amphibian diversity. Proc. R. Soc. B. 280:20131622.

R Core Team. 2021. R: A language and environment for statistical computing. Vienna, Austria: R Foundation for Statistical Computing.

Rabosky D.L. 2009. Ecological limits and diversification rate: Alternative paradigms to explain the variation in species richness among clades and regions. Ecol. Lett. 12:735–743.

Rabosky D.L., Chang J., Title P.O., Cowman P.F., Sallan L., Friedman M., Kaschner K., Garilao C., Near T.J., Coll M., Alfaro M.E. 2018. An inverse latitudinal gradient in speciation rate for marine fishes. Nature. 559:392–395.

Rangel T.F., Edwards N.R., Holden P.B., Diniz-Filho J.A.F., Gosling W.D., Coelho M.T.P., Cassemiro F.A.S., Rahbek C., Colwell R.K. 2018. Modeling the ecology and evolution of biodiversity: Biogeographical cradles, museums, and graves. Science. 361:eaar5452.

Ree R.H., Sanmartín I. 2009. Prospects and challenges for parametric models in historical biogeographical inference. J. Biogeogr. 36:1211–1220.

Revell L.J., Harmon L.J., Glor R.E. 2005. Under-parameterized model of sequence evolution leads to bias in the estimation of diversification rates from molecular phylogenies. Syst. Biol. 54:973–983.

Rolland J., Condamine F.L., Beeravolu C.R., Jiguet F., Morlon H. 2015. Dispersal is a major driver of the latitudinal diversity gradient of Carnivora. Global Ecol. Biogeogr. 24:1059– 1071.

Rolland J., Condamine F.L., Jiguet F., Morlon H. 2014. Faster speciation and reduced extinction in the tropics contribute to the mammalian latitudinal diversity gradient. PLoS Biol. 12:e1001775.

Ronquist F., Klopfstein S., Vilhelmsen L., Schulmeister S., Murray D.L., Rasnitsyn A.P. 2012. A total-evidence approach to dating with fossils, applied to the early radiation of the Hymenoptera. Syst. Biol. 61:973–999.

Rosenzweig M.L., others. 1995. Species diversity in space and time. Cambridge University Press.

Roy K., Goldberg E.E. 2007. Origination, extinction, and dispersal: Integrative models for understanding present-day diversity gradients. Am. Nat. 170:S71–S85.

Rundell R.J., Price T.D. 2009. Adaptive radiation, nonadaptive radiation, ecological speciation and nonecological speciation. Trends Ecol. Evol.. 24:394–399.

Salisbury C.L., Seddon N., Cooney C.R., Tobias J.A. 2012. The latitudinal gradient in dispersal constraints: Ecological specialisation drives diversification in tropical birds. Annu. Rev. Ecol. Evol. Syst. 15:847–855.

Salzmann U., Haywood A.M., Lunt D.J., Valdes P.J., Hill D.J. 2008. A new global biome reconstruction and data-model comparison for the Middle Pliocene. Global Ecol. Biogeogr. 17:432–447.

Saupe E.E., Myers C.E., Peterson A.T., Soberón J., Singarayer J., Valdes P., Qiao H. 2019. Spatio-temporal climate change contributes to latitudinal diversity gradients. Nat. Ecol. Evol. 3:1419–1429.

Sánchez-Ramírez S., Tulloss R.E., Guzmán-Dávalos L., Cifuentes-Blanco J., Valenzuela R., Estrada-Torres A., Ruán-Soto F., Díaz-Moreno R., Hernández-Rico N., Torres-Gómez M., León H., Moncalvo J.-M. 2015. In and out of refugia: Historical patterns of diversity and demography in the North American Caesar’s mushroom species complex. Mol. Ecol. 24:5938–5956.

Schemske D.W., Mittelbach G.G., Cornell H.V., Sobel J.M., Roy K. 2009. Is there a latitudinal gradient in the importance of biotic interactions? Annu. Rev. Ecol. Evol. Syst. 40:245– 269.

Schluter D. 2016. Speciation, ecological opportunity, and latitude: (American Society of Naturalists Address). Am. Nat. 187:1–18.

Siepielski A.M., Hasik A.Z., Ousterhout B.H. 2018. An ecological and evolutionary perspective on species coexistence under global change. Curr. Opin. Insect Sci. 29:71–77.

Siepielski A.M., Hung K.-L., Bein E.E.B., McPeek M.A. 2010. Experimental evidence for neutral community dynamics governing an insect assemblage. Ecology. 91:847–857.

Smith B.T., Bryson Jr R.W., Houston D.D., Klicka J. 2012. An asymmetry in niche conservatism contributes to the latitudinal species diversity gradient in New World vertebrates. Ecol. Lett. 15:1318–1325.

Spano C.A., Hernández C.E., Rivadeneira M.M. 2016. Evolutionary dispersal drives the latitudinal diversity gradient of stony corals. Ecography. 39:836–843.

Stadler T. 2011. Mammalian phylogeny reveals recent diversification rate shifts. Proc. Natl. Acad. Sci. U.S.A. 108:6187–6192.

Stephens P.R., Wiens J.J. 2003. Explaining species richness from continents to communities: The time-for-speciation effect in emydid turtles. Am. Nat. 161:112–128.

Stevens R.D. 2006. Historical processes enhance patterns of diversity along latitudinal gradients. Proc. R. Soc. B. 273:2283–2289.

Suvorov A., Scornavacca C., Fujimoto M.S., Bodily P., Clement M., Crandall K.A., Whiting M.F., Schrider D.R., Bybee S.M. 2021. Deep ancestral introgression shapes evolutionary history of dragonflies and damselflies. Syst. Biol.:syab063.

Svensson E.I. 2012. Non-ecological speciation, niche conservatism and thermal adaptation: How are they connected? Org. Divers. Evol. 12:229–240.

Svensson E.I., Eroukhmanoff F., Friberg M. 2006. Effects of natural and sexual selection on adaptive population divergence and premating isolation in a damselfly. Evolution. 60:1242–1253.

Svensson E.I., Gómez-Llano M.A., Torres A.R., Bensch H.M. 2018. Frequency dependence and ecological drift shape coexistence of species with similar niches. Am. Nat. 191:691–703.

Tiley G.P., Poelstra J.W., Dos Reis M., Yang Z., Yoder A.D. 2020. Molecular clocks without rocks: New solutions for old problems. Trends Genet.

Toussaint E.F.A., Bybee S.M., Erickson R.J., Condamine F.L. 2019. Forest giants on different evolutionary branches: Ecomorphological convergence in helicopter damselflies. Evolution. 73:1045–1054.

Troast D., Suhling F., Jinguji H., Sahlén G., Ware J. 2016. A global population genetic study of *Pantala flavescens*. PLoS One. 11:e0148949.

Wallace A.R. 1876. The geographical distribution of animals. Macmillan; Company.

Wallace A.R. 1878. Tropical nature, and other essays. Macmillan; Company.

Waller J.T., Svensson E.I. 2017. Body size evolution in an old insect order: No evidence for Cope’s Rule in spite of fitness benefits of large size. Evolution. 71:2178–2193.

Weir J.T., Schluter D. 2004. Ice sheets promote speciation in boreal birds. Proc. R. Soc. B 271:1881–1887.

Weir J.T., Schluter D. 2007. The latitudinal gradient in recent speciation and extinction rates of birds and mammals. Science. 315:1574–1576.

Wickham H. 2016. ggplot2: Elegant graphics for data analysis. Springer-Verlag New York.

Wiens J.J. 2007. Global patterns of diversification and species richness in amphibians. Am. Nat. 170:S86–S106.

Wiens J.J. 2011. The causes of species richness patterns across space, time, and clades and the role of “ecological limits.” Q. Rev. Biol. 86:75–96.

Wikelski M., Moskowitz D., Adelman J.S., Cochran J., Wilcove D.S., May M.L. 2006. Simple rules guide dragonfly migration. Biol. Lett.. 2:325–329.

Wills M.A. 2001. How good is the fossil record of arthropods? An assessment using the stratigraphic congruence of cladograms. Geol. J. 36:187–210.

Wright S., Keeling J., Gillman L. 2006. The road from Santa Rosalia: A faster tempo of evolution in tropical climates. Proc. Natl. Acad. Sci. U.S.A. 103:7718–7722.

Yu G., Smith D., Zhu H., Guan Y., Lam T.T.-Y. 2017. ggtree: An R package for visualization and annotation of phylogenetic trees with their covariates and other associated data. Methods Ecol. Evol. 1:28–36.

Zheng D., Wang B., Chang S.-C. 2017. *Palaeodisparoneura cretacica* sp. Nov., A new damselfly (Odonata: Zygoptera: Platycnemididae) from mid-Cretaceous Burmese amber. Comptes Rendus Palevol. 16:235–240.

