## Supporting Material for "Tropical Origin, Global Diversification and Dispersal in the Pond Damselflies (Coenagrionoidea) Revealed by a New Molecular Phylogeny"

2023-06-08

[1] Department of Zoology, Stockholm University, Stockholm 106-91, Sweden

[2] School of Biology, University of Costa Rica, San José 11501-2060, Costa Rica

[3] Department of Biological Sciences, Rutgers, The State University of New Jersey, Newark, NJ, U.S.A.

[4] Department of Biology, Evolutionary Ecology Unit, Ecology Building, Lund University, Lund 223-62, Sweden

**Keywords:** biogeography, extinction, graphical models, latitudinal diversity gradient, paleogeography, phylogeny

**Literature cited** **77**

### Extended Methods

#### Sequence data

We sampled 669 taxa in two of the families included in Coenagrionoidea (Table S1): pond damselflies (family Coenagrionidae) and featherlegs (family Platycnemididae). The identity of the sampled taxa, their identification codes and specimen housing locations are shown in Table S2. Genomic DNA was extracted from 1-2 legs using a Qiagen DNeasy tissue kit and following the manufacturer's instructions. To augment DNA yield, we increased the amount of proteinase K and incubation time for samples which have been collected before 2005 and thereafter maintained at room temperature in museum collections.

We amplified and sequenced five molecular markers: a 481 bp fragment of the cytochrome oxidase subunit I (COI; mitochondrial), the third domain of the 16S ribosomal DNA (16S; mitochondrial), a 328 bp coding region of the histone 3 gene (H3; nuclear), a fragment including coding (336 bp) and intronic regions of the arginine phenyl-methyltransferase gene (PMTR; nuclear) and the seventh hypervariable (divergent) region of the 28S ribosomal DNA (D7; nuclear). Four of these markers (COI, 16S, H3 and D7) are of common use for phylogenetic studies in dragonflies and damselflies (e.g. Ware et al. 2007; Bybee et al. 2008; Dijkstra et al. 2014), whereas PMTR has been recently identified as an informative region for phylogeographic studies in Coenagrionidae (Ferreira et al. 2014). Primers sequences were obtained from previous studies (Xiong and Kocher 1991; Kambhampati 1995; Colgan et al. 1998; Kjer et al. 2001; Ware et al. 2007; Ferreira et al. 2014), and can be found, along with Polymerase Chain Reaction programs, in Willink et al. (2019, Table A5). Amplification products were purified using the Affymetrix ExoSAP-IT reagent (Cat. No. 78200) or by ethanol precipitation. Purified products were sequenced on an ABI 3100 capillary sequencer at Lund University, or by Macrogen (Macrogen USA, Rockville, MD).

**Table S1.** Proportion of taxa sampled for phylogenetic analyses in this study. The taxa are currently classified into the two main clades composing the Coenagrionoidea superfamily, the pond damselflies (Coenagrionidae) and the featherlegs (Platycnemididae). Sampling fractions are based on the World Odonata List (Paulson and Schorr 2021).

| Family | No.species | Percent.species | No.genera | Percent.genera |
|---|---|---|---|---|
| Coenagrionidae | 556 | ~ 41% | 84 | ~ 69% |
| Platycnemididae | 113 | ~ 25% | 31 | ~ 72% |
| total | 669 | ~ 37% | 115 | ~ 70% |

To maximize the taxonomic sampling in our analysis we combined our sequence data (n = 1695 sequences, described above) with data publicly available on NCBI (n = 564). We also obtained unpublished sequence data (n = 105) for specimens of 33 taxa, which are part of the Odonata collection at the Naturalis Biodiversity Center, The Netherlands. These sequence data were obtained following a similar protocol as ours, which is described in Dijkstra et al. (2014). We preferentially used our own sequences and included NCBI and Naturalis data only when 1) taxa with sequence data already available were not represented in our species sampling (n = 205 taxa) and 2) when we failed to amplify a particular marker in our sample but sequence data for that marker and taxon was already available (n = 27 taxa).

When we had multiple samples for the same taxon we selected the sample with highest coverage for our five markers. If two or more samples of a taxon had equally high coverage we chose a single sample, preferentially from a specimen we collected ourselves or received as a permanent loan, rather than samples from museum collections. In the few cases where amplification and/or sequencing of at least one marker failed for the sample with highest coverage but sequence data from another individual was available (n = 14 taxa), we combined data from these different samples, aiming to maximize coverage in the concatenated alignment.

**Table S2. The full table is attached as a tab-delimited file.** Molecular sequence data used in this study. Sequence data was obtained for up to five loci (16S, COI, D7, H3, PMTR, see Extended Methods). Specimens sequenced in this study are identified with the letters "BEA." Accession numbers are given for sequence data downloaded from NCBI (GenBank). Sequence data provided by KD Dijkstra at the Naturalis Biodiversity Center are identified with the letters "RMNH.INS." and "DIJ." Missing data is coded as "–."

| Taxon | r16S | COI | H3 | rD7 | PMTR | Collection | Identification |
|---|---|---|---|---|---|---|---|
| *Acanthagrion adustum* | BEA019 | BEA019 | BEA019 | BEA019 | BEA019 | LU | NVE |
| *Acanthagrion amazonicum* | BEA351 | – | – | – | – | FSCA | – |
| *Acanthagrion apicale* | BEA044 | BEA044 | BEA044 | BEA044 | BEA044 | LU | NVE |
| *Acanthagrion abunae* | BEA388 | BEA388 | BEA388 | BEA388 | BEA388 | RU | MSH |
| ... | ... | ... | ... | ... | ... | ... | ... |

The housing facility of the specimens sequenced for this study is given for specimens held in academic institutions: National Institute of Advanced Industrial Science & Technology, Japan (AIST); Brigham Young University, USA (BYU); Florida State Collection of Arthropods, USA (FSCA); Laboratoire de Biométrie et Biologie Évolutive, Université de Lyon, France (LBBE); Svensson Lab, Lund University, Sweden (LU); Meier Lab, National University of Singapore, Singapore (NUS); New Zealand Arthropod Collection, New Zealand (NZAC), Naturalis Biodiversity Center, The Netherlands (RMNH); Ware Lab, Rutgers University, USA (RU); Smithsonian Institute, USA (SI); United States Department of Agriculture, USA (USDA). Identification of specimens not obtained from curated museum collections (FSCA and SI) was performed by us (BW and EIS) or provided by expert entomologists: Jackie Brown (JB), Seth Bybee (SB), Sylvain Charlat (SC), Klaas D. Dijkstra (KDD), Sonia Ferréira (SF), Ryo Futahashi (RF), Kawsar Khan (KK), Oleg Kosterin (OK), Rudolf Meier (RM), Antónia Monteiro (AM), Julio Neto (JN), Viktor Nilsson-Örtman (VNO), Samuel Renner (SR), Melissa Sánchez-Herrera (MSH), Issah Seidu (IS) and Nathalia von Ellenrieder (NVE).

#### Sequence alignment and inclusion criteria

We aligned COI, H3 and ribosome coding sequences using MUSCLE v. 3.8.31 (Edgar 2004) under default settings. There were no indels or stop codons in either of the two protein-coding sequences. Also, we found no evidence of substantial third-codon saturation using DAMBE (Xia and Xie 2001; Xia and Lemey 2009), or by visual inspection of plots of model-corrected vs. uncorrected genetic distances (Fig. S1). The ribosomal DNA sequences (D7 and 16S), initially aligned with MUSCLE, were subsequently matched to the secondary structure of the ribosomal molecules to manually verify or correct the alignment of all hydrogen-bonded regions (Kjer 1995). Hereafter, we excluded all sites within these single-stranded regions that were ambiguously aligned due to multiple insertions and deletions and over-representation of A and T nucleotides.

Our last marker (PMTR) included a combination of exonic and intronic regions. Because intronic regions are variable in length we aligned these sequences using PRANK (Löytynoja and Goldman 2005, 2008), in order to distinguish insertions and deletions in our alignment and to avoid overmatching gapped sites. However, PRANK alignments can be problematic for datasets combining conserved regions and regions of low complexity, which may have a higher rate of identical independent deletions. In this case, the conserved regions are properly aligned but variable regions are placed as non-homologous insertions, producing an extremely gapped alignment and vastly reducing the number the phylogenetic informative sites. To lessen this problem in our intronic data we enforced a branch length multiplier of two, which produced more mismatches in the alignment. This way we allowed for a contribution of the intronic regions to phylogenetic inference while still generating a more conservative alignment than a phylogenetically-naïve algorithm would. As the alignment was to be used to infer the tree topology, we did not provide a guide tree and let the first guide tree to be inferred from the input data. Likewise, we omitted the option +F that makes inferred insertions permanent and therefore is more sensitive to incorrect topologies in the guide tree. PRANK was set to iterate the phylogentically-aware alignment 20 times. This means a new guide tree was inferred after each alignment and the highest scoring alignment was selected by randomly breaking score ties at each iteration.

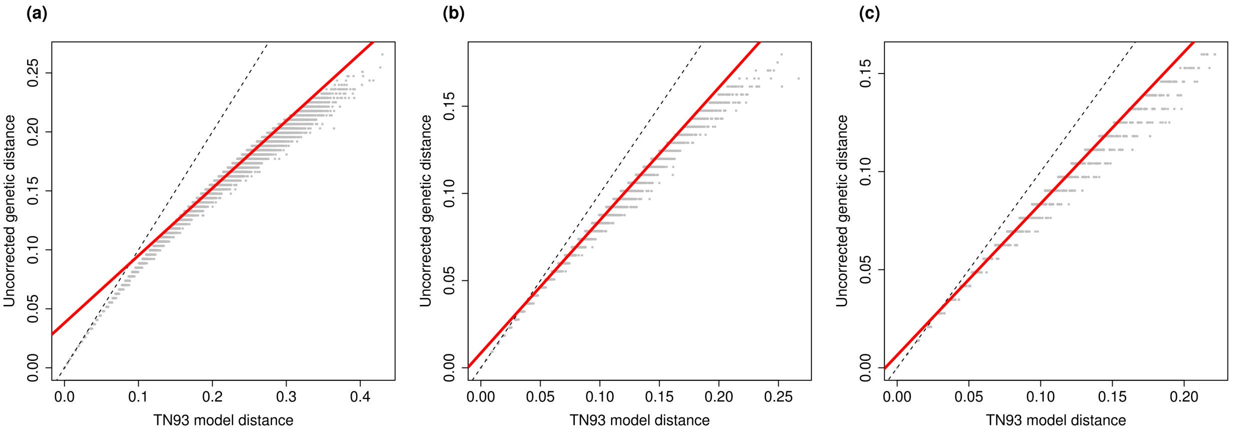

**Figure S1.** Exploratory visualization of substitution saturation in three protein-coding markers used for phylogenetic inference in this study: a) Cytochrome oxidase subunit I (COI), b) Histone 3 (H3) and c) Phenyl-methyl transferase (PMTR). A decay in uncorrected genetic distances with increasing model-corrected genetic distances would be suggestive of DNA saturation due to numerous substitution events, which may reduce the phylogenetic information in the sequence alignments. Corrected genetic distances are based on a TN93 model of molecular evolution, which parametrises three types of substitution rates, transversions and both types of transitions. Corrected distances were estimated using the *ape* package (Paradis and Schliep 2019) in R. The dashed line shows an identity relationship between corrected and uncorrected distances and the red lines represents a linear fit. In addition to not observing a strong deviation from linearity in the relationship, entropy-based analyses in DAMBE (Xia and Xie 2001; Xia and Lemey 2009) indicated no evidence of substantial saturation in these sequences, assuming invariant proportions of 0.05, 0.1, 0.20 and 0.4, except for extremely asymmetrical trees, which are generally unlikely, and high invariant proportions (0.4) in COI and PMTR.

#### Topology inference

To select an appropriate data-partition scheme for phylogenetic inference we estimated the marginal likelihoods of six alternative partition schemes using the stepping stone algorithm (Xie et al. 2010) (Table S3). We started with a model with only gene partitions (M1) and compared it with a model in which the first and second codon positions of protein-coding loci shared model parameters that differed from the third codon position (M2). We then compared this partition scheme against models in which all codon positions for one or more loci evolved under different parameters (M3-M6). We selected the least parametrised model not outperformed by a more parametrised alternative, by considering a difference in marginal likelihood $> 2$ as sufficient support for the more parametrised alternative.

**Table S3.** Alternative partition schemes compared using marginal likelihoods approximations in RevBayes (Höhna et al. 2016). Each colour represents a data partition with its own substitution-rate parameters. Marginal likelihoods (ML) were approximated using the stepping stone algorithm (Xie et al. 2010) for each partition scheme. In all partition schemes a GTR + $\Gamma$ model was assumed for all markers. The model with highest marginal likelihood is highlighted in **bold**.

| Model | ML | COI.p1 | COI.p2 | COI.p3 | H3.p1 | H3.p2 | H3.p3 | TR.p1 | TR.p2 | TR.p3 | TR.int | r16S | rD7 |
|---|---|---|---|---|---|---|---|---|---|---|---|---|---|
| M1 | -132564.7 | | | | | | | | | | | | |
| M2 | -131576.0 | | | | | | | | | | | | |
| **M3** | **-129993.7** | | | | | | | | | | | | |
| M4 | -131637.8 | | | | | | | | | | | | |
| M5 | -132228.4 | | | | | | | | | | | | |
| M6 | -130120.6 | | | | | | | | | | | | |

*Note:* p = codon position, int = intronic region, TR = phenyl-methyl transferase (PMTR)

In all partition schemes we assumed a general time-reversible (GTR) model of molecular evolution and rate heterogeneity among sites. We specified flat Dirichlet priors on both the stationary base frequencies and the exchangeability rates of the GTR models. For all partitions, we modelled among-site heterogeneity according to a discretised single-parameter gamma distribution with four rate categories. We used a lognormal hyperprior on the mean and standard deviation of the gamma-shape parameter to reflect our uncertainty about the variation in substitution rates among sites. We chose parameter values (mean = 5.0, SD = 0.587405) for which 95% of the hyperprior density spans an order of magnitude. We did not estimate both the proportion of invariant sites and the gamma-distributed rate heterogeneity, as the non-independence of these parameters is prone to make their simultaneous estimation inaccurate (Yang 2006).

We used a rooted uniform tree prior, thereby assuming equal prior probabilities for all tree topologies. The root node was fixed to the splitting event between the two Coenagrionoidea families (Platycnemididae and Coenagrionidae). This topological restriction is justified by all available large-scale molecular phylogenies (Carle et al. 2008; Dijkstra et al. 2014; Kim et al. 2014; Toussaint et al. 2019), including recent phylogenomic studies Bybee et al. (2021), all of which strongly support the monophyly of these two families, as well as their relationship as sister clades. Branch lengths were drawn from an exponential distribution (rate = 10.0) and using per-partition rate multipliers drawn from a flat Dirichlet prior, to account for rate variation among data partitions.

Two independent chains for this model were run for 80 000 iterations with a pre-burn in of 20 000 iterations tuning the MCMC proposals. The mixing properties and convergence of these two runs were diagnosed in R v. 4.0.3 (R Core Team 2021) using the coda package (Plummer et al. 2006). Model inference was summarised with the maximum a posteriori (MAP) tree from the combined posteriors of the two independent runs, after an additional burn-in of 60 000 iterations. We then transformed this summary tree into a ultrametric tree, using the *ape* package (Paradis and Schliep 2019) in R, which assumed a correlated molecular clock. This assumption was made only so that this topology inference could be used as a reasonable starting value for the bigeographic dating model below.

#### Biogeographic data

We recorded species distributions across 25 geographic areas spanning the largest land masses worldwide, following Landis (2017) (Table S4, see paleogeographic model below). To do this, we obtained data on the country occurrences of extant species of Coenagrionoidea combining literature, museum and web resources (Table S4). We decided to use countries as geographic units for data collection because this was the finest scale at which several resources reported species distributions and it was possible to obtain information with this resolution for all sampled species. We mined field and identification guides, primary literature, mainly in the form of checklists and taxonomic reviews, IUCN Red List assessments and reputable websites maintained by expert odonatologists with regional species accounts. Also, for most species sampled at two of the largest collections, the Florida State Collection of Arthropods (FSCA) and the New Zealand Arthropod Collection (NZAC) we recorded all countries where specimens had been collected.

For some of the world's largest countries, we also obtained distribution data at the scale of states and territories (Australia, Brazil, India and the USA) provinces and territories (Canada and China) or regions (Russia), whenever possible. Two other countries, Indonesia and Malaysia, also spanned across geographic areas in the paleogeographic model (Table S5; see below), so we divided each of these countries into two regions. For Indonesia we recorded whether species were endemic to the Papua province in the island of New Guinea (AusE) or if they occurred elsewhere in the Malay Archipelago (Mly), and for Malaysia we recorded whether species had island (Mly) or mainland (peninsular; AsSE) distributions. All data on regional and country distributions and their sources will be available on Dryad and in the Odonate Phenotypic Database (Waller et al. 2019) (http://www.odonatephenotypicdatabase.org/shiny/odonates/)

**Table S4. The full table is attached as a tab-delimited file.** Geographic distribution data for the 669 Coenagrionoidea species included in this study. For each species, we obtained distribution data as presence or absence from administrative areas, namely countries and for the world's largest countries (Australia, Brazil, Canada, China, India, Russia, USA), regions, states or provinces, whenever possible. Two other countries (Indonesia and Malaysia) spanned two biogeographic areas. Therefore, we recorded species occurrence

| Taxon | Countries | Regions | Areas | Sources |
|---|---|---|---|---|
| *Acanthagrion adustum* | BRA, COL, GUF, GUY, PAR, SUR, VEN | Amazonas, Para, Rondonia, Roraima | SAmN, SAmE | FSCA, Heckman (2008) |
| *Acanthagrion amazonicum* | BRA | Amazonas, Rondonia | SAmN | Heckman (2008) |
| *Acanthagrion apicale* | BOL, BRA, COL, ECU, GUF, GUY, PER, SUR, VEN | Amazonas, Para, Rondonia | SAmN, SAmE | FSCA, Heckman (2008) |
| *Acanthagrion abunae* | BOL, BRA, COL, GUY, PAN, PAR | Mato Grosso, Rondonia | SAmN, SAmE, SAmC | FSCA, Heckman (2008) |
| ... | ... | ... | ... | ... |

separately for Peninsular Malaysia (coded as AsSE) and the Malaysian Archipelago (coded as Mly), and for the province of Papua (coded as AusE) and the rest of Indonesia (coded as Mly). These distribution data were then used to determine the biogeographic area(s) of occurrence for each species. References to consulted sources are given for each species. In some cases distribution data were taken from or complemented with annotations on museums specimens at the Florida State Collection of Arthropods (FSCA) and the New Zealand Arthropod Collection (NZAC).

#### Empirical paleogeography model

We used the palegeographic model in Landis (2017) to define historical dispersal routes for pond damselflies and featherlegs (superfamily Coenagrionoidea). This empirical model establishes the presence or absence of dispersal paths between all geographic areas during discrete epochs over geological time. The epochs in this model do not necessarily coincide with any of the time units used in the Geological Time Scale. Instead, they reflect periods of time coarsely differentiated by land mass movements that have altered the availability of potential dispersal routes and are not necessarily of equal duration.

To estimate the probability of ancestral dispersal events, all existing biogeographic areas are grouped into communicating classes during each of the discretely defined epochs. Communicating classes are groups of geographic areas that are connected by dispersal routes. This is equivalent to an undirected graph in which the geographic areas represent the vertices and the dispersal routes correspond to the edges. There are three dispersal modes which define different structures of communicating classes. Short-distance dispersal is available only between immediately adjacent areas. Medium-distance dispersal also links these areas as well as areas separated by small water barriers, for example Southern Africa and Madagascar in present-day geography. Finally, long-distance dispersal is unconstrained, linking all geographic areas.

All edges in the dispersal graph for each dispersal mode are piece-wise constant, that is, either present or absent within each epoch. For example, before the split of Pangea, Eastern South America and Western Africa belonged to the same short-distance communicating class, but when a narrow sea was formed between the two continents (epoch 120-110 Ma) this communicating class was broken, and the two areas became connected only by medium- and long-distance dispersal (Landis 2017). The continents then continued to drift apart and after 90 Ma this oceanic barrier is considered wide enough that dispersal is only possible through the long-distance graph.

We adopted this piece-wise-constant graphical-based empirical model from Landis (2017). However, some damselfly species in our sample (n = 22) are endemic to oceanic islands in Micronesia, Polynesia and Hawaii, and their distribution ranges cannot be accommodated within the biogeographic areas of the model (Table S5). In a preliminary analysis, we expanded the paleogeographic model of Landis (2017) to include these islands, as well as the distinct paleogeography of other Archipelagos (Philippines, New Guinea, New Caledonia, Fiji, Vanuatu and Solomon Islands) with high endemism for Zygoptera. However, such an expansion of dispersal matrices resulted in a prohibitively complex model. For the final analysis we therefore used the simpler empirical paleogeographic model in Landis (2017), treating endemics from Micronesian, Polynesian and Hawaiian islands as missing data, and subsuming Philippine endemics within the Malaysian Archipelago (Mly), and Melanesian endemics within Eastern Australia (AusE).

**Table S5.** Geographic areas and state values used in the paleogeographic model of dispersal paths by Landis (2017).

| State | Abbreviation | Name |
|---|---|---|
| 0 | SAmN | South America (N) |
| 1 | SAmE | South America (E) |
| 2 | SAmS | South America (S) |
| 3 | NAmNW | North America (NW) |
| 4 | NAmNE | North America (NE) |
| 5 | NAmSE | North America (SE) |
| 6 | NAmSW | North America (SW) |
| 7 | Grn | Greenland |
| 8 | Eur | Europe |
| 9 | AsiaC | Central Asia |
| A | AsE | Asia (E) |
| B | AsSE | Asia (SE) |
| C | AsNE | Asia (NE) |
| D | AfrW | Africa (W) |
| E | AfrS | Africa (S) |
| F | AfrE | Africa (E) |
| G | AfrN | Africa (N) |
| H | AusW | Australia (W) |
| I | AusE | Australia (E) |
| J | Ind | India |
| K | Mdg | Madagascar |
| L | AntW | Antartida (W) |
| M | AntE | Antartida (E) |
| N | Mly | Malaysian Archipelago |
| O | NZ | New Zealand |

#### Biogeographic dating with empirical paleogeography

We used the MAP tree from our initial phylogenetic inference as an observed topology for the biogeographic dating model (Landis 2017), using RevBayes v 1.0.12 (Höhna et al. 2016). With this statistical approach, we simultaneously inferred branching times and biogeographic history in the Coenagrionoidea phylogeny. This model is novel in that it explicitly considers uncertainty about the ease of dispersal of the lineages in question. To accomplish this, a weighing parameter is estimated for each dispersal mode. The weighing parameters are forced to add up to one and are used to rescale the instantaneous matrix of transition probabilities between geographic areas.

The biogeographic process that informs the absolute speciation times is modelled as a time-heterogenous continous-time Markov Chain (CTMC). The transition probabilities between geographic states depend on $\mathbf{Q}(\kappa)$, the instantaneous rate matrix for each epoch $\kappa$, and on the clock rate $\mu$. For each epoch, the rates in $\mathbf{Q}$ depend on the available dispersal routes, defined by empirical paleogeography as explained above, and the weight of the different dispersal modes. We used a strong prior on the weighing parameter of the alternative dispersal modes as in Landis (2017), assuming short-distance dispersal occurred with an order of magnitude higher probability than medium-distance dispersal, in turn occurring at an order of magnitude higher rate than long distance dispersal.

We linked the biogeographic clock to the molecular clock by a rate multiplier with an uniform prior bounded between 0.001 and 1000, so inference on the biogeographic process would inform the absolute timing of phylogenetic branching events. For the clock model, we assumed uncorrelated branch rates drawn from a discretised lognormal distribution with 8 rate categories. To constrain variance in the distribution of relaxed clocks, an exponential hyperprior with rate = 0.1 was placed for the standard deviation of the lognormal. The mean of the lognormal was given by the cumulative molecular evolution and the root age, both of which are in turn random variables. As Landis (2017), we used a broad uniform prior on the tree length (in terms of molecular substitutions per site) between 0 and 10 000. As in the topology inference, nucleotide substitutions were modelled under a GTR process for each partition and assuming gamma distributed site-rate heterogeneity with 4 rate categories. For the root age, we used two alternative priors in separate analyses.

First, a weak uniform prior bounded the root age between 240 and 40 Ma. Crown Odonata (i.e. all extant dragonflies and damselflies) probably arose shortly after the Permia-Triassic mass extinction about 250 Ma (Misof et al. 2014; Suvorov et al. 2021; Kohli et al. 2021). Our weak prior therefore prevents sampling origin times for Coenagrionoidea prior to the presumed common ancestor of the entire order. The minimum age of 40 Ma was based on the lower limit of the 95% HPD interval [44, 80] inferred for this node in a previous multi-locus study (Waller and Svensson 2017), and is also below the 95% HPD interval [112, 125] in a recent phylogenomic study (Suvorov et al. 2021).

Second, a strong root-age prior was informed by two recent phylogenomic studies based on different data sets (Suvorov et al. 2021; Kohli et al. 2021). Kohli et al. (2021) and Suvorov et al. (2021) estimated the origin of Coenagrionoidea at 109 Ma and 116 Ma respectively. We thus drew the root age in this analysis from a truncated normal distribution with mean = 120, sd = 20, minimum = 40, maximum = 240. This choice of prior density follows a strong belief that posterior origin times for Coenagrionoidea in previous phylogenomic studies (Suvorov et al. 2021) are unbiased and adequately capture node-age uncertainty. The results of this informed analysis are presented in the main text and used for subsequent estimation of diversification rates, dispersal and biome-shift histories.

By contrasting results of analyses using a weak (uniform) and a strong (normal) prior we assessed how a strong belief in the results of previous phylogenomic studies under the biogeographic dating model affects node age estimates across the Coenagrionoidea tree. We also conducted an additional analysis with a broader uniform prior on the root age, bounded between 240 and 0 Ma and ignoring the empirical paleogeographic model. This analysis was done to confirm that without paleogeographic information the prior and posterior root age distributions are essentially the same, and therefore, dating estimates are informed by the empirical paleogeography and not an artefact of model construction.

To improve sampling of the posterior root age in all analyses, we use three different moves (mvScale with starting strengths of 1.0 and 0.1 and mvSlide) and 10 proposals for each move in each iteration of the MCMC algorithm. Branch-specific model parameters received one proposal per iteration while the remaining model parameters received between 10 and 20 proposals per iteration. Each model was run for 30 000 iterations of which 20% were discarded as burn-in. As for the topology inference, dating inference was summarised with a MAP tree for plotting and downstream diversification analyses.

#### Comparison to dating under fossil constraints

We conducted an additional dating analysis using fossil constraints instead of empirical paleogeography to explore how these alternative approaches compare to each other and to node-age estimates from previous studies. The molecular evolution module of this model was as similar as possible to the biogeographic dating analysis. We used the same GTR model of substitution rates with gamma-distributed site-rate heterogeneity, and an uncorrelated relaxed clock model, discretised to eight rate categories. As there is no link between biogeographic and molecular clocks in this model, we used an exponential hyperprior with rate = 2 for the mean of the lognormal distribution of branch rates. Also, as in our biogeographic dating, the tree module in this analysis used a normal distribution (mean = 120, sd = 20), truncated between 240 and 40 Ma, as the root age prior. In contrast to the paleogeographic dating model, here we used a pure birth process as the tree prior. Diversification events followed a lognormal distribution, with mean = 0.059, based on the total number of species described in Coenagrionoidea (~1 805) and the root age estimate in Suvorov et al. (2021) (116 Ma), and sd = 2.0.

The key feature of this analysis is that node ages were further informed by fossil constraints, as opposed to any paleogeograhic data. We revised the literature on fossils assigned to Coenagrionoidea in the Paleobiology Database https://paleobiodb.org/, and identified only four fossils that adequately meet the criteria for inclusion outlined by Parham et al. (2012). These fossils have been each identified as crown members of the genera *Argia* (Zheng et al. 2018), *Nehalennia* (Ross et al. 2016) and *Ischnura* (Bechly 2000), and the family Platycnemididae (Poinar Jr et al. 2010). Fossils were therefore used to constrain the minimum ages of these clades to 16, 16, 13.7 and 93.9 Ma, respectively.

In RevBayes, fossil observations are treated as offsets on the ages of internal nodes, which are in turn stochastic variables in the model. For example, the MRCA of the genus *Argia* is offset by 16 My, according to fossil evidence (Zheng et al. 2018). This means that all MCMC samples with younger ages for this node will be discarded, resulting in a hard-minimum age calibration. RevBayes's node-dating approach circumvents the use empirical node-age priors which may be incompatible with the tree-wide pure birth prior. However, this does not imply that realized and specified priors will be identical. Thus, we first ran the MCMC analysis without any molecular data or fossil constraints and confirmed that we recovered the specified root age prior as the posterior (mean = 119.17, sd = 20.13). In this and the subsequent analysis with data, we accounted for missing taxa at the superfamily level by specifying the sampling fraction (Höhna et al. 2011; Höhna 2014), according to the World Odonata List (Paulson and Schorr 2021). The model was run for 60,000 iterations, after 10,000 iteration were discarded as burn-in and with multiple proposals on each parameter per iteration, as in the paleogeographic dating model.

#### Diversification analysis

##### Diversification through time

We used the episodic birth-death (EBD) process (Stadler 2011; Höhna 2015) to model the temporal dynamics of diversification in pond damselflies and featherlegs. EBD models enable detection of temporal shifts in speciation and extinction rates in a likelihood framework, while accounting for incomplete taxon sampling. The EBD process assumes that diversification rates vary in a piecewise constant manner, that is, rates are constant within time intervals (episodes) but vary between intervals. To reflect our uncertainty on current diversification rates in Coenagrionoidea, we used uniform distributions bounded between -10.0 and 10.0 for the log speciation and extinction rates of the latest time interval (ending at the present). To estimate diversification rates in previous intervals, we assumed that rates were temporally autocorrelated and followed a Horseshoe Markov random field (HSMRF) prior distribution (Magee et al. 2020). This means that diversification rates in each time interval were centered around the mean rate of the previous interval, going backwards in time.

Following the HSMRF parametrisation in Magee et al. (2020), differences in log-scale speciation and extinction rates between time intervals were drawn from a normal distribution, with mean 0 and standard deviation controlled by global and local scale priors. Both scale priors were drawn from a Half-Cauchy with location = 0 and scale = 1, and were multiplied by a global scale hyperprior. The appropriate value for the hyperprior depends on the number of intervals and can be approximated following Magee et al. (2020). This parametrisation of the HSMRF model is particularly useful to detect when rapid shifts occur occasionally in a background of slowly changing diversification rates over most time intervals (Carvalho et al. 2010). Here, we used 20 equal-length time intervals between the root age and the present, and a global scale hyperprior of 0.016.

We accounted for our empirical taxon sampling strategy (Höhna et al. 2011; Höhna 2014) by specifying the number of missing taxa for each of the major clades within Coenagrionoidea, that were identified in our initial topology inference and in previous phylogenetic studies: the featherlegs (family Platycnemididae), the 'core' pond damselflies (*sensu* Dijkstra et al. 2014) the dancers (genus *Argia*), the neotropical threadtails and the remainder of the 'ridge-face' pond damselflies (*sensu* Dijkstra et al. 2014) (Fig. S4). Two independent chains of the model were run for 50 000 iterations with a pre-burn-in of 10 000 iterations and tuning proposals every 200 iterations. Model parameters receive 5-10 proposals per iteration. Diversification through time analyses were conducted separately for the MAP trees from strongly-informed and weakly-informed dating models.

##### Biome-dependent diversification

We used Hidden-State Dependent Speciation and Extinction (HiSSE) models to investigate how diversification rates have varied across latitude-delimited biomes (Tropical, warm-temperate and cold-temperate). HiSSE models are in essence similar to Binary-State and Multi-State Dependent Speciation and Extinction models (Maddison et al. 2007; FitzJohn 2012), but they accommodate diversification rate shifts that are independent of the trait of interest by incorporating an additional unobserved character. This hidden trait accounts for background diversification heterogeneity across branches (Beaulieu and O'meara 2016). Thus HiSSE models help overcome the high susceptibility to type I errors, that seems to undermine BiSSE and MuSSE models in cases where additional traits influence diversification (Maddison and FitzJohn 2014; Rabosky and Goldberg 2015).

We assumed a binary-state hidden character and modelled biome-state transitions as anagenetic changes. Latitudinal biomes were assigned to each taxa based on present-day geography, using distribution data at the level of administrative areas (Table S4), and biome reconstructions for the last 5 My (Jolly et al. 1998; Salzmann et al. 2008; Otto-Bliesner et al. 2020). Taxa with distribution ranges spanning across biomes were assigned to the coldest biome they occupy (results in the main text) or were coded as ambiguous between their current states (results below).

We used exponential priors with a mean of 10 for the transition rates between biomes and for transitions between states of the hidden character, which were assumed to be symmetrical. We used a log-normal prior on the trait-dependent speciation and extinction rates. Also, we assumed equal prior distributions for speciation and extinction rates, with mean equal to the expected net diversification rate and standard deviation = 2.0. Differences in log-scale diversification rates between hidden states were in turn drawn from an exponential distribution (rate = 1) for speciation and a normal distribution (mean = 0, sd = 1.0) for extinction.

We first compared the HiSSE model to a null model with background heterogeneity in diversification rates, but with equal speciation and extinction rates in all biomes. Models were compared using marginal likelihood approximation via the stepping stone algorithm (Xie et al. 2010). As differences in diversification rates between biomes were strongly supported (see Results), we proceeded to run the HiSSE model using the MAP tree from our strongly-informed dating analysis, and the two alternative character matrices (with and without ambiguous states) as input. For each data set, two replicates of the model were run for 50 000 iterations, with a pre-burn-in of 10 000 iterations in which parameter proposals were updated every 200 iterations. Two to five moves were proposed for each model parameter in each iteration. An additional 20% burn-in was applied before summarizing posterior rates and ancestral states. To confirm that our results are not a product of model parametrisation, we also conducted an analysis under the prior (i.e. without data). Finally, we tested the robustness of our results to our previous dating inferences by also estimating state-dependent diversification rates on the MAP tree from weakly-informed dating analysis.

#### Modelling biome shifts and dispersal events

To understand how lineage movements have contributed to the latitudinal diversity gradient (LDG) in Coenagrionoidea we jointly modelled shifts between latitude-delimited biomes and dispersal events across regions, following the approach developed by Landis et al. (2021). This approach uses literature-informed graphs to capture the availability and connectivity among paleobiomes and paleoregions and a time-stratified model of transitions between compound biome-region states. Here, we first describe the features of the empirical connectivity graphs used in the analysis, and then provide an overview of the model developed in Landis et al. (2021) and used here. Finally, we explain how stochastic histories were used to estimate the frequency and timing of different types of biogeographic events that have contributed to global diversity patterns today.

##### Empirical paleobiome-region model

Landis et al. (2021) constructed graphs to represent the availability and connectivity of mesic forests (i.e. forest with intermediate levels of moisture content) over the last 100 My. These graphs were informed by paleobiological and paleogeographic literature, focusing on the six regions occupied by extant representatives of their study system (the plant genus *Viburnum*). Damselflies in the superfamily Coenagrionoidea originated approximately 105 Ma (see Results), so the temporal scope of Landis et al. (2021) was also suitable for our study system. We also adopted the distinction of eight epochs from the Late Cretaceous to the present, each with their own set of time-dependent graphs (Table S6).

For each epoch, paleogeographic graphs determined land connectivity between regions, in a similar way as in our biogeographic dating model. However, due to computational constraints when combining biome and region states, here we recoded our distribution data to a reduced number of region states. The analysis included the six regions used in Landis et al. (2021) (Southeast Asia, East Asia, Europe, South America, Central America, and North America), and three additional regions that contain an important fraction of pond damselfly and featherleg diversity (Africa, Australia, and India). The extent of land connectivity between regions was determined by applying strong, weak or marginal features to each pair of regions, analogous to the short-, medium-, and long-distance dispersal routes in the biogeographic dating model. For example, present-day connectivity between South America and Africa is marginal (i.e. long-distance dispersal would be required for a lineage to move between these regions), but in the Late Cretaceous, when the continents were closer together, weak connectivity was assumed.

In addition to paleoregions, used to model dispersal events, we constructed graphs to represent the availability and connectivity of paleobiomes within regions. Similarly to Landis et al. (2021), we distinguished three broad latitude-delimited biomes, Tropical, warm-temperate and cold-temperate. For each biome and epoch, a strong feature indicated high regional coverage of $\geq 25\%$, a weak feature indicated regional coverage of $< 25\%$, and a marginal feature indicated regional coverage of $< 1\%$. In the model, biome shifts occur within regions and, if ecological features determine lineage movements, biome shifts should occur with higher probability as the coverage of the incoming biome increases. While we refer to these biomes as latitude-delimited, their geographic boundaries have changed with the temperature trends of Earth's history. For example, cold-temperate biomes were essentially unavailable during the Hothouse and Warmhouse periods prior to the Oligocene, and have since increased in coverage, first in the Northern Hemisphere and more recently in the South (Fig. S2-S3).

Paleobiome graphs were based on those constructed in Landis et al. (2021). However, as we expanded the number of regions compared to this previous study, we expanded the paleobiome graphs accordingly. Literature consulted to determine biome availability in each of the newly included regions are reported for each epoch (Table S6). We note that, unlike *Viburnum* in Landis et al. (2021), pond damselflies and their relatives inhabit biomes ranging from xeric to very humid. However, as the focus of the present study is on the LDG, and the geographic scope of our analysis is global (resulting in an already large state space), here we explore only the temperature component of biomes. We represent the expanded paleoregion and paleobiome graphs as schematic adjacency matrices, together with a uninformative set of matrices indicating the null scenario of lineage movements being unconstrained by either water barriers or ecological affinities (Fig. S2-S3).

**Table S6.** Epochs during the last 100 My used in Landis et al. (2021) and in the present study to model time-heterogeneous dispersal and biome shifts. Global temperature trends are based on O'Brien et al. (2017) and Westerhold et al. (2020). For the three additional regions included in this study, we provide literature references describing paleobiomes in each epoch.

| Epoch | Start (Ma) | Temperature trend | Sources for Africa | Sources for India | Sources for Australia |
|---|---|---|---|---|---|
| Late Cretaceous | 106.0 | Cooling trend from Hothouse peaking ~ 100-90 Ma | Maley (1996), Jacobs (2004); Jacobs et al. (2010), Warny et al. (2019) | Specht et al. (1992); Carpenter et al. (2015); Rundel et al. (2016); Otto-Bliesner et al. (1997); Ohba and Ueda (2010) | Otto-Bliesner et al. (1997); Ohba and Ueda (2010); Dzombak et al. (2020) |

*(continued)*

| Epoch | Start (Ma) | Temperature trend | Sources for Africa | Sources for India | Sources for Australia |
|---|---|---|---|---|---|
| Paleocene | 66.0 | Stable Warmhouse | Jacobs (2004); Jacobs et al. 2010; Neumann and Bamford (2015) | Hill and Hall (2002); Greenwood et al. (2003); Barreda and Palazzesi (2007); Bowman et al. (2014) | Prasad et al. (2018); Bhatia et al. (2021) |
| Early Eocene | 56.0 | Heating to thermal maximum Hothouse ~ 55 Ma | Herold et al. (2014); Neumann and Bamford (2015) | Hill and Hall (2002); Pross et al. (2012); Herold et al. (2014) | Utescher and Mosbrugger (2007); Herold et al. (2014) |
| Mid/Late Eocene | 48.0 | Continuous cooling back to Warmhouse | Herold et al. (2014); Neumann and Bamford (2015); Pound and Salzmann (2017) | Hill and Hall (2002); Pound and Salzmann (2017) | Utescher and Mosbrugger (2007); Singh et al. (2011); Pound and Salzmann (2017) |
| Oligocene | 33.9 | Cooling to stable Coolhouse | Neumann and Bamford (2015); Pound and Salzmann (2017) | Hill and Hall (2002); Buerki et al. (2013); Pound and Salzmann (2017); Korasidis et al. (2019) | Kent and Muttoni (2008); Srivastava et al. (2012); Pound and Salzmann (2017); Su et al. (2018) |
| Early Miocene | 23.0 | Stable Coolhouse | Jacobs (2004); Vincens et al. (2006); Neumann and Bamford (2015); Roberts et al. (2017) | Hill and Hall (2002); Byrne et al. (2008); Herold et al. (2011) | Guo et al. (2008); Potter and Szatmari (2009); Klaus et al. (2016); Chen et al. (2019) |
| Mid/Late Miocene | 16.0 | Continuous cooling to Icehouse | Pound et al. (2011); Pound et al. (2012) | Pound et al. (2011); Pound et al. (2012) | Pound et al. (2011); Pound et al. (2012) |
| Recent | 5.3 | Icehouse | Jolly et al. (1998); Salzmann et al. (2008); Otto-Bliesner et al. (2020) | Salzmann et al. (2008); Otto-Bliesner et al. (2020) | Salzmann et al. (2008); Otto-Bliesner et al. (2020) |

##### Time-stratified model of compound biome-region states

The model in Landis et al. (2021) addresses how geographic and ecological opportunities influence lineage movements, by jointly modelling two interacting subprocesses: dispersal events and biome shifts. Dispersal events occur when a lineage colonizes a novel region, for example, if a member of a Southeast Asian clade reaches Australia. Critically, dispersal events depend on both land connectivity and continuity of a suitable biome. For instance, a temperate North American lineage would more likely disperse to South America during periods of time in which both land masses are connected by a land bridge and in which temperate biomes have extensive coverage in the South. As mentioned above, the biome shift process, when lineages move to warmer or colder habitats, occurs within regions, and thus depends on the availability and coverage of the incoming biome in the present region. The adjacency matrices described in the previous section inform the

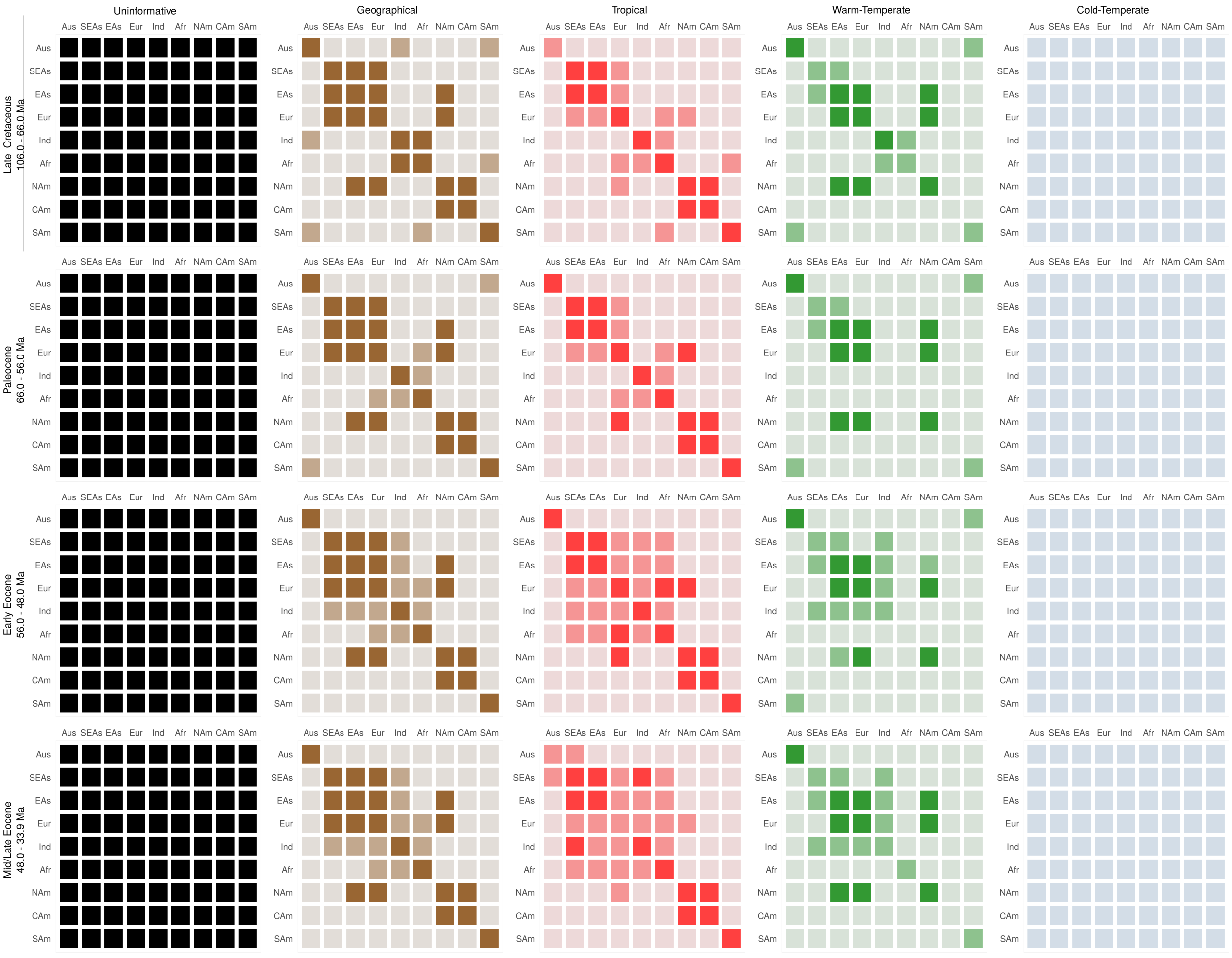

**Figure S2.** Availability and connectivity of null features, paleoregions and paleobiomes from the Late Cretaceous to the Mid/Late Eocene. Rows represent epochs and columns correspond to regional features. The model includes four types of features: uninformative (i.e. null; black), geographical (brown), and biome-specific features for Tropical (red), warm-temperate (green), and cold-temperate (blue) biomes. Cells are shaded to represent features as being strong (dark), weak (medium), or marginal (light). Regional availability is encoded in the diagonal elements of the matrices, while connectivity between regions is represented by off-diagonal elements. Illustration of adjacency matrices is based on Landis *et al.* (2021).

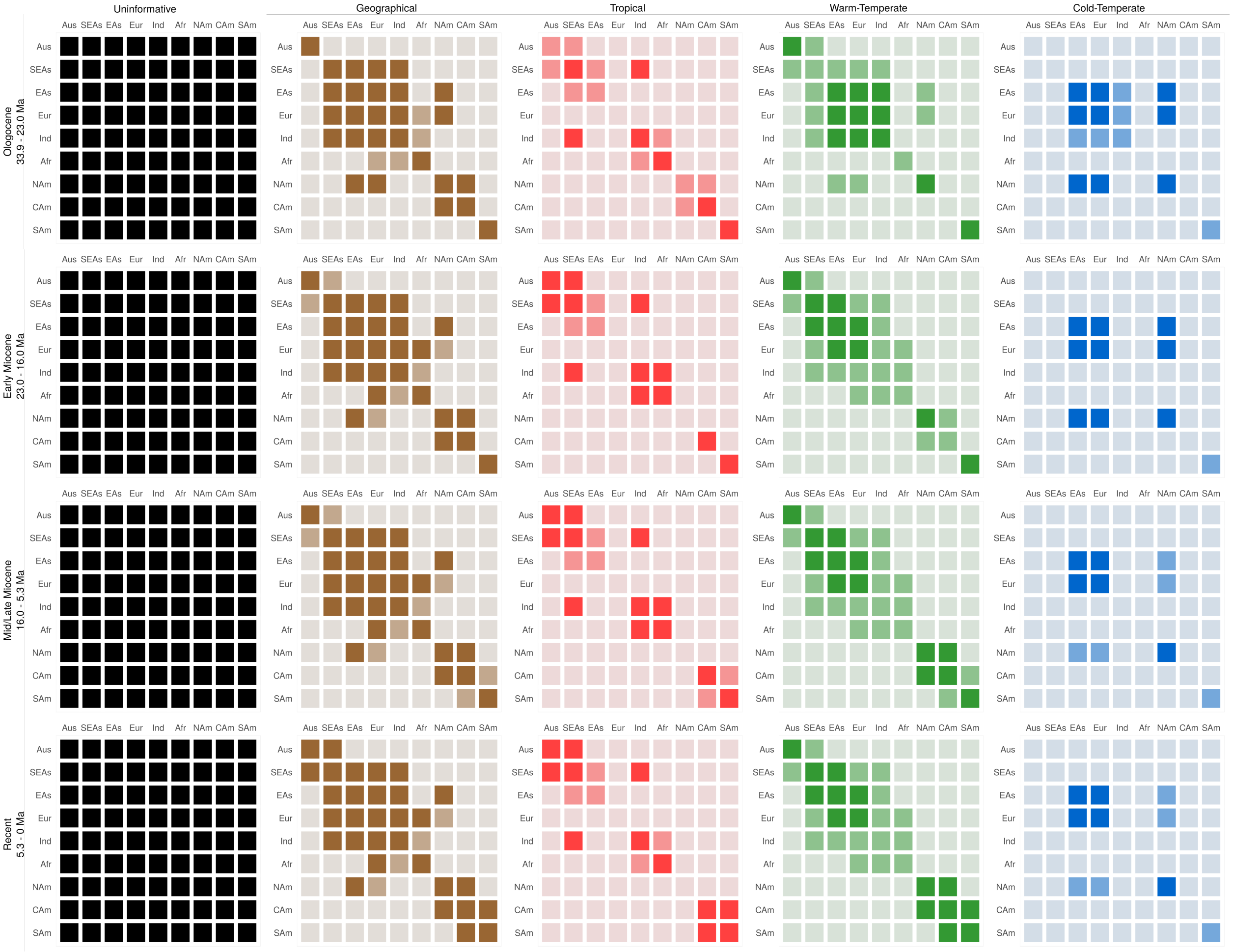

**Figure S3.** Availability and connectivity of null features, paleoregions and paleobiomes from the Oligocene to the present (Recent). Rows represent epochs and columns correspond to regional features. The model includes four types of features: uninformative (i.e. null; black), geographical (brown), and biome-specific features for tropical (red), warm-temperate (green), and cold-temperate (blue) biomes. Cells are shaded to represent features as being strong (dark), weak (medium), or marginal (light). Regional availability is encoded in the diagonal elements of the matrices, while connectivity between regions is represented by off-diagonal elements. Illustration of adjacency matrices is based on Landis *et al.* (2021).

probability of each of these events in a time-specific manner, accounting for continental drift and historical changes in the steepness of the LDG.

The joint dispersal and biome shift processes are modelled using a continuous-time Markov chain with time-stratified rates between compound biome-region states (Table S7), which captures the interdependence between biome and region characters. Details on how transition probabilities are computed are given in Landis et al. (2021). Here, we provide an overview of some of the features of the model implementation that are most relevant to our study.

First, the instantaneous rate matrix for the time interval $m$, $Q(m)$, which defines the transition probabilities between biome-region states, is a weighted average of three rate matrices: the uninformative (time and context-independent) rate matrix, $\mathrm{Q}_U$, the geographical (biome-independent) rate matrix, $\mathrm{Q}_G(m)$, and the biome rate matrix, $\mathrm{Q}_B(m)$. Thus, by estimating the matrix feature weights ($\mathrm{w}_U$, $\mathrm{w}_G$, $\mathrm{w}_B$) from the data we can evaluate the relative importance of land and biome connectivity in constraining lineage movements.

We would also like to learn from the data the extent to which strong and weak features influence lineage movements. Landis et al. (2021) accomplish this by specifying a free parameter $y$ that controls the effect to weak features relative to marginal (fixed to 0) and strong features (fixed to 1) across all geographical and biome adjacency matrices in Fig. S2-S3.

Importantly, because transitions between biome-region states are neither symmetric nor equal over time, biome-region states can exhibit different source-sink dynamics during different epochs. For example, biome-regions with low accessibility are expected to gradually lose lineages, as these states receive few dispersing lineages from other regions and lose occupants to biome shifts towards more extensive biomes within the same region. Approximating the stationary distributions of these time-specific source-sink dynamics can therefore be used to estimate the proportion of lineages in each biome-region state for a given time interval.

Finally, the model in Landis et al. (2021) makes several assumptions, whose implications warrant further investigation. First, a lineage cannot shift to a new biome and disperse to a new region at the same time. This assumption implies that the order of events matters and may be problematic if dispersal events are in fact not restricted by ecological affinities. Second, the model assumes no time or state-dependent variation in speciation and extinction. However, our diversification analyses suggest otherwise. The potential consequences of violating this assumption require further study, but it seems plausible that biome shifts and dispersal events towards speciation-enhancing states could be overestimated, while transitions to extinction-enhancing states would be underestimated (Maddison 2006). Third, the models assumes that lineages are predominantly present in a single state. While this is true for the majority of pond damselflies and featherlegs, and substantially reduces the state space of the model, range size varies vastly in Coenagrionoidea (Fig. 1) and even within genera (Blow et al. 2021).

Two independent chains of the model were run in RevBayes v 1.1.1 (Höhna et al. 2016), for 25 000 iterations with 5 000 iterations of burn-in. Parameter values were sampled every 50 iterations. As in previous analysis, each random variable was subjected to multiple moves in each iteration of the Markov chain. We visually assessed convergence between the two runs and stationarity of the parameter posteriors. The MAP tree of our previous strongly informed dating analysis was used as the tree input. Biome-region data for extant taxa was obtained from our geographic distribution data set (Table S4), using data collected at the level of administrative areas, and biome reconstructions for the *Recent* epoch (5.3 - 0 Ma) (Jolly et al. 1998; Salzmann et al. 2008; Otto-Bliesner et al. 2020).

**Table S7.** Biome-region states, abbreviations, and symbols, used in the dispersal and biome shift model.

| State | Abbreviation | Symbol |
|---|---|---|
| Tropical Australia | Tropical+Aus | A |
| Tropical Southeast Asia | Tropical+SEAs | B |
| Tropical East Asia | Tropical+EAs | C |
| Tropical Europe | Tropical+Eur | D |
| Tropical India | Tropical+Ind | E |
| Tropical Africa | Tropical+Afr | F |
| Tropical North America | Tropical+NAm | G |
| Tropical Central America | Tropical+CAm | H |
| Tropical South America | Tropical+SAm | I |
| Warm-Temperate Australia | Warm+Aus | J |
| Warm-Temperate Southeast Asia | Warm+SEAs | K |
| Warm-Temperate East Asia | Warm+EAs | L |
| Warm-Temperate Europe | Warm+Eur | M |
| Warm-Temperate India | Warm+Ind | N |
| Warm-Temperate Africa | Warm+Afr | O |
| Warm-Temperate North America | Warm+NAm | P |
| Warm-Temperate Central America | Warm+CAm | Q |
| Warm-Temperate South America | Warm+SAm | R |
| Cold-Temperate Australia | Cold+Aus | S |
| Cold-Temperate Southeast Asia | Cold+SEAs | T |
| Cold-Temperate East Asia | Cold+EAs | U |
| Cold-Temperate Europe | Cold+Eur | V |
| Cold-Temperate India | Cold+Ind | W |
| Cold-Temperate Africa | Cold+Afr | X |
| Cold-Temperate North America | Cold+NAm | Y |
| Cold-Temperate Central America | Cold+CAm | 0 |
| Cold-Temperate South America | Cold+SAm | Z |

#### Summary of model inferences

Following Landis et al. (2021), we used ancestral state reconstruction and stochastically mapped character histories to summarise our model inferences. We plot the posterior probabilities for the three most probable biome-region states in each internal node, based on ancestral state estimates. We use the posterior distribution of stochastically mapped character histories to obtain lineage-state proportions through time. We do this by computing the posterior mean count of lineage-states for each time bin (of 1 My), then dividing by the number of lineages present during that interval.

To assess the congruence between model inferences and empirical paleobiome availability, we classified each lineage-state in each time bin as congruent or not congruent with any locally extensive biomes. Non congruent lineage-states are those in which the lineage's biome had only a marginal presence in the lineage's region and epoch. Non congruent states were labelled as biome mismatches and congruent states were labelled as biome matches. We then computed the proportion of biome matches and mismatches for each time bin.

Finally, we computed the proportion of different ordered *event series* as in Landis et al. (2021). These *event series* correspond to two consecutive state transitions that result in changes in regions, biomes or both (Table S8). For example, a *biome-first* series designates a biome shift that is followed by a dispersal event. In addition to *biome-first* and *region-first* series we recorded biome and region *reversals* and biome and region *flights* (Table S8). The proportion of each *event series* type was computed for each posterior sample by classifying stochastically mapped state triplets (e.g. Tropical+CAm to Tropical+NAm to Warm+NAm) using a root-to-tip recursion described in Landis et al. (2021). We report the 95% highest posterior density interval (HPD) and the 80% HPD for each *event series* type across all posterior samples in our analysis.

**Table S8.** *Event series* involving two state transions between biomes (A, B, C) and/or regiones (X, Y, Z).

| Series name | Event 1 | Event 2 |
|---|---|---|
| biome-first | AX to BX | BX to BY |
| region-first | AX to AY | AY to BY |
| biome reversal | AX to BX | BX to AX |
| region reversal | AX to AY | AY to AX |
| biome flight | AX to BX | BX to CX |
| region flight | AX to AY | AY to AZ |

### Extended Results

#### Tree topology inference

Our phylogenetic inference for Coenagrionoidea identified five main clades (Fig. S4) also identified with the most recent backbone phylogeny for Odonata (Bybee et al. 2021) and other recent phylogenies aimed at resolving inter-generic relationships in Coenagrionoidea (Dijkstra et al. 2014; Toussaint et al. 2019). Nonetheless, some of the relationships among and within these main clades remain at odds across recent studies (Fig. S5). We note that some nodes in the maximum *a posteriori* (MAP) tree that summarises our phylogenetic inference were highly uncertain (Fig. S6-S8). Of 668 internal nodes ~60% are supported by a posterior probability > 90%, and ~77% of the internal nodes are supported by posterior probability > 50% (Fig. S9). Nodes that are supported by high posterior probabilities in our phylogenetic inference are generally congruent across studies (see Results) or, alternatively, congruent between the present study and the backbone phylogeny for Odonata (Bybee et al. 2021) (Fig. S5b).

We also investigate whether monophyly is supported within the genera of Coenagrionoidea and how our results contrast previous molecular and morphological studies. For each polytypic genus sampled, we recorded the cladistic implications of our phylogentic inference - i.e. whether the genus is monophyletic or paraphyletic in the MAP tree (Table S9). We also noted relationships supported in previous studies and whether the cladistic implications of our analysis are congruent with these previous studies (Table S9). Below, we review each of the clades containing putatively paraphyletic genera. We discuss the strength of the evidence for our inferred phylogenetic relationships in the light of previous molecular and morphological studies.

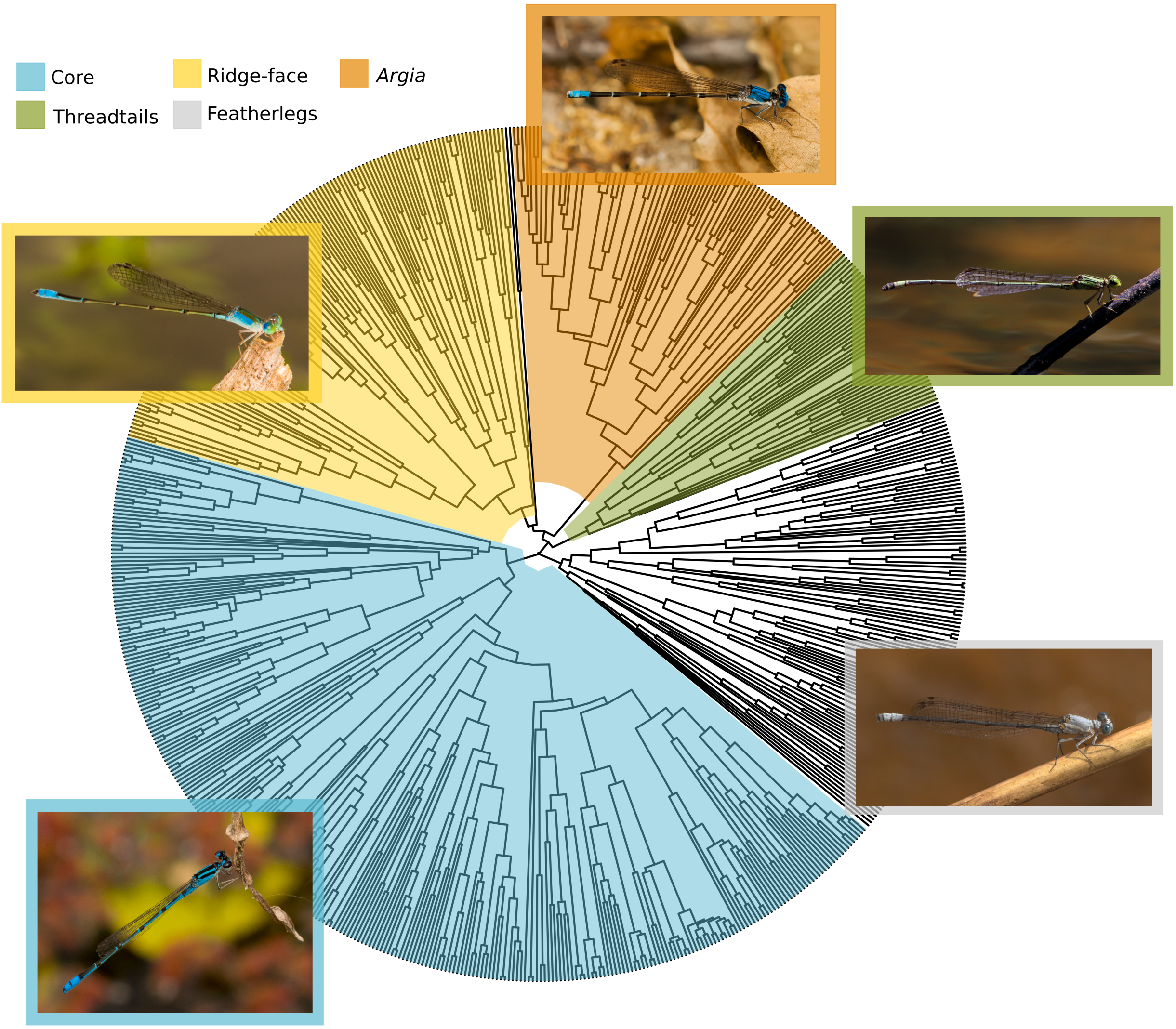

**Figure S4.** Coenagrionoidea phylogeny showing the five main clades identified in the topology inference and used to specify sampling fractions in the episodic birth-death (EBD) model. The illustrated species and sampling fractions are as follows: 'core' pond damselflies (N = 290/672): *Acanthagrion temporale*, dancers (N = 91/132): *Argia apicalis*, neotropical threadtails (N = 44/107): *Neoneura confundens*, other 'ridge-face' pond damselflies (N = 131/449): *Telebasis demarara*, featherlegs (N = 113/445): *Mesocnemis singularis*. Photos: EIS and Erland Refling Nielsen (*N. confundens*).

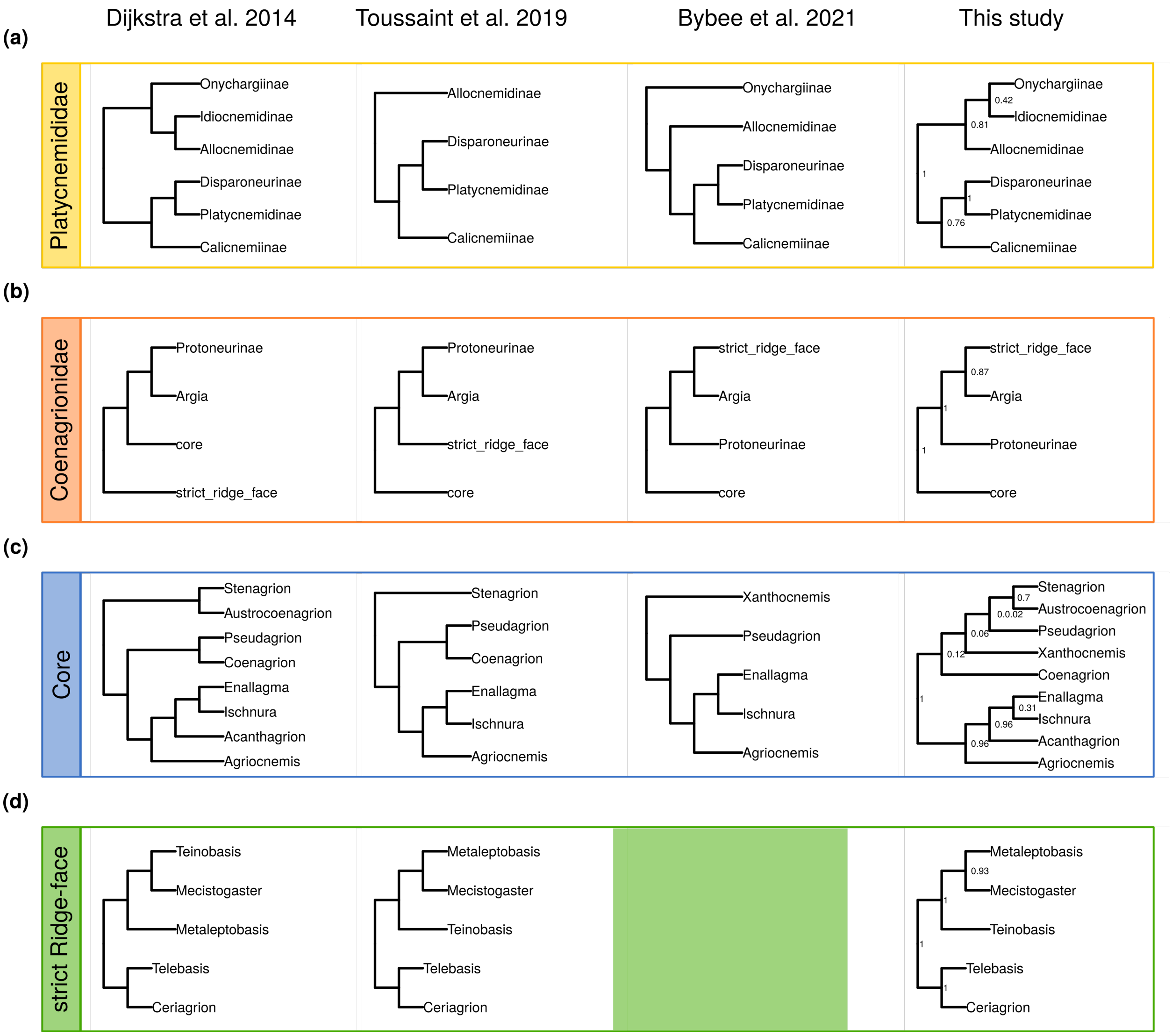

**Figure S5.** Summary of phylogenetic relationships among the main clades and some of the best represented genera of Coenagrionoidea. We compare the phylogenetic inference in the present study to three previous studies with a broad taxonomic scope. Internal node labels represent posterior probabilities (PP) in the maximum *a posteriori* tree of Coenagrionoidea, presented in this study. **a** Relationships among subfamilies of featherlegs (Platycnemididae). **b** Relationships among the main clades of pond damselflies (Coenagrionidae). **c** Relationships among the best represented genera of 'core' pond damselflies (Coenagrionidae). **d** Relationships among the best represented genera of strict-sense 'ridge-face' pond damselflies (Coenagrionidae). Here, the strict-sense 'ridge-face' clade of pond damselflies refers to the clade described in Dijkstra textitet al. (2014), whereas the broad-sense 'ridge-face' clade (or simply the ridge-face clade in this study) also includes the dancers (genus *Argia*) and the subfamily Protoneurinae.

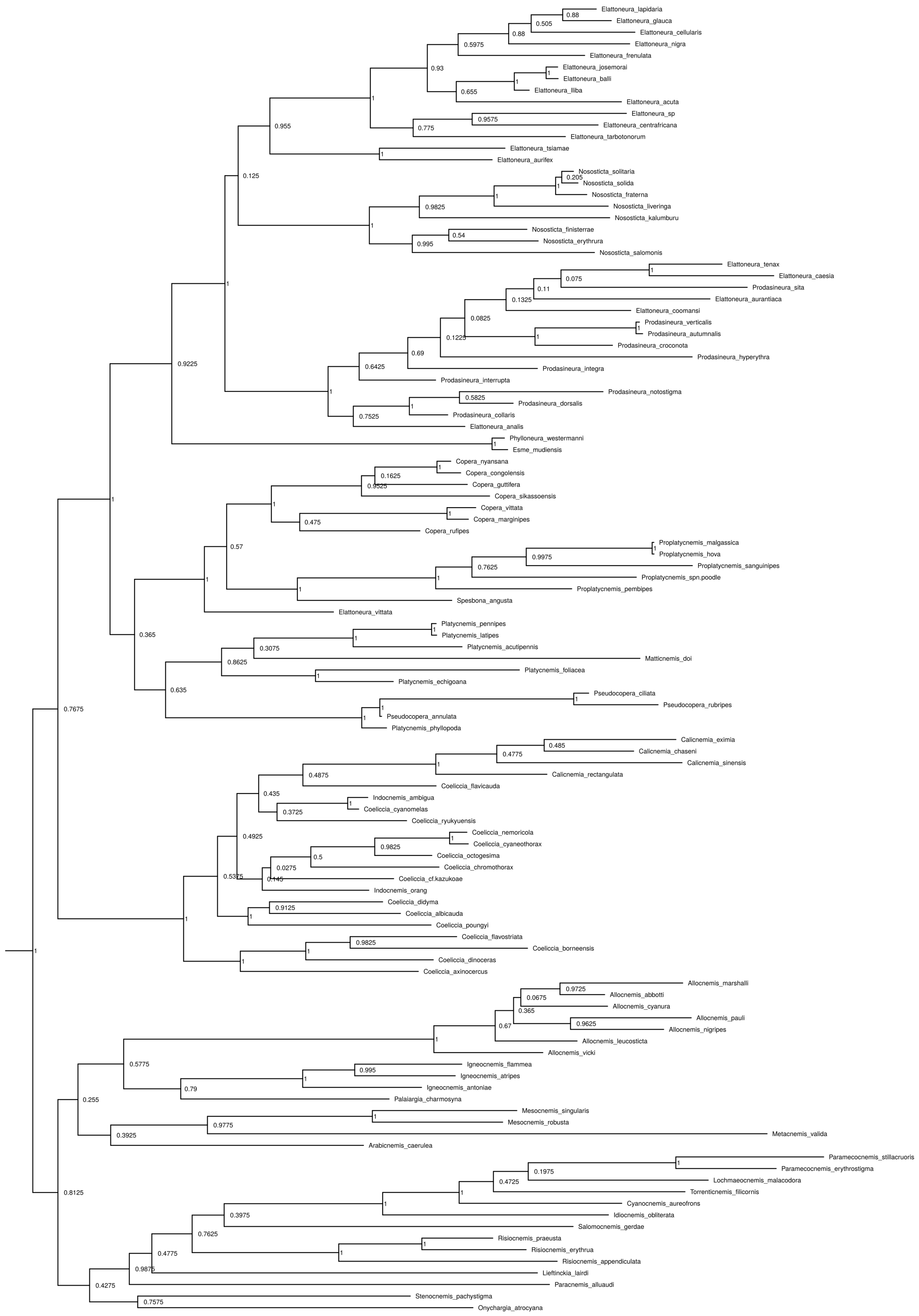

**Figure S6.** Phylogenetic relationships among featherlegs (family Platycnemididae) in the maximum *a posteriori* tree of Coenagrionoidea. Internal node labels represent posterior probabilities (PP).

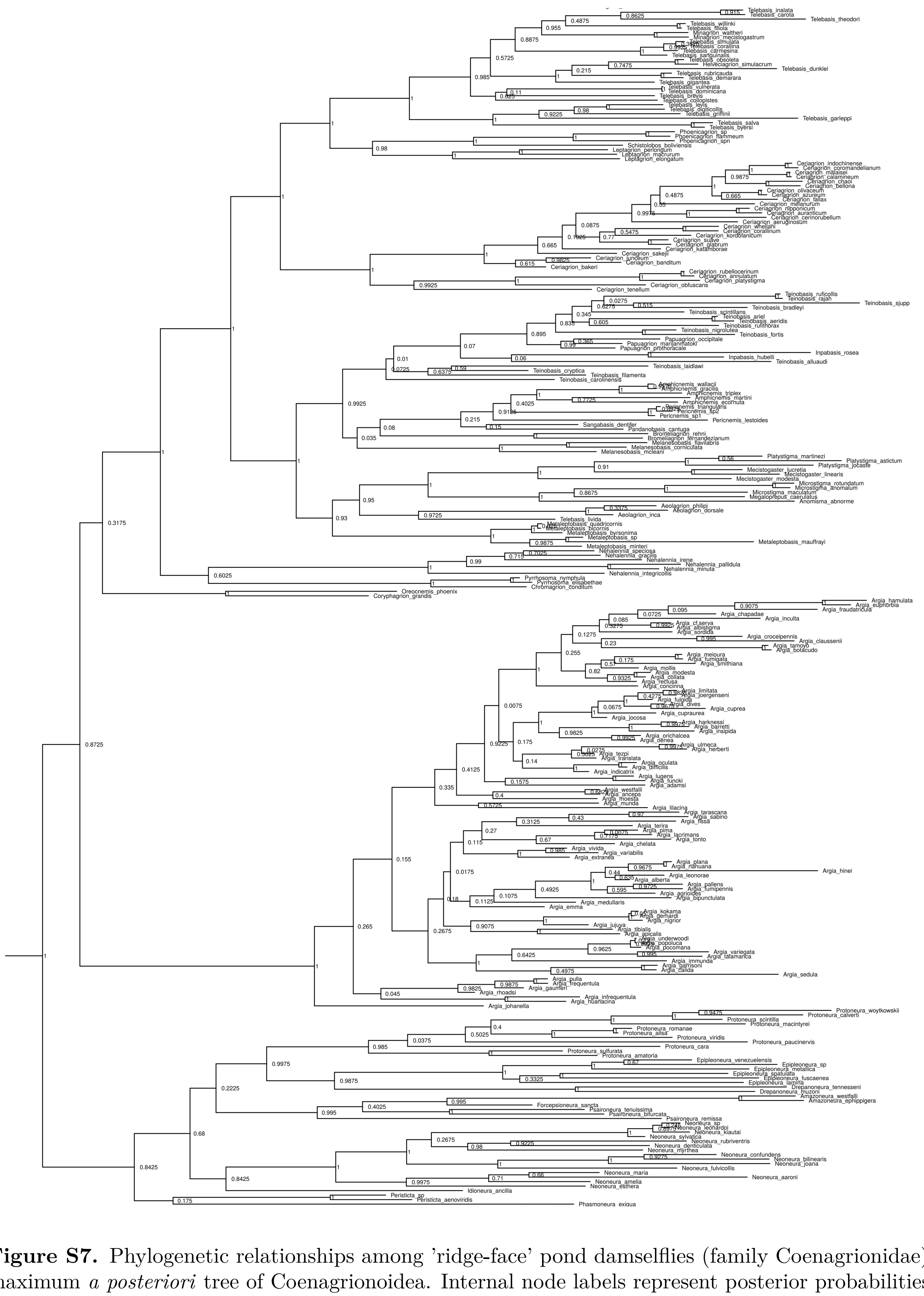

**Figure S7.** Phylogenetic relationships among 'ridge-face' pond damselflies (family Coenagrionidae) in the maximum *a posteriori* tree of Coenagrionoidea. Internal node labels represent posterior probabilities (PP).

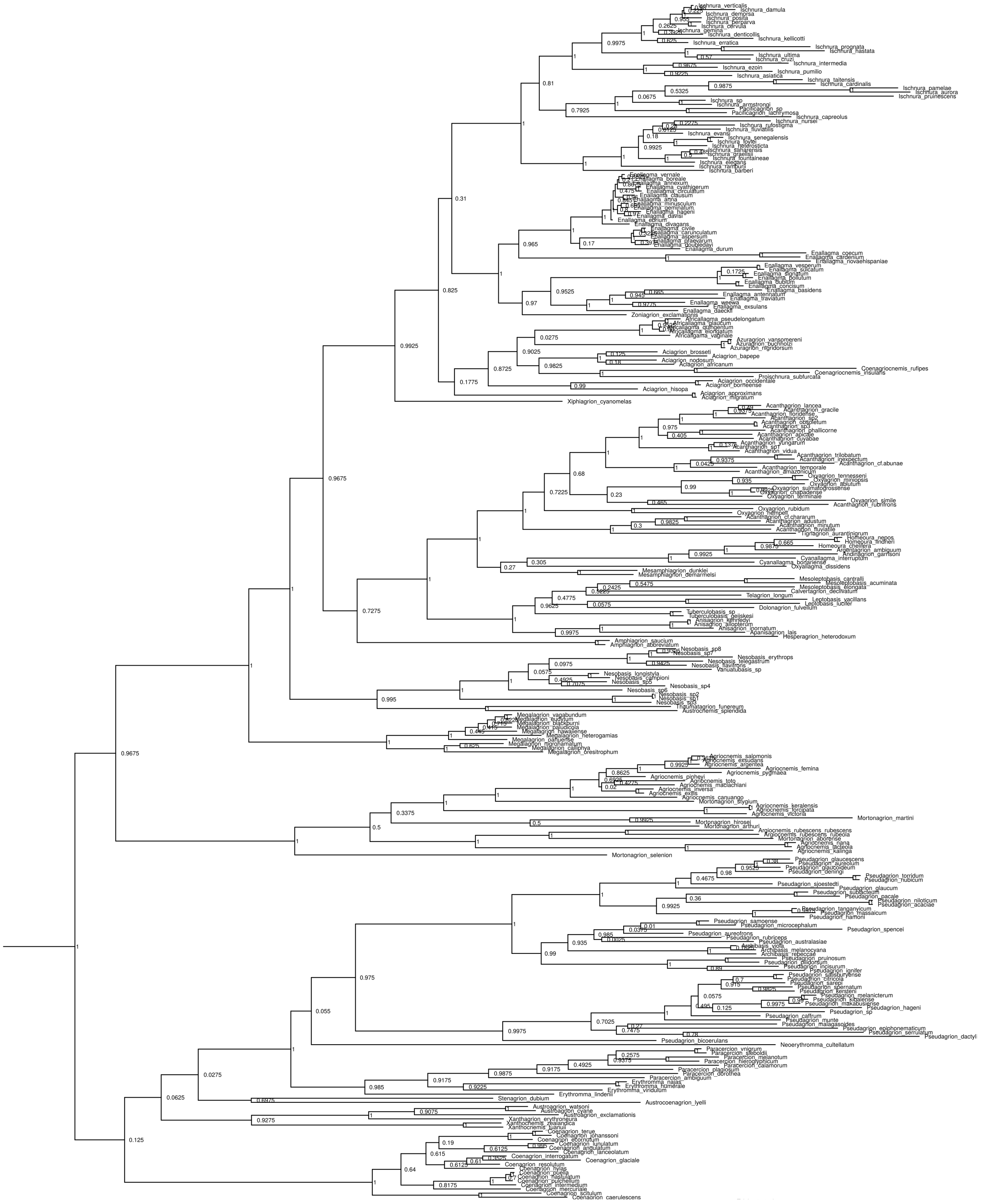

**Figure S8.** Phylogenetic relationships among 'core' pond damselflies (family Coenagrionidae) in the maximum *a posteriori* tree of Coenagrionoidea. Internal node labels represent posterior probabilities (PP).

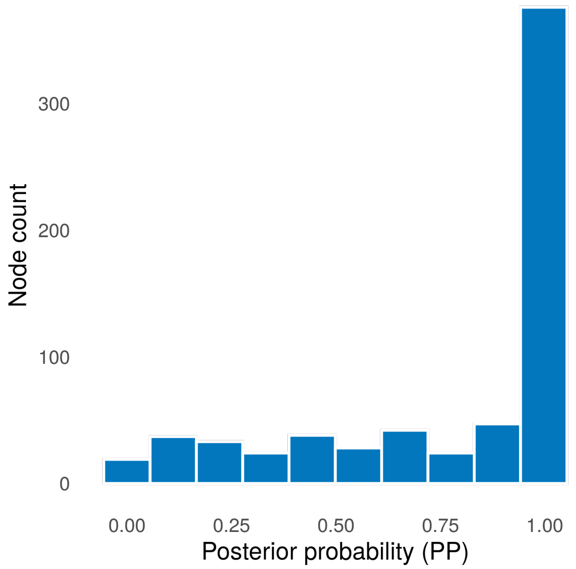

**Figure S9.** Histogram of posterior probabilities (PP) supporting the internal nodes in the maximum *a posteriori* tree of Coenagrionoidea. The phylogeny consists of 669 taxa, and 668 internal nodes. The root node, was constrained to designate the split between featherlegs (family Platycnemididae) and pond damselflies (family Coenagrionidae), based on recent phylogenomic studies.

**Table S9.** Genera in the damselfly superfamily Coenagrionoidea and cladistic implications of the present and previous studies. We studied the two largest families in Coenagrionoidea: the pond damselflies (Coenagrionidae, abbreviated as C) and the featherlegs (Platycnemididae, abbreviated as P).

| Family | Genus | Posterior Probability | Implication of present study | Contributions of previous studies | Congruence with previous studies |
|---|---|---|---|---|---|
| C | *Acanthagrion* | NA | Paraphyletic: Oxyagrion and Tigriagrion nested within | Von Ellenrieder and Lozano (2007) define Acanthagrion and Oxyagrion as closely related but monophyletic based on morphological characters | Inferred relationships challenge morphological studies |
| C | *Aciagrion* | NA | Paraphyletic: Proischnura, Coenagriocnemis, Azuragrion and Africallagma nested within Afrotropical clade | African clade supported in Dijkstra et al. (2014) | Inferred relationships partly supported in previous phylogenetic studies, appropriate level studies are missing |

*(continued)*

| Family | Genus | Posterior Probability | Implication of present study | Contributions of previous studies | Congruence with previous studies |
|---|---|---|---|---|---|
| C | *Aeolagrion* | 1.00 | Monophyletic | Morphological review in Tennessen (2009) | Inferred relationships congruent with morphological studies |
| C | *Africallagma* | 1.00 | Monophyletic but nested within Aciagrion | Monophyly supported in May (2022), placement supported in Dijkstra et al. (2014) | Inferred relationships partly supported in previous phylogenetic studies |
| C | *Agriocnemis* | NA | Paraphyletic: Mortonagrion and Argiocnemis nested within | Paraphyly supported in Dijkstra et al. (2014), Toussaint et al. 2019 | Inferred relationships partly supported in previous phylogenetic studies, appropriate level studies are missing |
| C | *Amazoneura* | 1.00 | Monophyletic | Morphological review in Machado (2004) | Inferred relationships congruent with morphological studies |
| C | *Amphiagrion* | 1.00 | Monophyletic | NA | Appropriate level studies are missing |
| C | *Amphicnemis* | 1.00 | Monophyletic | Morphological review in Dow et al. (2010) and Villanueva (2012) | Inferred relationships congruent with morphological studies |
| C | *Andinagrion* | NA | Single species sampled | NA | NA |
| C | *Anisagrion* | 1.00 | Monophyletic | NA | Appropriate level studies are missing |
| C | *Apanisagrion* | NA | Monotypic | NA | NA |
| C | *Archibasis* | 1.00 | Monophyletic but nested within Pseudagrion | Placement supported in Dijkstra et al. (2014) | Inferred relationships partly supported in previous phylogenetic studies, appropriate level studies are missing |
| C | *Argentagrion* | NA | Monotypic but nested within Cyanallagma | Morphological review in Von Ellenrieder (2008a) | NA |

*(continued)*

| Family | Genus | Posterior Probability | Implication of present study | Contributions of previous studies | Congruence with previous studies |
|---|---|---|---|---|---|
| C | *Argia* | 1.00 | Monophyletic | Reviewed in Caesar and Wenzel (2009) and Torres-Pachón et al. (2017) | Inferred relationships congruent with morphological studies |
| C | *Argiocnemis* | NA | Single species sampled but nested within Agriocnemis | Placement supported in Dijkstra et al. (2014) | Inferred relationships partly supported in previous phylogenetic studies, appropriate level studies are missing |
| C | *Austroagrion* | 0.90 | Monophyletic | NA | Appropriate level studies are missing |
| C | *Austrocoenagrion* | NA | Monotypic | NA | NA |
| C | *Azuragrion* | 1.00 | Monophyletic but nested within Aciagrion | Monophyly supported in May (2022), placement supported in Dijkstra et al. (2014) | Inferred relationships partly supported in previous phylogenetic studies |
| C | *Bromeliagrion* | 1.00 | Monophyletic | Morphological review in De Marmels and Garrison (2005) | Inferred relationships congruent with morphological studies |
| C | *Calvertagrion* | NA | Single species sampled | Morphological review in Tennessen (2015) | NA |
| C | *Ceriagrion* | 1.00 | Monophyletic | Monophyly supported in Dijkstra et al. (2014), Toussaint et al. (2019) | Inferred relationships supported in previous phylogenetic studies, appropriate level studies are missing |
| C | *Chromagrion* | NA | Monotypic | Morphological review in De Marmels (2002) | NA |
| C | *Coenagriocnemis* | 1.00 | Monophyletic but nested within Aciagrion | NA | Appropriate level studies are missing |
| C | *Coenagrion* | 1.00 | Monophyletic | Reviewed in Swaegers et al. (2014) | Inferred relationships congruent with morphological studies |

*(continued)*

| Family | Genus | Posterior Probability | Implication of present study | Contributions of previous studies | Congruence with previous studies |
|---|---|---|---|---|---|
| C | *Coryphagrion* | NA | Monotypic | NA | NA |
| C | *Cyanallagma* | NA | Paraphyletic: Andinagrion, Argentagrion and Homeoura nested within | Morphological review in von Ellenrieder and Garrison (2008a) | Inferred relationships challenge morphological studies |
| C | *Dolonagrion* | NA | Monotypic | Morphological review in Garrison and von Ellenrieder (2008) | NA |
| C | *Drepanoneura* | 1.00 | Monophyletic | Morphological review in von Ellenrieder and Garrison (2008b) | Inferred relationships congruent with morphological studies |
| C | *Enallagma* | NA | Paraphyletic: Zoniagrion nested within | No previous study considers Enallagma and Zoniagrion. Latest review in Callahan and McPeek (2016) | Inferred relationships partly supported in previous phylogenetic studies, appropriate level studies are missing |
| C | *Epipleoneura* | 1.00 | Monophyletic | Monophyly supported in Pessacq (2008), Morphological review in Pessacq (2014) | Inferred relationships congruent with morphological studies |
| C | *Erythromma* | NA | Paraphyletic: Paracercion nested within | Different relationships supported in Weekers and Dumont (2004) | Inferred relationships challenge previous phylogenetic studies |
| C | *Forcepsioneura* | NA | Single species sampled but nested within Psaironeura | Monophyly supported in Pimenta et al. (2019), close relationship with Psaironeura supported in Pessacq (2008) | Inferred relationships partly supported in previous phylogenetic studies, appropriate level studies are missing |
| C | *Hesperagrion* | NA | Single species sampled | Morphological review in De Marmels (2002) | NA |

*(continued)*

| Family | Genus | Posterior Probability | Implication of present study | Contributions of previous studies | Congruence with previous studies |
|---|---|---|---|---|---|
| C | *Homeoura* | 0.66 | Monophyletic | Morphological review in Von Ellenrieder (2008) | Inferred relationships congruent with morphological studies |
| C | *Idioneura* | NA | Single species sampled | NA | Appropriate level studies are missing |
| C | *Inpabasis* | 1.00 | Monophyletic but nested within Teinobasis | Different relationships supported in Dijkstra et al. (2014) | Inferred relationships challenge previous phylogenetic studies |
| C | *Ischnura* | NA | Paraphyletic: Pacificagrion nested within | Paraphyly supported in Blow et al. (2021) | Inferred relationships congruent with morphological studies |
| C | *Leptagrion* | 1.00 | Monophyletic | Morphological review in De Marmels and Garrison (2005) | Inferred relationships congruent with morphological studies |
| C | *Leptobasis* | 1.00 | Monophyletic | Morphological review in Garrison and von Ellenrieder (2010) | Inferred relationships congruent with morphological studies |
| C | *Mecistogaster* | NA | Paraphyletic: Platystigma nested within | Paraphyly supported in Garrison et al. (2010), Ingley et al. (2012) and Toussaint el al. (2019), monophyly supported in Machado et al. (2017) | Inferred relationships congruent with phylogenetic studies and some morphological studies |
| C | *Megalagrion* | 1.00 | Monophyletic | Reviewed in Jordan et al. (2003) | Inferred relationships congruent with morphological studies |
| C | *Megaloprepus* | NA | Single species sampled | Placement supported in Ingley et al. (2012) and Toussaint et al. (2019) | Inferred relationships congruent with morphological studies |

*(continued)*

| Family | Genus | Posterior Probability | Implication of present study | Contributions of previous studies | Congruence with previous studies |
|---|---|---|---|---|---|
| C | *Melanesobasis* | 1.00 | Monophyletic | Reviewed in Beatty et al. (2017) | Inferred relationships congruent with morphological studies |
| C | *Mesamphiagrion* | 1.00 | Monophyletic | Morphological review in von Ellenrieder and Garrison (2008a) | Inferred relationships congruent with morphological studies |
| C | *Mesoleptobasis* | 0.55 | Monophyletic | Morphological review in Garrison and von Ellenrieder (2009) | Inferred relationships congruent with morphological studies |
| C | *Metaleptobasis* | 1.00 | Monophyletic | Morphological review in von Ellenrieder (2013) | Inferred relationships congruent with morphological studies |
| C | *Microstigma* | 1.00 | Monophyletic | Monophyly supported in Ingley et al. (2012) and Toussaint et al. (2019) | Inferred relationships congruent with morphological studies |
| C | *Minagrion* | 1.00 | Monophyletic but nested within Telebasis | Morphological review in Vilela et al. (2020) | Inferred relationships challenge morphological studies |
| C | *Mortonagrion* | NA | Paraphyletic: Agriocnemis nested within | Placement supported in Dijkstra et al. (2014) | Inferred relationships supported in previous phylogenetic studies, appropriate level studies are missing |
| C | *Nehalennia* | 1.00 | Monophyletic | Monophyly supported in De Marmels (1984) | Inferred relationships congruent with morphological studies |
| C | *Neoerythromma* | NA | Single species sampled | NA | Appropriate level studies are missing |

*(continued)*

| Family | Genus | Posterior Probability | Implication of present study | Contributions of previous studies | Congruence with previous studies |
|---|---|---|---|---|---|
| C | *Neoneura* | 1.00 | Monophyletic | Monophyly supported in Pessacq (2008) and Toussaint et al. (2019) | Inferred relationships congruent with morphological studies |
| C | *Nesobasis* | NA | Paraphyletic: Vanuatubasis nested within | Paraphyly supported in Ferguson et al. (2023) | Inferred relationships congruent with morphological studies |
| C | *Oreocnemis* | NA | Monotypic | Placement supported in Toussaint et al. (2019) | Inferred relationships supported in previous phylogenetic studies, appropriate level studies are missing |
| C | *Oxyagrion* | NA | Paraphyletic: Acanthagrion nested within | Von Ellenrieder and Lozano (2007) define Acanthagrion and Oxyagrion as closely related but monophyletic based on morphological characters | Inferred relationships challenge morphological studies |
| C | *Oxyallagma* | NA | Single species sampled | NA | Appropriate level studies are missing |
| C | *Pacificagrion* | 1.00 | Monophyletic but nested within Ischnura | Placement supported in Blow et al. (2021) | Inferred relationships congruent with morphological studies |
| C | *Pandanobasis* | NA | Single species sampled | Morphological review in Villanueva (2012) | Inferred relationships congruent with morphological studies |
| C | *Papuagrion* | 0.99 | Monophyletic but nested within Teinobasis | Placement supported in Toussaint et al. (2019) | Inferred relationships supported in previous phylogenetic studies, appropriate level studies are missing |

*(continued)*

| Family | Genus | Posterior Probability | Implication of present study | Contributions of previous studies | Congruence with previous studies |
|---|---|---|---|---|---|
| C | *Paracercion* | 0.99 | Monophyletic but nested within Erythromma | Different relationships supported in Weekers and Dumont (2004) | Inferred relationships challenge previous phylogenetic studies |
| C | *Pericnemis* | 1.00 | Monophyletic | Morphological review in Villanueva (2012) | Inferred relationships congruent with morphological studies |
| C | *Peristicta* | 1.00 | Monophyletic | Monophyly supported in Pessacq (2008) | Inferred relationships congruent with morphological studies |
| C | *Phasmoneura* | NA | Single species sampled | Morphological review in Lencioni (1999), reviewed in Pessacq (2008) | NA |
| C | *Phoenicagrion* | 1.00 | Monophyletic | Morphological review in Von Ellenrieder (2008b) | Inferred relationships congruent with morphological studies |
| C | *Platystigma* | 1.00 | Monophyletic but nested within Mecistogaster | Placement supported in Ingley et al. (2012) and Toussaint el al. (2019), not supported in Machado et al. (2017) | Inferred relationships congruent with phylogenetic studies and some morphological studies |
| C | *Proischnura* | NA | Single species sampled but nested within Aciagrion | NA | Appropriate level studies are missing |
| C | *Protoneura* | 0.98 | Monophyletic | Monophyly supported in Pessacq (2008) | Inferred relationships congruent with morphological studies |
| C | *Psaironeura* | NA | Paraphyletic: Forcepsioneura nested within | Relationship to Forcepsioneura supported in Pessacq (2008) | Inferred relationships supported in previous phylogenetic studies, appropriate level studies are missing |

*(continued)*

| Family | Genus | Posterior Probability | Implication of present study | Contributions of previous studies | Congruence with previous studies |
|---|---|---|---|---|---|
| C | *Pseudagrion* | NA | Paraphyletic: Archibasis nested within | Paraphyly supported in Dijkstra et al. (2014), reviewed in Dijkstra et al. (2007) | Inferred relationships congruent with morphological studies |
| C | *Pyrrhosoma* | 1.00 | Monophyletic | Reviewed in Guan et al. (2013) | Inferred relationships congruent with morphological studies |
| C | *Sangabasis* | NA | Single species sampled | Morphological review in Villanueva (2012) | Inferred relationships congruent with morphological studies |
| C | *Schistolobos* | NA | Monotypic | Morphological review in von Ellenrieder and Garrison (2008c) | Inferred relationships congruent with morphological studies |
| C | *Stenagrion* | NA | Single species sampled | NA | Appropriate level studies are missing |
| C | *Teinobasis* | NA | Paraphyletic: Papuagrion nested within | Paraphyly supported in Toussaint et al. (2019) | Inferred relationships supported in previous phylogenetic studies, appropriate level studies are missing |
| C | *Telagrion* | NA | Single species sampled | Morphological review in von Ellenrieder and Garrison (2008c) | Appropriate level studies are missing |
| C | *Telebasis* | 1.00 | Paraphyletic: Minagrion nested within | Morphological review in Garrison (2008) | Appropriate level studies are missing |
| C | *Thaumatagrion* | NA | Monotypic | NA | NA |
| C | *Tigriagrion* | NA | Monotypic but nested within Acanthagrion | NA | Appropriate level studies are missing |
| C | *Tuberculobasis* | 1.00 | Monophyletic | Morphological review in Machado (2009) | Inferred relationships congruent with morphological studies |

*(continued)*

| Family | Genus | Posterior Probability | Implication of present study | Contributions of previous studies | Congruence with previous studies |
|---|---|---|---|---|---|
| C | *Vanuatubasis* | NA | Single species sampled but nested within Nesobasis | Placement supported in Ferguson et al. (2023), Morphological review in Saxton et al. (2022) | Inferred relationships congruent with morphological studies |
| C | *Xanthagrion* | NA | Monotypic | NA | NA |
| C | *Xanthocnemis* | 1.00 | Monophyletic | Reviewed in Marinov et al. (2016) | Inferred relationships congruent with morphological studies |
| C | *Xiphiagrion* | NA | Single species sampled | NA | Appropriate level studies are missing |
| C | *Zoniagrion* | NA | Monotypic but nested within Enallagma | Morphological review in De Marmels (2002) | Appropriate level studies are missing |
| P | *Allocnemis* | 1.00 | Monophyletic | Monophyly supported in Dijkstra et al. (2014) | Inferred relationships partly supported in previous phylogenetic studies, appropriate level studies are missing |
| P | *Arabicnemis* | NA | Monotypic | NA | NA |
| P | *Calicnemia* | 1.00 | Monophyletic but nested within Coeliccia | NA | Appropriate level studies are missing |
| P | *Coeliccia* | NA | Paraphyletic: Calicnemia and Indocnemis nested within | Paraphyly supported in Dijkstra et al. (2014) | Inferred relationships partly supported in previous phylogenetic studies, appropriate level studies are missing |
| P | *Copera* | 1.00 | Monophyletic | Reviewed in Lim et al. (2013) | Inferred relationships congruent with morphological studies |
| P | *Cyanocnemis* | NA | Monotypic | NA | NA |

*(continued)*

| Family | Genus | Posterior Probability | Implication of present study | Contributions of previous studies | Congruence with previous studies |
|---|---|---|---|---|---|
| P | *Elattoneura* | NA | Paraphyletic: Nosostict annd Prodasineura nested within | Paraphyly supported in Dijkstra et al. (2014) | Inferred relationships supported in previous phylogenetic studies, appropriate level studies are missing |
| P | *Esme* | NA | Single species sampled | NA | NA |
| P | *Idiocnemis* | NA | Single species sampled | Morphological review in Gassmann (2005) | NA |
| P | *Igneocnemis* | 1.00 | Monophyletic | Morphological review in Gassmann (2005) | Inferred relationships congruent with morphological studies |
| P | *Indocnemis* | NA | Paraphyletic: Coeliccia nested within | Paraphyly supported in Dijkstra et al. (2014) | Inferred relationships supported in previous phylogenetic studies, appropriate level studies are missing |
| P | *Lieftinckia* | NA | Single species sampled | Morphological review in Gassmann (2005) | NA |
| P | *Lochmaeocnemis* | NA | Monotypic | Morphological review in Gassmann (2005) | NA |
| P | *Matticnemis* | NA | Monotypic but nested within Platycnemis | Morphological review in Djkstra (2013), Placement as sister of Platycnemis in Dijkstra et al. (2014) | Inferred relationships challenge previous phylogenetic studies |
| P | *Mesocnemis* | 1.00 | Monophyletic | Monophyly supported in Dijkstra et al. (2014) | Inferred relationships supported in previous phylogenetic studies, appropriate level studies are missing |

*(continued)*

| Family | Genus | Posterior Probability | Implication of present study | Contributions of previous studies | Congruence with previous studies |
|---|---|---|---|---|---|
| P | *Metacnemis* | NA | Monotypic | Morphological review in Djkstra (2013) | Inferred relationships congruent with morphological studies |
| P | *Nososticta* | 1.00 | Monophyletic but nested within Elattoneura | Paraphyly supported in Dijkstra et al. (2014) | Inferred relationships supported in previous phylogenetic studies, appropriate level studies are missing |
| P | *Onychargia* | NA | Single species sampled | NA | Appropriate level studies are missing |
| P | *Palaiargia* | NA | Single species sampled | NA | Appropriate level studies are missing |
| P | *Paracnemis* | NA | Single species sampled | NA | Appropriate level studies are missing |
| P | *Paramecocnemis* | 1.00 | Monophyletic | Morphological review in Orr et al. (2012) | Inferred relationships congruent with morphological studies |
| P | *Phylloneura* | NA | Monotypic | NA | NA |
| P | *Platycnemis* | NA | Paraphyletic: Matticnemis and Pseudocopera nested within | Different relationships supported in Dijkstra et al. (2014) | Inferred relationships challenge previous phylogenetic studies |
| P | *Prodasineura* | NA | Paraphyletic: Elattoneura nested within | Paraphyly supported in Dijkstra et al. (2014) | Inferred relationships supported in previous phylogenetic studies, appropriate level studies are missing |
| P | *Proplatycnemis* | 1.00 | Monophyletic | NA | Appropriate level studies are missing |
| P | *Pseudocopera* | 1.00 | Monophyletic but nested within Platycnemis | Placement supported in Dijkstra et al. (2014) | Inferred relationships supported in previous phylogenetic studies, appropriate level studies are missing |

*(continued)*

| Family | Genus | Posterior Probability | Implication of present study | Contributions of previous studies | Congruence with previous studies |
|---|---|---|---|---|---|
| P | *Risiocnemis* | 1.00 | Monophyletic | Morphological review in Gassmann (2005) | Inferred relationships congruent with morphological studies |
| P | *Salomocnemis* | NA | Monotypic | NA | NA |
| P | *Spesbona* | NA | Monotypic | Morphological review in Djkstra (2013) | Inferred relationships congruent with morphological studies |
| P | *Stenocnemis* | NA | Monotypic | NA | NA |
| P | *Torrenticnemis* | NA | Monotypic | NA | NA |

##### ***Ischnura* and *Pacificagrion***

The South Pacific species of the cosmopolitan genus *Ischnura* and the genus *Pacificagrion* form a strongly supported clade (PP = 1.00, Fig. S10). Support for this clade was also found by Blow et al. (2021), who used an overlapping data set but a different tree prior in their dated phylogeny of *Ischnura*, and by mitochondrial markers in Karube et al. (2012). While relationships within the South Pacific clade are not fully resolved, current evidence supports the inclusion of *Pacificagrion* within *Ischnura*.

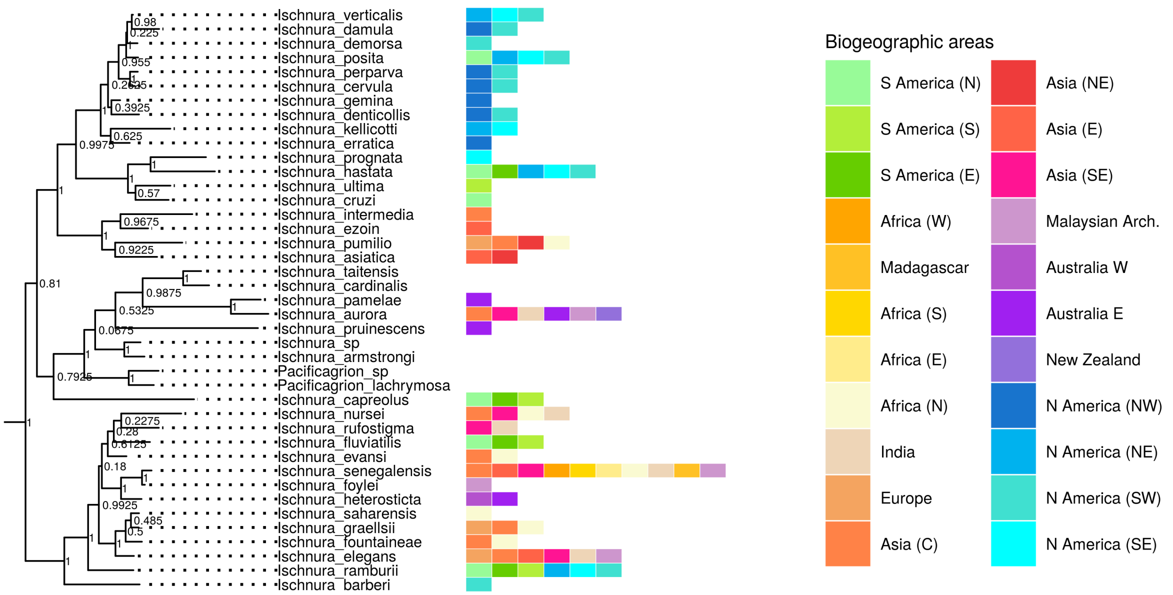

**Figure S10.** View of the clade containing the genera *Ischnura*, and *Pacificagrion* in the maximum *a posteriori* tree of Coenagrionoidea. Internal node labels represent posterior probabilities (PP). Taxa with missing distribution data are endemic to Pacific Islands.

##### ***Enallagma* and *Zoniagrion***

The genus *Enallagma* consists of about 47 species distributed across North America, with a few taxa reaching into the Neotropics and the Palearctic. *Zoniagrion* is a monotypic genus found in the Western Coast of North America. Here, we found strong support (PP = 0.97) for a clade comprising *Zoniagrion* and the 'Southern clade' of *Enallagma* (Fig. S11). The 'Southern clade' of *Enallagma* was also recovered in Callahan and McPeek (2016) and Turgeon et al. (2005). However, these previous molecular phylogenetic studies did not incorporate *Zoniagrion* in their taxonomic sampling. The latest morphological revision of *Zoniagrion*, identified shared characters with *Enallagma* and *Acanthagrion*, but concluded that the genus either shared a more recent common ancestor with *Acanthagrion*, or diverged from the common ancestor of both *Enallagma* and *Acanthagrion* (De Marmels 2002). Due to the possibility of a missidentified specimen of *Z. exclamationis*, additional genetic data should be generated to confirm or refute the present finding.

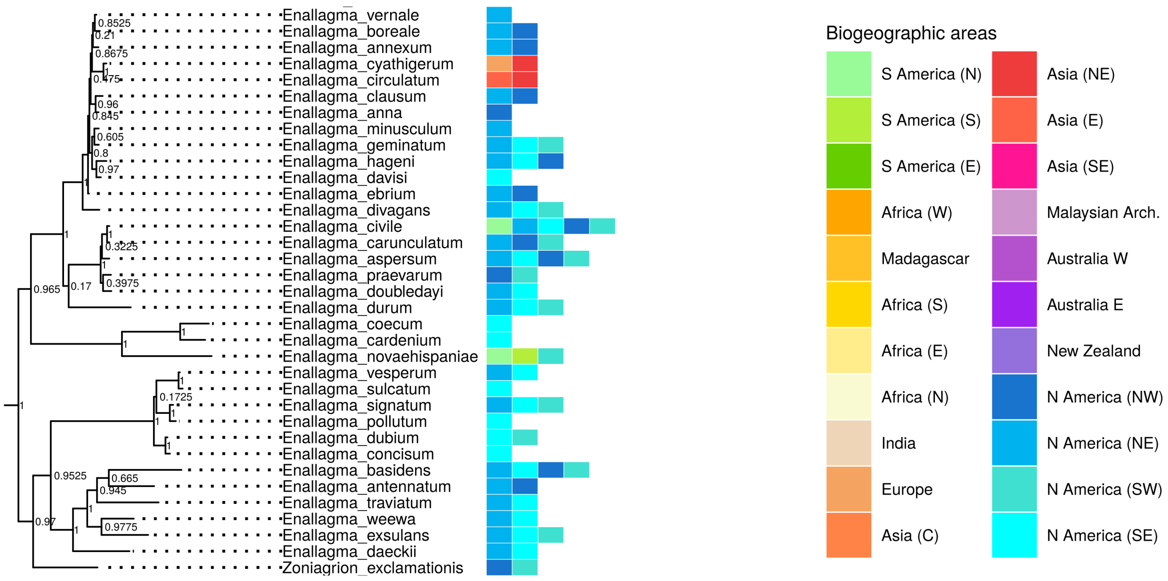

**Figure S11.** View of the clade containing the genera *Enallagma*, and *Zoniagrion* in the maximum *a posteriori* tree of Coenagrionoidea. Internal node labels represent posterior probabilities (PP).

##### ***Aciagrion*, *Proischnura*, *Coenagriocnemis*, *Africallagma* and *Azuragrion***

Our phylogenetic analysis did not recover the Paelotropical *Aciagrion* as a monophyletic clade (Fig. S12). Instead, the African representatives of the genus were more closely related to other African genera (*Proischnura*, *Coenagriocnemis*, *Africallagma* and *Azuragrion*), with relatively strong support (PP = 0.90). Within this African clade, monophyly of each genera was strongly supported (all PP = 1.00). Three of the remaining *Aciagrion* (*A. occidentale*, *A. hisopa* and *A. borneense*) are distributed across India, Southeast Asia, and the Malaysian Archipelago and form a strongly supported clade (PP = 0.99), sister to the African clade described above. Finally the two remaining taxa in the present study (*A. migratum*, *A. approximans*) form a clade that might be sister to all other *Aciagrion*, although this relationship is weakly supported (PP = 0.17). We are not aware of any other study testing the monophyly of *Aciagrion*, and therefore a phylogenetic and taxonomic revision of the clade is warranted.

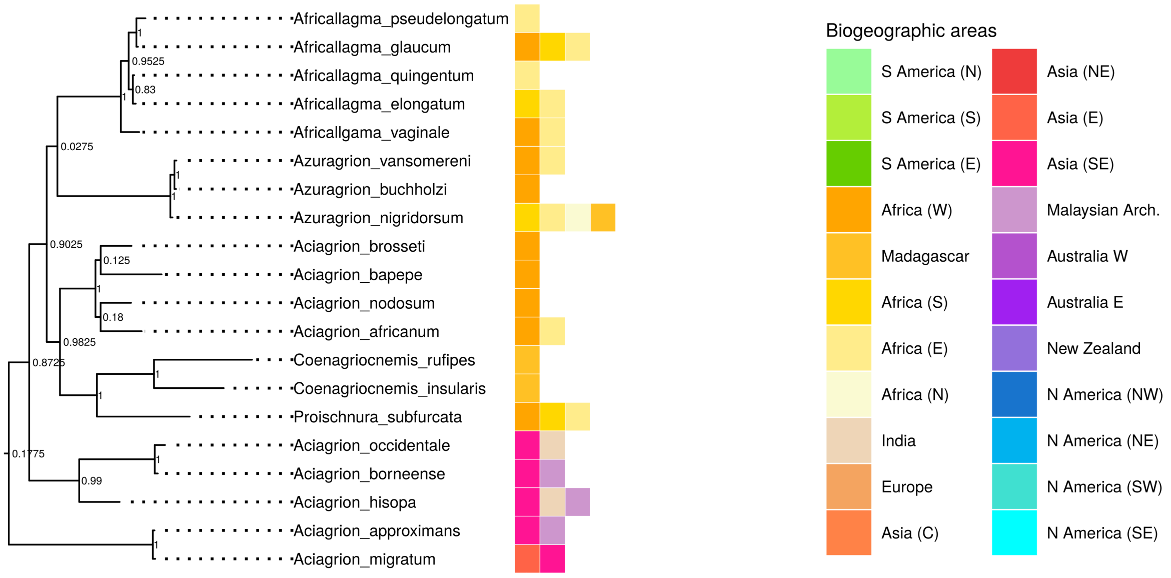

**Figure S12.** View of the clade containing the genera *Aciagrion*, *Agricallagma*, *Azuragrion*, *Coenagriocnemis*, and *Proischnura* in the maximum *a posteriori* tree of Coenagrionoidea. Internal node labels represent posterior probabilities (PP).

##### ***Acanthatrion*, *Oxyagrion*, and *Tigriagrion***

The close affinity between *Acanthagrion* and *Oxyagrion* was established already by Selys(1876), upon description of the genera. Most recently, von Ellenrieder and Lozano (2008) redefined the two genera based on morphological traits and reassigned species according to their classification. Our results strongly support the monophyly of a clade containing *Acanthagrion*, *Oxyagrion* and the monotypic *Tigriagrion* (Posterior probability = 1.00), in which neither *Acanthagrion* nor *Oxyagrion* are monophyletic (Fig S13). Nonetheless, support for clades composed of species in both genera varies from intermediate (PP = 0.72) to low (PP = 0.23). Support for the placement of *Tigriagrion* within *Acanthagrion* is instead strong (PP = 1.00). Our study included nearly half of all *Acanthagrion* species and a third of *Oxyagrion*. Yet fully resolving the relationships in this clade will likely require more extensive sampling.

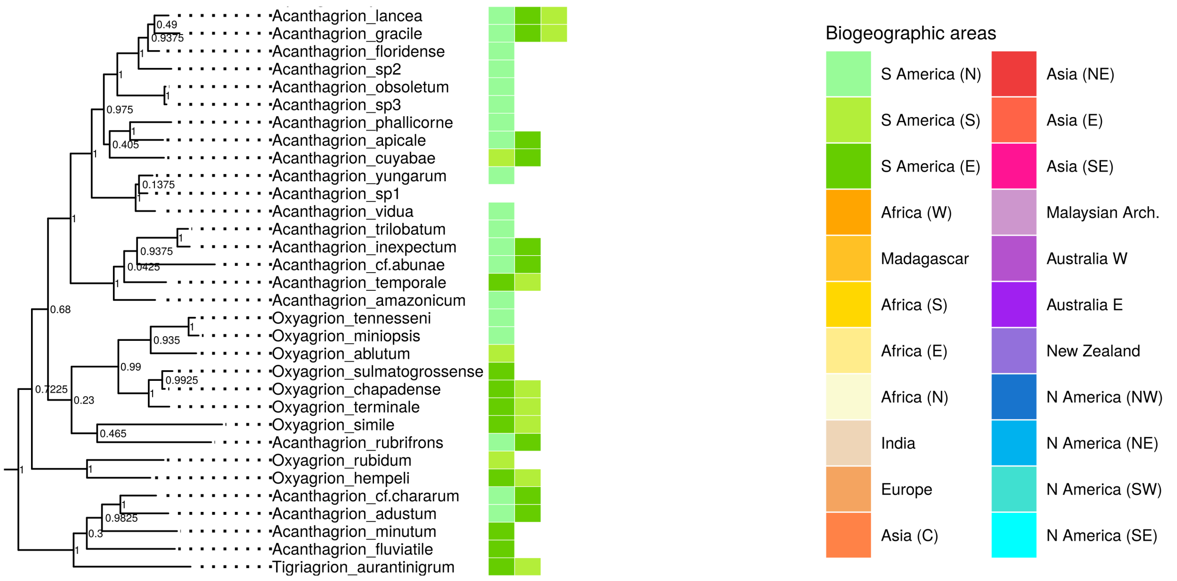

**Figure S13.** View of the clade containing the genera *Acanthagrion*, *Oxyagrion*, and *Tigriagrion* in the maximum *a posteriori* tree of Coenagrionoidea. Internal node labels represent posterior probabilities (PP).

##### ***Cyanallagma*, *Andinagrion*, *Argentagrion*, and *Homeoura***

The three South American genera *Andinagrion*, *Argentagrion*, and *Homeoura* form a strongly supported clade (PP = 1.00). Monophyly of the genus *Homeoura* has intermediate support (PP = 0.66), while the other two genera are either monotypic (*Argentagrion*), or represented by a single species in this study (*Andinagrion*). Our phylogenetic inference strongly suggests that the genus *Cyanallagma* is paraphyletic, with *C. interruptum* being a sister taxon to a clade containing the three genera mentioned above, and *C. bonariense* having a more distant common ancestor (Fig. S14). Von Ellenrieder and Garrison (2008a) conducted a thorough morphological revision of *Cyanallagma*, but noted that the genus lacks unique characters and is rather diagnosed by a combination of characters, some shared with other genera. Moreover, Von Ellenrieder and Garrison (2008a) focused on contrasting *Cyanallagma* to *Mesamphiagrion* and *Oreiallagma* (not included in the present study), and thus specimens of *Andinagrion*, *Argentagrion*, and *Homeoura* were not examined. We thus conclude that a systematic study encompassing the six genera, as well as *Oxyallagma*, is warranted.

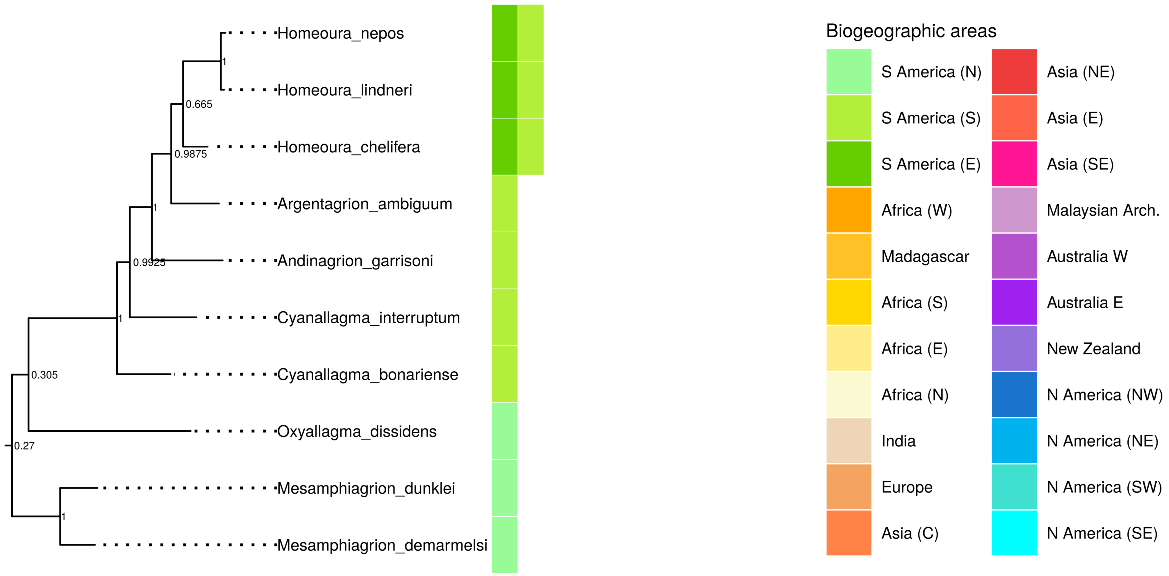

**Figure S14.** View of the clade containing the genera *Andinagrion*, *Argentagrion*, *Cyanallagma*, *Homeoura*, and *Mesamphiagrion* in the maximum *a posteriori* tree of Coenagrionoidea. Internal node labels represent posterior probabilities (PP).

##### *Nesobasis* and *Vanuatubasis*

We found an undescribed species of *Vanuatubasis* (collected in Vanuatu) to be nested within *Nesobasis* (from Fiji), although the exact placement of this species within *Nesobasis* remained uncertain (Fig. S15). Nonetheless, the paraphyly of *Nesobasis* was also supported in a recent phylogenetic study focusing specifically on the relationship between the two genera (Ferguson et al. 2023). Similarly to the present analysis, Ferguson et al. (2023) found *Vanuatubasis* to be closely related to the *longystila* and *erythrops* species groups of *Nesobasis*.

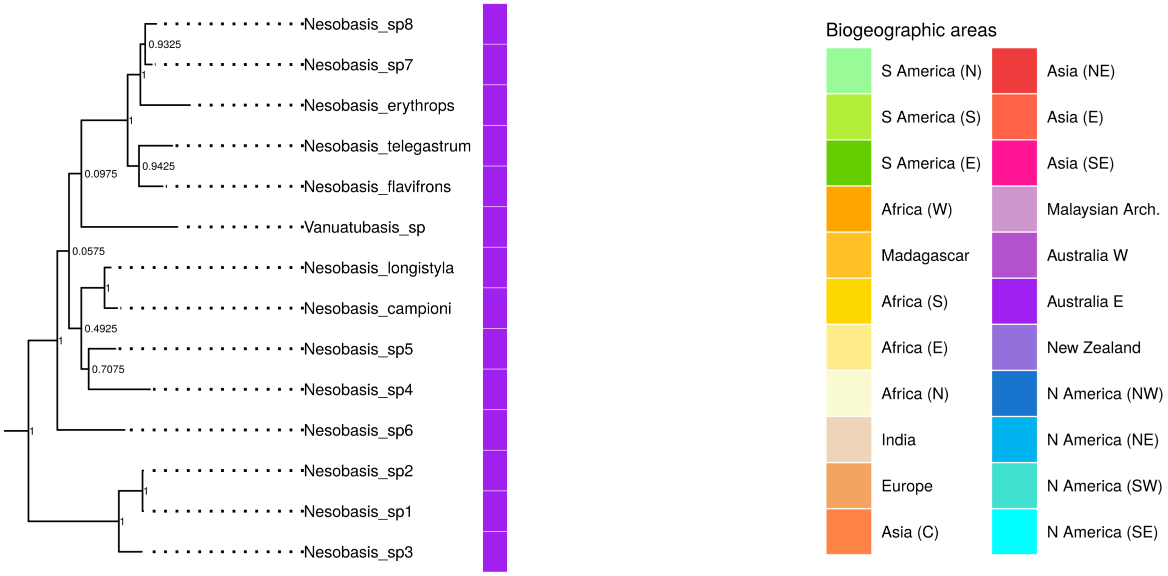

**Figure S15.** View of the clade containing the genera *Nesobasis* and *Vanuatubasis* in the maximum *a posteriori* tree of Coenagrionoidea. Internal node labels represent posterior probabilities (PP).

##### ***Agriocnemis*, *Argiocnemis* and *Mortonagrion***

The genus *Agriocnemis* has a Paleotropical distribution, ranging from Africa to Eastern Australia. A well supported clade (PP = 1.00) includes all African taxa, a strongly supported subclade of Australasian taxa (PP = 1.00), and the only African species attributed to the genus *Mortonagrion* (*M. stygium*) (Fig. S16). This close relationship between *M. stygium* and African *Agriocnemis* was also found in Dijkstra et al. (2014), which treated the taxon as *Agriocnemis stygia*, and in Toussaint et al. (2019). Three Asian species of *Agriocnemis* form another well-supported clade with the Australasian genus *Argiocnemis*, and the Southeast Asian *M. aborense* (PP = 1.00, Fig. S16). Finally, we sampled four more species of *Mortonagrion*, primarily from Southeast and East Asia, which are also closely related to *Agriocnemis* and *Argiocnemis* but their exact placement within the clade is uncertain (Fig. S16). Previous work had already suggested that some of these *Mortonagrion* taxa contribute to the paraphyly of *Agriocnemis* (Dijkstra et al. 2014). However, to our knowledge, there are no densely sampled phylogenetic studies looking specifically into the relationships among the approximately 60 species in this clade. We thus echo previous calls for a systematic revision of the three genera (Dijkstra et al. 2014).

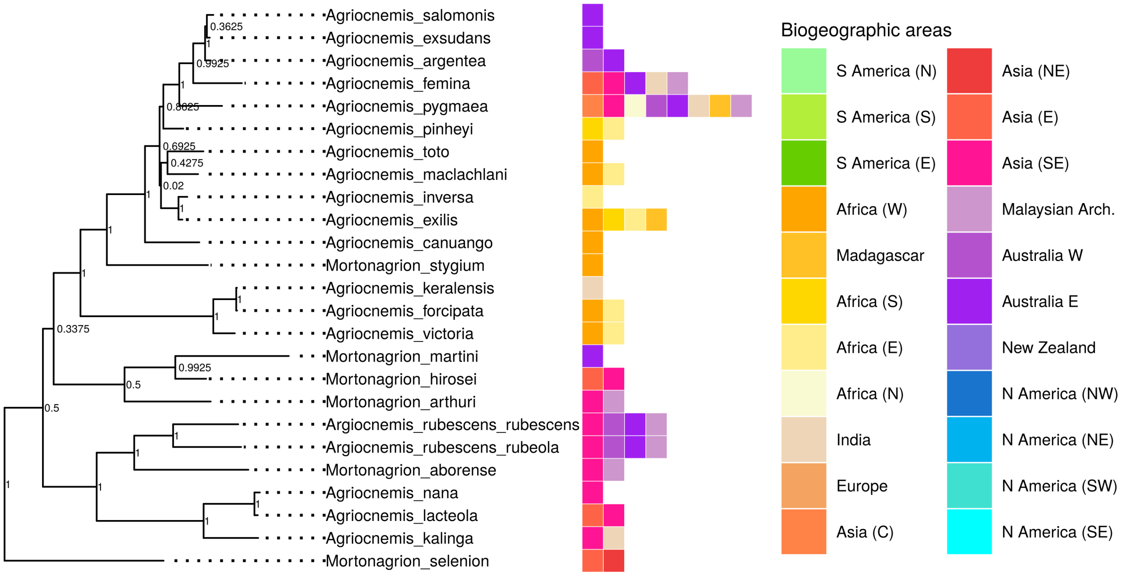

**Figure S16.** View of the clade containing the genera *Agriocnemis*, *Argiocnemis*, and *Mortonagrion* in the maximum *a posteriori* tree of Coenagrionoidea. Internal node labels represent posterior probabilities (PP).

##### ***Pseudagrion* and *Archibasis***

With over 150 species, *Pseudagrion* is the largest genus of Coenagrionoidea. It is distributed across the entire Paleotropics and temperate regions of East Asia. In contrast, *Archibasis*, with approximately nine species, is restricted to Southeast Asia and the Malaysian Archipelago. Our phylogenetic inference uncovered a well-supported clade (PP = 0.99) consisting of *Archibasis* and the Australasian representatives of *Pseudagrion* (Fig. S17). A similar finding was obtained by Dijkstra et al. (2014) under more limited taxonomic sampling. We thus suggest a taxonomic revision might be warranted, either reclassifying part of *Pseudagrion* or subsuming *Archibasis* into *Pseudagrion*.

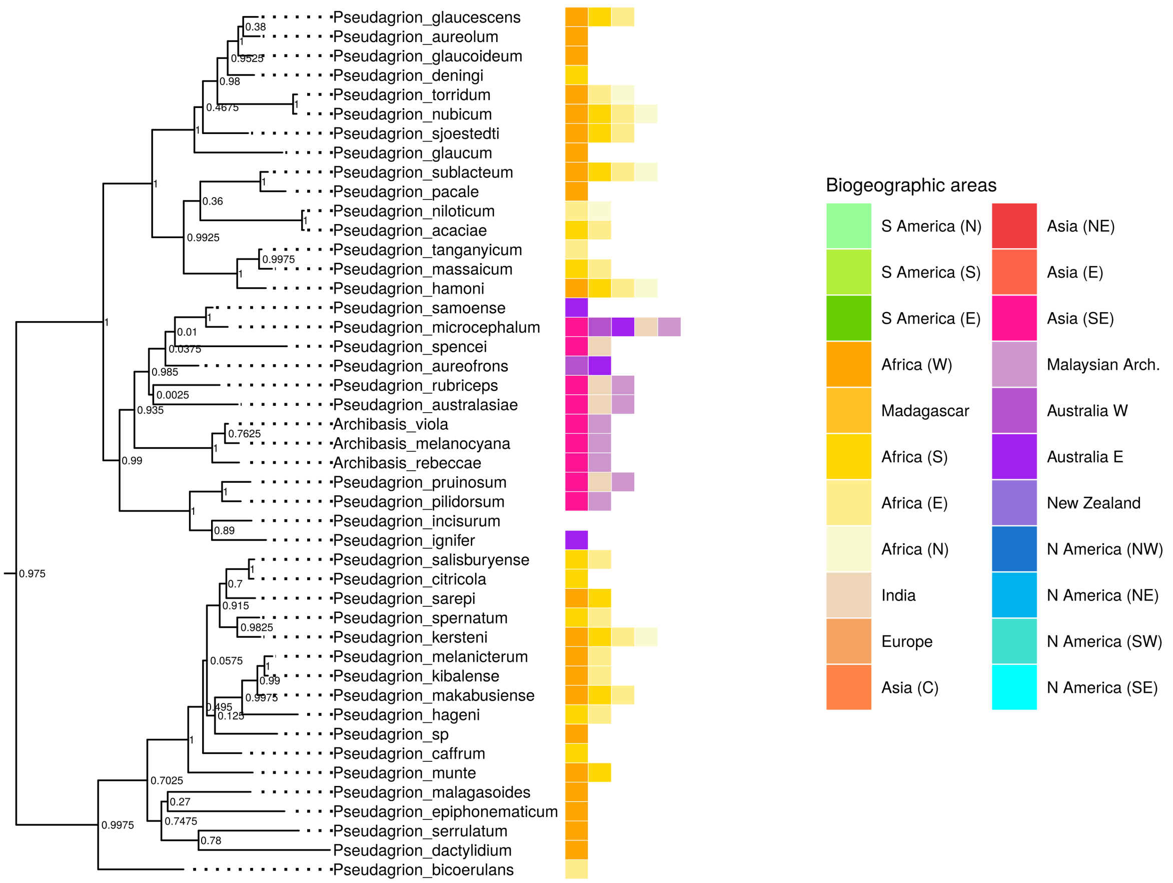

**Figure S17.** View of the clade containing the genera *Pseudagrion*, and *Archibasis* in the maximum *a posteriori* tree of Coenagrionoidea. Internal node labels represent posterior probabilities (PP).

##### ***Erythromma* and *Paracercion***

Our phylogenetic inference challenges an early neighbor-joining phylogeny that recovered both *Erythromma* and *Paracercion* as monophyletic (Weekers and Dumont 2004). Instead, we found that *Paracercion* and *Erythromma*, excluding *E. lindenii*, are monophyletic, while *E. lindenii* is a well supported outgroup to the rest of the clade (PP = 0.92, Fig. S18). Our study includes all species of *Erythromma* and eight out of the approximately 12 species of *Paracercion*. Phylogenomic studies would be beneficial to resolve the phylogenetic position of *E. lindenii*, potentially calling for a revision of the genus *Erythromma*.

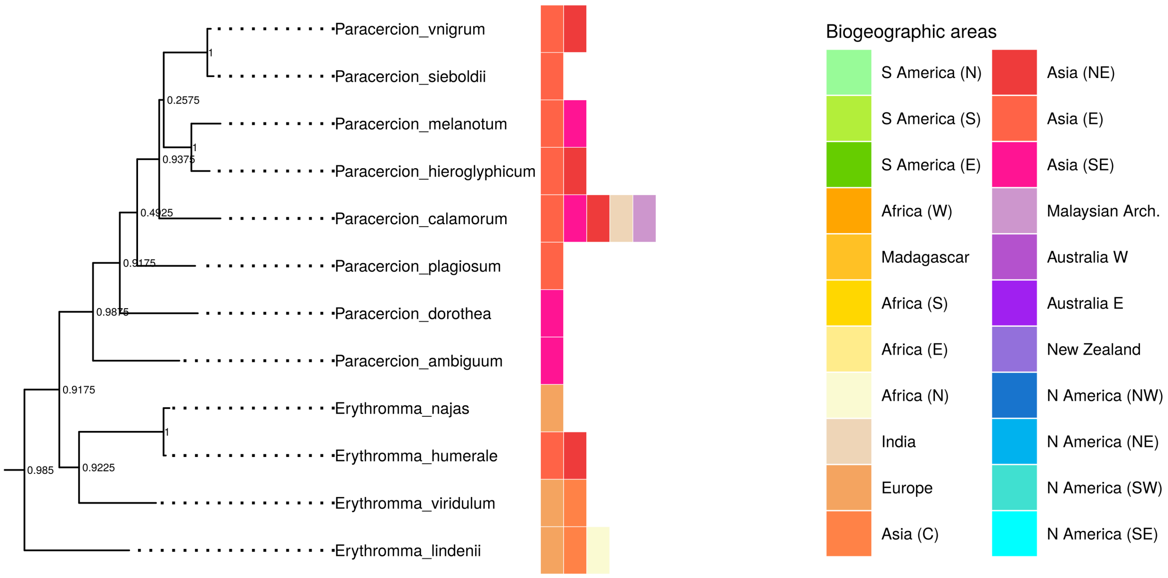

**Figure S18.** View of the clade containing the genera *Erythromma*, and *Paracercion* in the maximum *a posteriori* tree of Coenagrionoidea. Internal node labels represent posterior probabilities (PP).

##### ***Telebasis*, and *Minagrion***

*Telebasis* is one of the largest genera of Neotropical pond damselflies, with approximately 57 species. We uncovered a strongly supported clade (PP = 0.95) comprising the smaller genus *Minagrion*, with a total of 6 species, and a subset of Southamerican *Telebasis* (Fig. S19). To our knowledge, there is no other phylogenetic study of *Telebasis*, and not molecular phylogeny encompassing both *Telebasis* and *Minagrion* representatives. A morphological synopsis of *Telebasis* (2009) differentiated this genus from *Minagrion* on the basis of a tubercle on the sternum of the first abdominal segment, which is present in *Minagrion* but absent in *Telebasis*, as well as in many other pond damselfly genera. A recent morphological review of *Minagrion* did not directly compare the two genera (Vilela et al. 2020). While we tentatively conclude that *Telebasis* is likely paraphyletic, further work is therefore required to fully resolve the affinities of all taxa attributed to Minagrion.

We also noted that one species of *Telebasis* (*T. livida*) had an unexpected position in our phylogenetic inference, showing a close relationship to *Aeolagrion* and the Neotropical helicopter damselflies (subfamily Pseudostigmatinae). Our genetic data for *T. livida* comes from one of the oldest museum specimens used in this study, collected in 1999, and for which we could only amplify one mitochondrial marker. We therefore cannot rule out contamination of a sample with a very low starting concentration of DNA as causing an erroneous inference. Work with freshly collected tissues is required to clarify the affinity of *T. livida*.

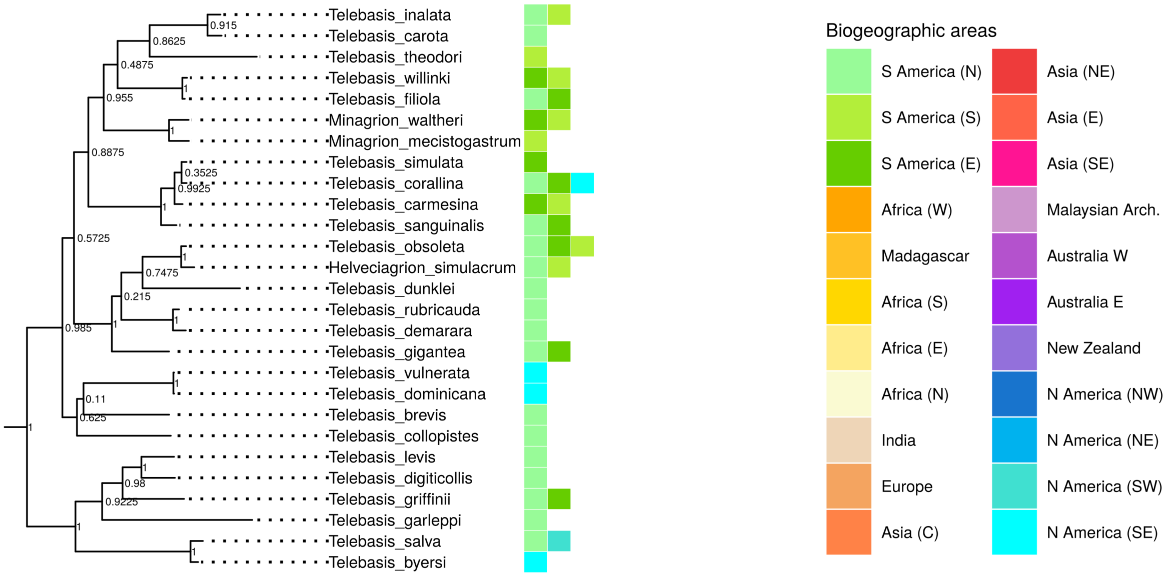

**Figure S19.** View of the clade containing the genera *Minagrion*, and *Telebasis* in the maximum *a posteriori* tree of Coenagrionoidea. Internal node labels represent posterior probabilities (PP).

##### ***Teinobasis*, *Inpabasis* and *Papuagrion***

*Teinobasis* is one of the largest genera of pond damselflies, with approximately 77 species distributed from the Western coast of the Indian Ocean to Micronesia, including Eastern Australia, Papua New Guinea and the Malaysian Archipelago. We recovered a clade with relatively high support (PP = 0.89) that includes both the genus *Papuagrion*, endemic to Papua New Guinea, and most of the Eastern *Teinobasis* species (distributed across Micronesia, Solomon Islands, Papua New Guinea and Eastern Australia) (Fig. S20). This clade is also strongly supported in Toussaint et al. (2019). Yet, our taxonomic sampling for both genera is limited (~10% of *Papuagrion* and ~20% of *Teinobasis*), and relationships within *Teinobasis* species are generally uncertain.

Unexpectedly, we also found the genus *Inpabasis* from South America, to have *Teinobasis alluaudi*, from Eastern Africa, Madagascar and the Seychelles, as its closest relative, and thus contribute to the paraphyly of *Teinobasis* (Fig. S20). However, this relationship had very weak support in our analysis (P = 0.06). Different analyses have recovered contrasting results on the phylogenetic affinities of *Inpabasis*, as a close relative of *Leptagrion* and *Bromeliagrion* (Toussaint et al. 2019), a close relative *Bromeliagrion* and Pseudostigmatinae (Dijkstra et al. 2014), or an outgroup to *Teinobasis* (Dijkstra et al. 2014). A phylogenetic study with dense taxonomic sampling and higher genetic coverage of these forest-dwelling 'ridge-face' pond damselflies (*Bromeliagrion*, *Inpabasis*, *Leptagrion*, *Papuagrion* and *Teinobasis*) is necessary to resolve their enigmatic relationships.

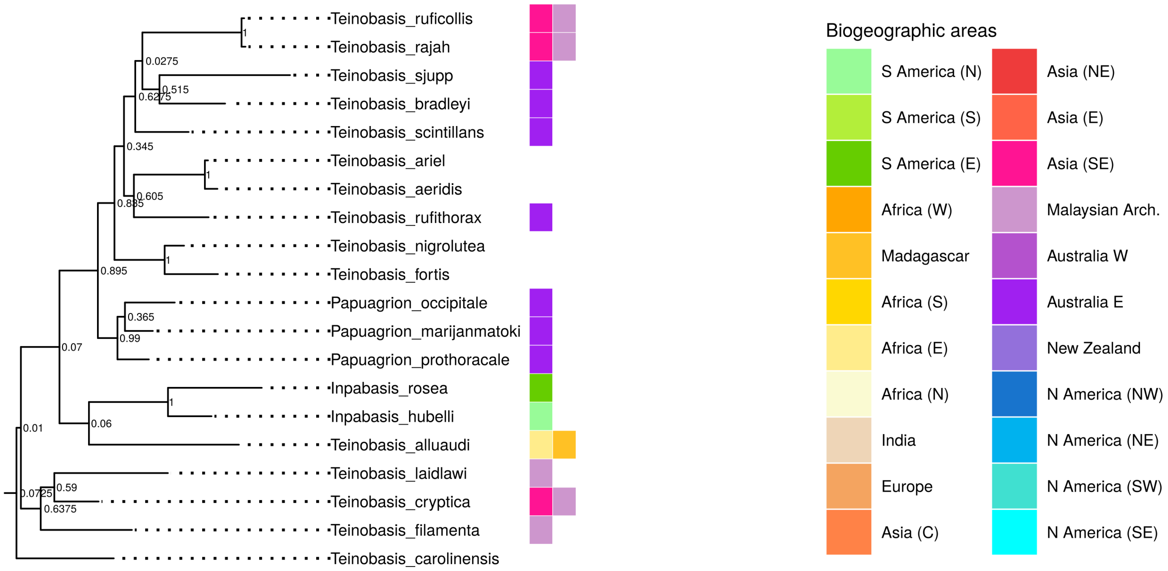

**Figure S20.** View of the clade containing the genera *Inpabasis*, *Papuagrion*, and *Teinobasis* in the maximum *a posteriori* tree of Coenagrionoidea. Internal node labels represent posterior probabilities (PP). Taxa with missing distribution data are endemic to Pacific Islands.

##### ***Mecistogaster* and *Platystigma***

The genus *Platystigma* Kennedy, 1920 was erected for a group of the Neotropical helicopter damselflies (now in the subfamily Pseudostigmatinae), based on morphological characters of their genitalia. *Platystigma* was then subsumed into *Mecistogaster* in Garrison et al. (2010) and revalidated by Machado and Lacerda (2017). Consistent with a recent molecular study focusing on this clade (Toussaint et al. 2019), we find substantial support (PP = 0.91) for the paraphyly of *Mecistogaster* if *Platystigma* is recognized (Fig. S21). Our results thus support the taxonomic designation of *Mecistogaster* for all taxa in the clade, used by Garrison et al. (2010).

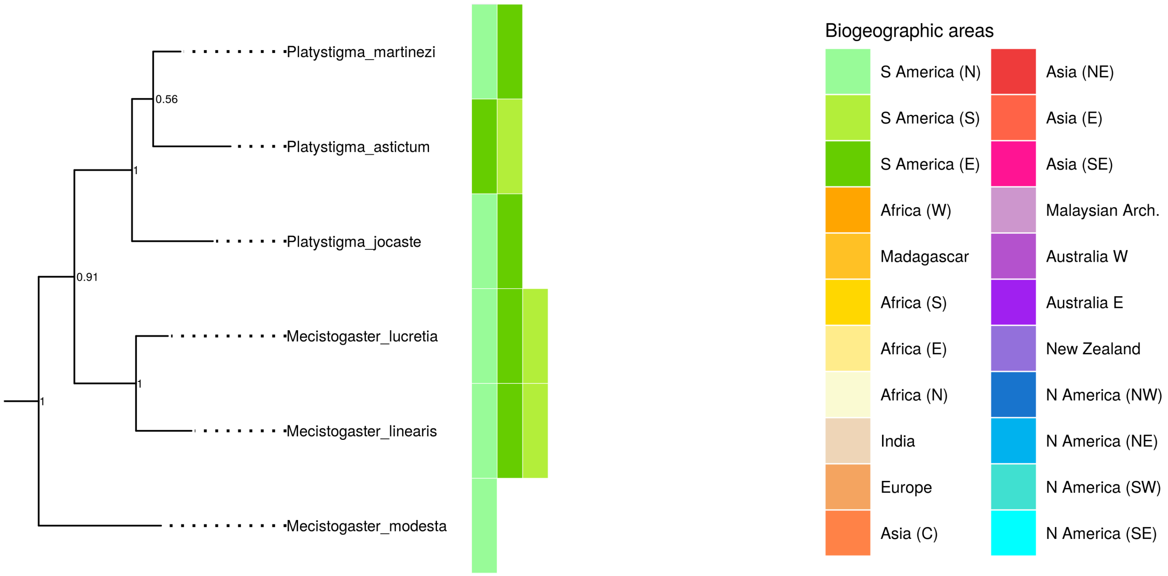

**Figure S21.** View of the clade containing the genera *Mecistogaster*, and *Platystigma* in the maximum *a posteriori* tree of Coenagrionoidea. Internal node labels represent posterior probabilities (PP).

##### ***Psaironeura*, *Amazoneura*, and *Forcepsioneura***

Our phylogentic analysis recovered a close relationship among three relatively small genera of Neotropical threadtails (subfamily Protoneurinae), *Amazoneura*, *Forcepsioneura*, and *Psaironeura* (PP = 0.99, Fig. S22). A close relationship between *Forcepsioneura*, and *Psaironeura* was also found in a previous phylgenetic study of Neotropical threadtails based on morphology (Pessacq 2008). Here we found that the South American species of *Psaironeura* form a clade with the South American genera *Amazoneura* and *Forcepsioneura*, which is in turn sister of the Central American *Psaironeura remissa* (Fig. S22). However, the paraphyly of *Psaironeura* hinges on a node with modest support (PP = 0.40). We thus conclude that further studies of the clade are required to test the monophyly of *Psaironeura*.

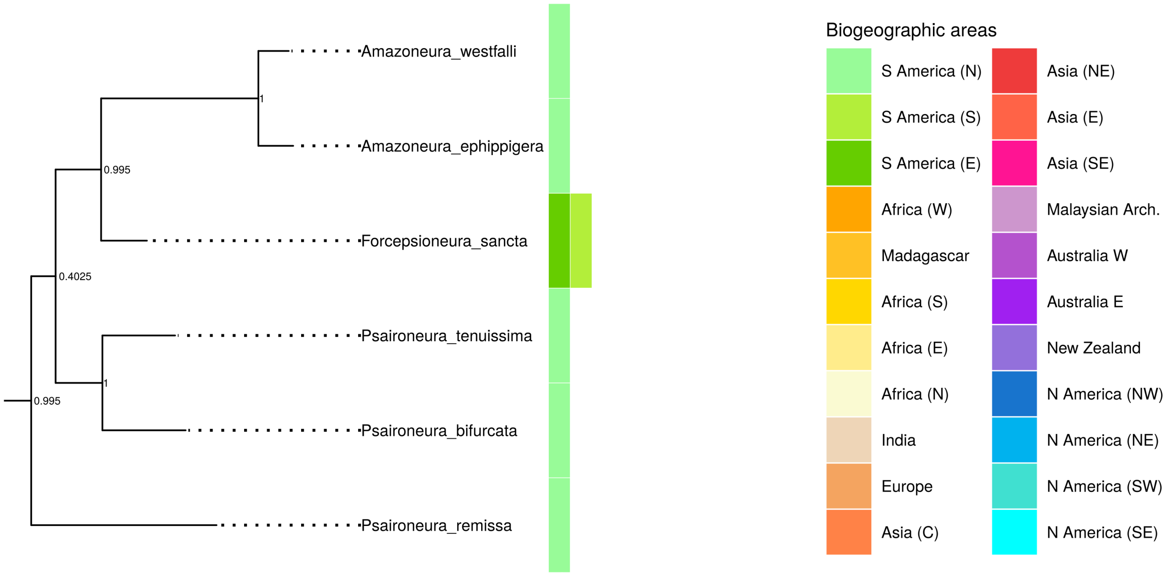

**Figure S22.** View of the clade containing the genera *Amazoneura*, *Forcepsioneura*, and *Psaironeura* in the maximum *a posteriori* tree of Coenagrionoidea. Internal node labels represent posterior probabilities (PP).

##### ***Elattoneura*, *Nososticata* and *Prodasineura***

*Elattoneura*, *Nososticata* and *Prodasineura* are three speciose genera of featherlegs (family Platycnemididae), which together account for most of the subfamily Disparoneurinae. Our phylogenetic inference shows strong support (PP = 0.95) for a clade composed exclusively of the African representatives of *Elattoneura*, and strong support (PP =1.00) for a clade including both *Elattoneura* and *Prodasineura* species from India, Southeast Asia, and the Malaysian Archipelago (Fig S23), thus rendering *Elattoneura* paraphyletic. This result was foreshadowed in Dijkstra et al. (2014) although with a more limited taxonomic sampling.

We also recovered the African *Elattoneura* as a sister clade to *Nososticta*, distributed mainly in Australia and Papua New Guinea (Fig S23). This relationship was also found by Dijkstra et al. (2014), but we caution that in the present analysis it was weakly supported (PP = 0.12). To our knowledge, there are no phylogenetic studies focused on Disparoneurinae. Such studies would benefit from a more extensive sampling of *Nososticta*, which has underrepresented (~10% of species sampled) in the present study.

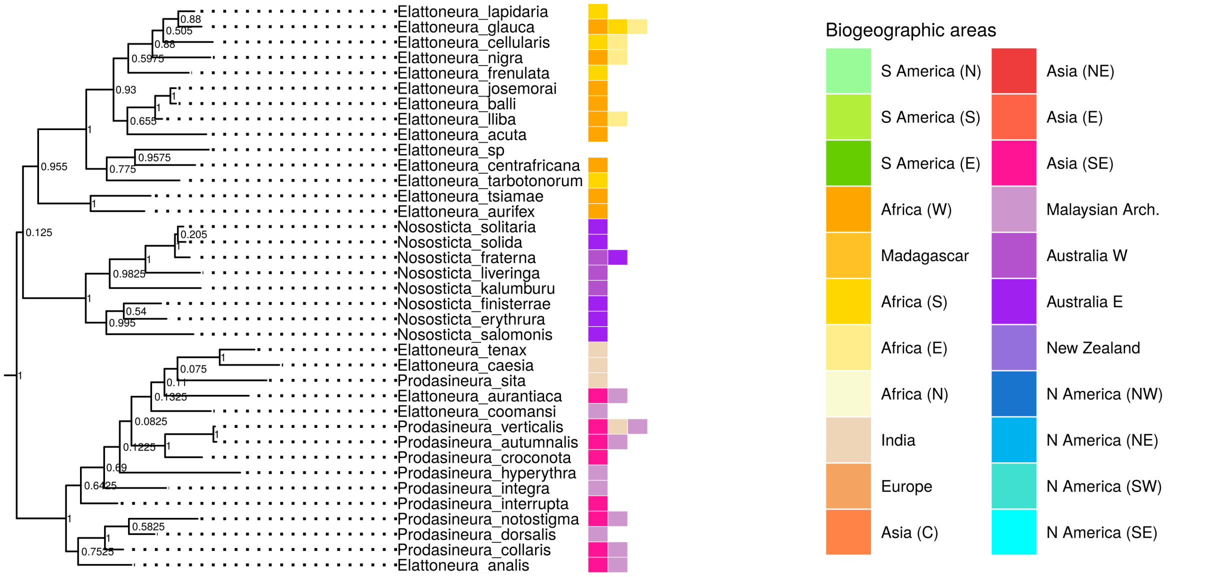

**Figure S23.** View of the clade containing the genera *Elattoneura*, *Nososticta*, and *Prodasineura* in the maximum *a posteriori* tree of Coenagrionoidea. Internal node labels represent posterior probabilities (PP).

##### ***Platycnemis*, *Pseudocopera* and *Matticnemis***

We obtained strong support (PP = 1.00) for a clade consisting of the East Asian *Platycnemis phyllopoda* and the mostly East Asian genus *Pseudocopera*, thus pointing to paraphyly in *Platycnemis* (Fig. S24). The remaining species of *Platycnemis*, distributed across the Palearctic, and the Southeast Asian monotypic genus *Matticnemis* formed a sister clade to the one described above, with moderate support (PP = 0.63, Fig. S24). Evidence of paraphyly in *Platycnemis* due to the phylogenetic placement of *Pseudocopera* was already suspected by Dijkstra et al. (2014). In contrast, Dijkstra et al. (2014) recovered *Matticnemis* as a sister clade to *Platycnemis* and *Pseudocopera*. In the present analysis, the affinity of *Matticnemis* within *Platycnemis* is weakly supported (PP = 0.31). Further phylogenetic studies are thus required to resolve the relationships among these three genera.

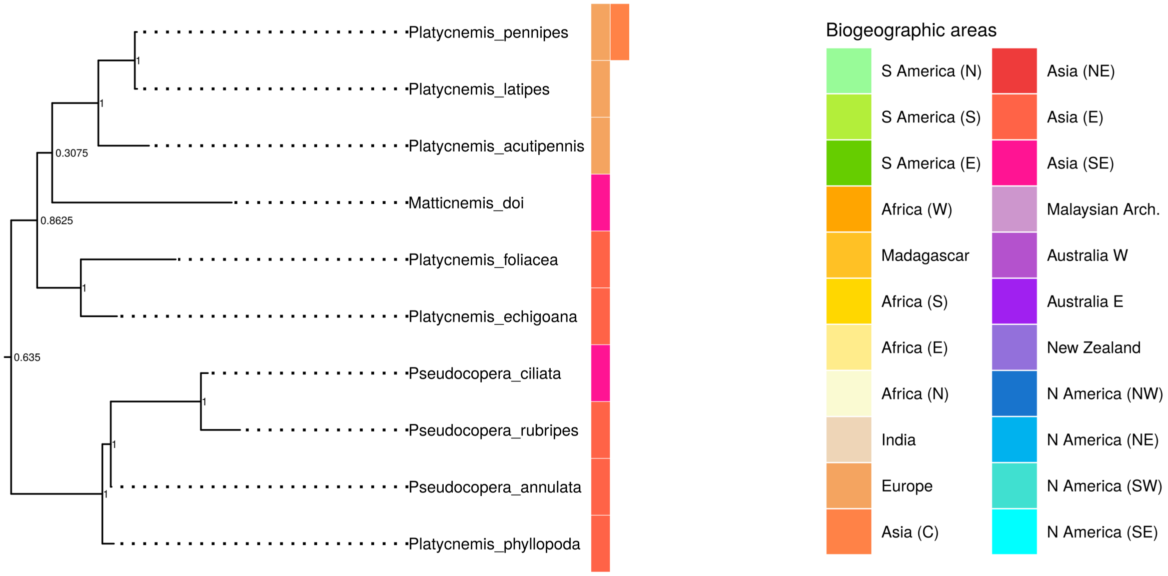

**Figure S24.** View of the clade containing the genera *Matticnemis*, *Platycnemis*, and *Pseudocopera* in the maximum *a posteriori* tree of Coenagrionoidea. Internal node labels represent posterior probabilities (PP).

##### ***Coeliccia*, *Calicnemia* and *Indocnemis***

*Coeliccia* and *Calicnemia*, with 66 and 24 species respectively, are two relatively large genera comprising most of the Calicnemiinae subfamily of featherlegs. *Indocnemis* is a small genus with only three species described. *Coeliccia* is distributed across East Asia, Southeast Asia and the Malaysian Archipelago, with at least one species reaching into India. *Calicnemia* and *Indocnemis* are instead primarily distributed in Southeast Asia. The monophyly of *Calicnemia* was strongly supported (PP = 1.00), but the genus was nested within a East to Southeast Asian clade of *Coeliccia* (Fig. S25), albeit with low support (PP = 0.43). The two species of *Indocnemis* sampled in this study also contributed to the paraphyly of *Coeliccia* and were not recovered as sister taxa (Fig. S25). Our results thus suggest that a revision of *Coeliccia*, *Indocnemis* and potentially *Calicnemia*, based on further phylegenetic research is warranted. We are unaware of previous studies focusing on these genera, but Dijkstra et al. (2014) also recovered both *Coeliccia* and *Indocnemis* as paraphyletic, while their analysis returned *Calicnemia* as diverging from the common ancestor of the other two genera.

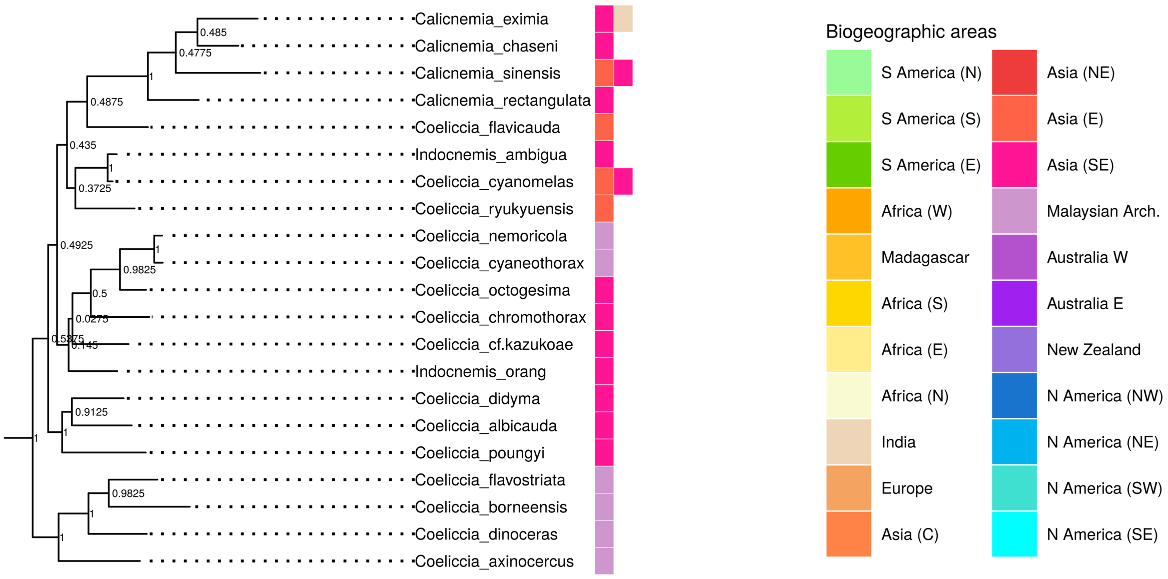

**Figure S25.** View of the clade containing the genera *Calicnemia*, *Coeliccia*, and *Indocnemis* in the maximum *a posteriori* tree of Coenagrionoidea. Internal node labels represent posterior probabilities (PP).

#### Comparison of node age estimates

We inferred ancestral biogeographic areas and divergence times for Coenagrionoidea under two biogeographic models: 1) a model with a strongly informed root age prior, based on two phylogenomic studies (Suvorov et al. 2021; Kohli et al. 2021) and 2) a model with a weakly-informed broadly-uniform prior (see Extended Methods). Here, we compare node age estimates in these two models against each other and to a pure-birth model using observed fossils as age constraints on internal nodes (Table S10). We also report age estimates of comparable clades in previous studies and, whenever available, dated fossils in the Paleobiology Database https://paleobiodb.org/ (Table S10).

**Table S10.** Divergence time estimates under two biogeographic dating models with different root age priors in comparison to models with traditional fossil calibrations. Divergence time estimates are shown for comparable clades across studies and models. Suvorov et al. (2021) inferred a backbone phylogeny for Odonata using a supermatrix of 1603 gene orthologs and fossil calibrations with 20 crown fossil constraints. We report the mean of 5 independent runs in Suvorov et al. (2021). Toussaint et al. (2019) sampled nine gene fragments from 33 Coenagrionoidea genera and 85 species, focusing on Pseudostigmatinae (helicopter damselflies) and relied on three fossil calibrations to constrain internal nodes. Waller and Svensson (2017) produced a species-level phylogeny for all Odonata taxa with openly available genetic data for at least one of 14 gene fragments. The phylogeny in Waller and Svensson (2017) used multiple fossil calibrations and its topology was constrained by taxonomy, including phylogenetically unsupported family classifications within Coenagrionoidea. A few other studies have dated phylogenies for specific coenagrionid genera: (1) Beatty et al. (2017) focused on the genera *Nesobasis* and *Melanesobasis* from Fiji, and used a combination of fossil data and island emergence times for calibration, (2) Swaegers et al. (2014) dated the origin of *Coenagrion* using prior information on evolutionary rates in three loci, (3) Callahan and McPeek (2016) and (4) Blow et al. (2021) used multi-species coalescent models and similar fossil calibrations to date the origin of *Enallagma* and *Ischnura*, respectively.

| Clade | Strongly.informed | Weakly.informed | Fossil.constraints | Suvorov.et.al.2021 | Toussaint.et.al.2019 | Waller.Svensson.2017 | Genus.level.studies | Earliest.fossil |
|---|---|---|---|---|---|---|---|---|
| Coenagrionoidea | 105.7 | 66.9 | 102.1 | 115.8 | 131.6 | 72.7 | | |
| Platycnemididae | 99.0 | 62.4 | 94.5 | 98.3 | 108.9 | | | 99.6-93.5 |
| *Platycnemis* | 62.4 | 40.4 | 24.1 | | 33.2 | 50.0 | | 38-33.9 |
| Coenagrionidae | 105.5 | 66.7 | 98.0 | 89.9 | 118.0 | | | |
| Ridge-face | 101.6 | 64.7 | 62.2 | 64.5 | 110.8 | | | |
| *Mecistogaster + Megaloprepus* | 64.7 | 41.1 | 26.8 | 23.2 | 65.4 | 41.9 | | |
| *Nehalennia* | 42.6 | 28.1 | 17.0 | | 23.8 | 14.8 | | 23.0-16.0 |
| *Melanesobasis* | 29.6 | 21.3 | 13.9 | | | 22.4 | 8.5 (1) | |
| *Argia* | 73.1 | 47.5 | 27.5 | | 34.6 | 47.0 | | 23.0-16.0 |
| Core | 97.1 | 62.2 | 90.4 | 62.9 | 100.6 | 58.0 | | |
| *Coenagrion* | 45.4 | 28.4 | 23.6 | | | 43.2 | ~ 15 (2) | |
| *Nesobasis* | 46.1 | 31.5 | 22.9 | | | 33.1 | 11.8 (1) | |
| *Enallagma* | 53.2 | 34.8 | 23.1 | | 9.0 | 34.0 | 9.0 (3) | |
| *Ischnura* | 56.0 | 36.3 | 20.8 | 26.6 | 29.7 | 34.7 | 16.2 (4) | 20.4-13.6 |

*Note:* In contrast to Dijkstra et al. (2014), the 'ridge-face' clade here includes the genus *Argia*.

**Root age estimates under strongly informed and weakly informed priors**

Root age estimates were highly sensitive to prior information. The biogeographic dating analysis using a strongly informed root age prior based on recent phylogenomic studies (Suvorov et al. 2021; Kohli et al. 2021) returned a mean age estimate for the MRCA of Coenagrionidae and Platycnemididae of 105 Ma (95% HPD interval = 64 − 145 Ma; Fig. S26). In contrast, applying a broad uniform prior on the root age resulted in a much younger estimate of 67 Ma (95% HPD interval = 40 − 118 Ma; Fig. S26). An analysis excluding empirical paleogeographic data produced, as expected, a flat posterior root age distribution between 240 and 0 Ma (Fig. S26).

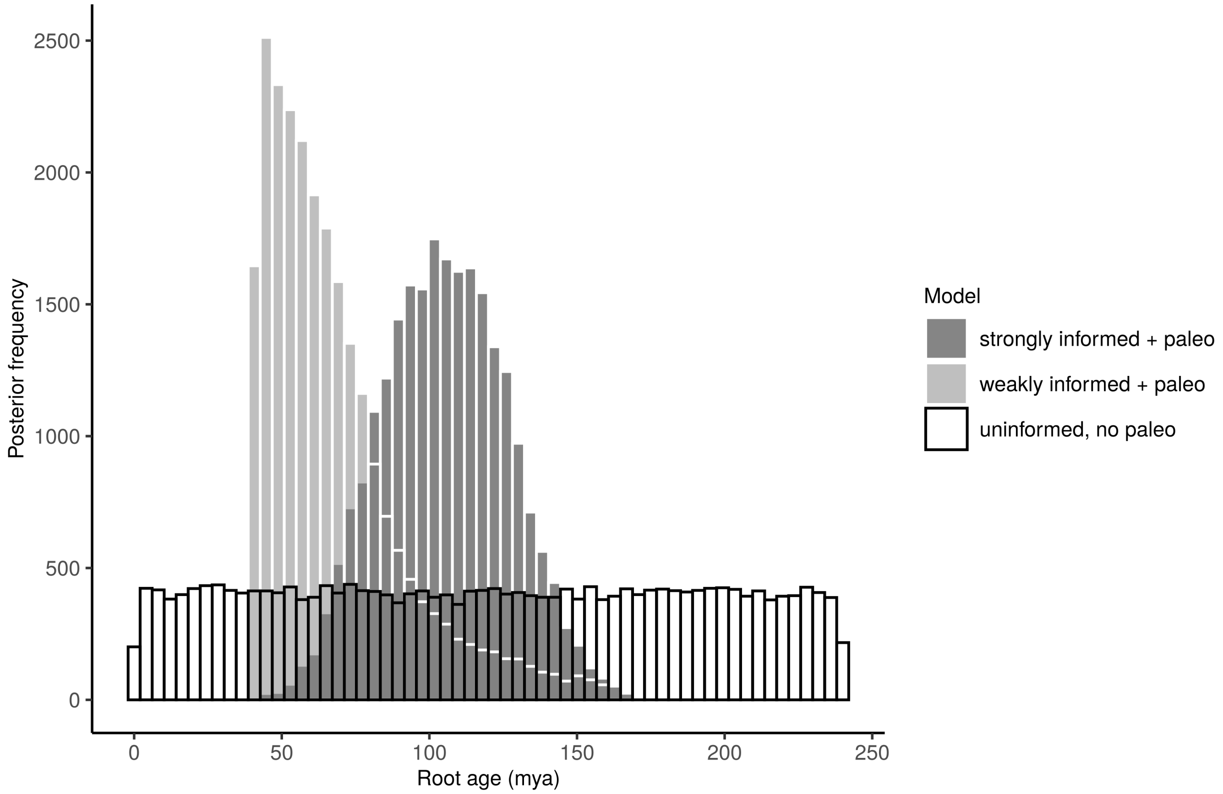

**Figure S26.** Posterior distribution of age estimates for the MRCA of pond damselflies and fetherlegs, under three alternative biogeogrphic dating models. The "strongly informed + paleo" model uses the empirical paleographic model in Landis (2017) to inform divergence times throughout the tree and a strong, normally distributed prior on the root age, with mean = 120 and sd = 20 Ma, based on two recent phylogenomic studies (Kohli et al. 2021; Suvorov et al. 2021). The "weakly informed + paleo" model uses the same empirical paleographic model, but assumes *a priori* that Coenagrionoidea could have originated any time, with equal probability, between 240-40 Ma. Finally, the "weakly informed, no paleo" model was used to confirm that biogeographic dating does not introduce unexpected biases in node ages. It does not incorporate any paleogeographic data and assumes that Coenagrionoidea could have originated any time, with equal probability, between 240-0 Ma.

#### Ancestral distribution of pond damselflies and featherlegs

The general features of ancestral biogeography were largely shared by the two models (Fig. 1; S27-S33). We first plot a cladogram of the most ancestral nodes of Coenagrionoidea under the weakly informed root age prior (Fig. S27), similar to the one in the bottom right corner of Fig. 1 for the analysis with a strongly informed root age prior.

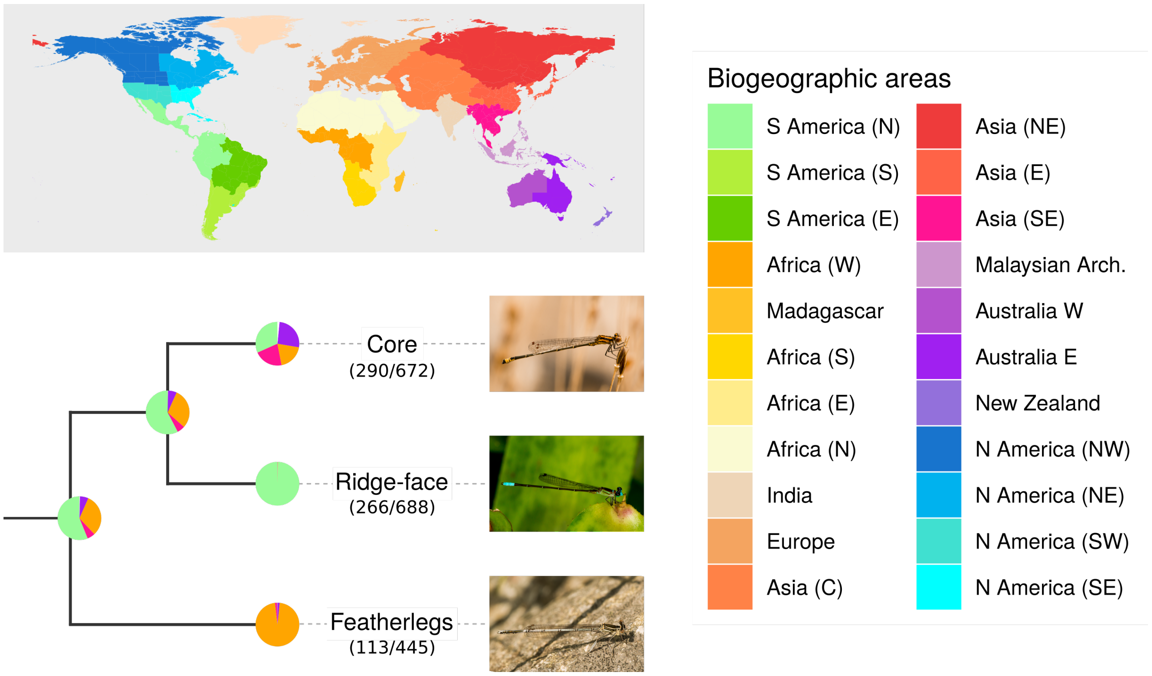

**Figure S27.** Inferred ancestral ranges for the most recent common ancestor of Coenagrionoidea, and for each of the three major clades in the superfamily. Pies on the cladogram show the proportion of each ancestral state in a random sample of 1000 posterior trees. The empirical paleogeographic model in Landis (2017) was used to determine dispersal edges between 25 geographic areas across the globe (Greenland, Antarctica (E) and Antarctica (W) not shown in the colour legend) and according to 3 different dispersal modes (see Methods and Extended Methods). The fraction of taxa sampled is shown in parenthesis. One representative species is illustrated for each clade: *Acanthagrion adustum* ('core'), *Leptagrion elongatum* ('ridge-face'), *Platycnemis pennipes* (featherlegs). Photos: EIS.

To make taxon names readable, results from both models (strongly informed and weakly informed) are presented separately for each of the three main clades discussed in this paper: the featherlegs (family Platycnemididae; Fig. S28-S29), the 'ridge-face' clade of Coeangrionidae (Fig. S30-31) and the 'core' clade of Coenagrionidae (Fig. 32-S33).

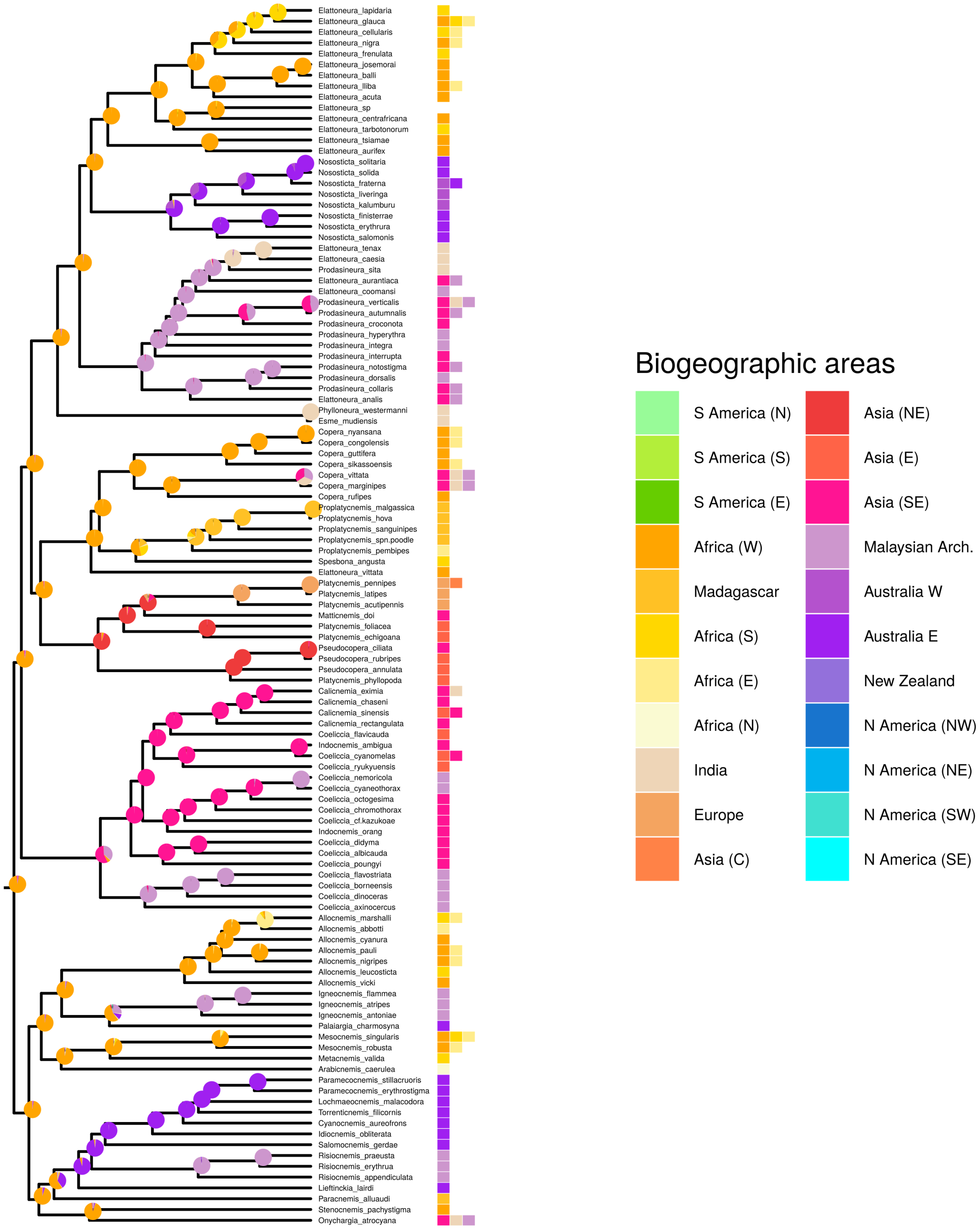

**Figure S28.** Ancestral range and time tree estimation in the damselfly superfamily Coenagrionoidea, using a data-dependent biogeographic model based on Landis (2017). Speciation times and ancestral ranges were jointly estimated using empirical paleogeography, molecular sequence data, **an informed root age prior (dnNormal(120, 20))** based on previous studies (Suvorov et al. 2021; Kohli et al. 2021), and distribution data for extant taxa. Species ranges were classified to one or more of 25 geographic areas used in the paleogeographic model. Phylogenetic inference was summarised using the maximum *a posteriori* tree. For clarity, the figure shows only the **featherleg clade** (family Platycnemididae).

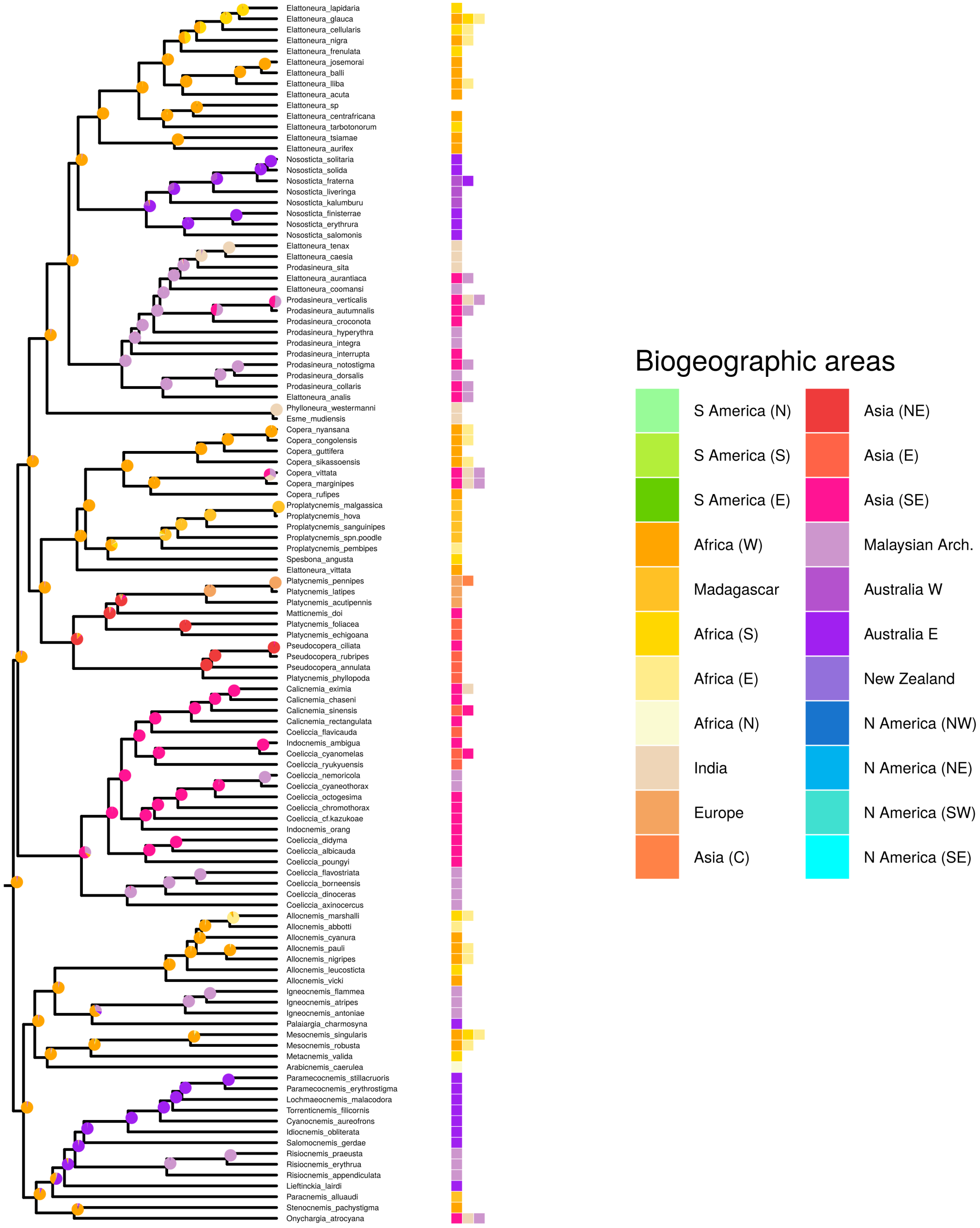

**Figure S29.** Ancestral range and time tree estimation in the damselfly superfamily Coenagrionoidea, using a data-dependent biogeographic model based on Landis (2017). Speciation times and ancestral ranges were jointly estimated using empirical paleogeography, molecular sequence data, **a broad uniform root age prior (dnUniform(40,240))**, and distribution data for extant taxa. Species ranges were classified to one or more of 25 geographic areas used in the paleogeographic model. Phylogenetic inference was summarised using the maximum *a posteriori* tree. For clarity, the figure shows only the **featherleg clade** (family Platycnemididae).

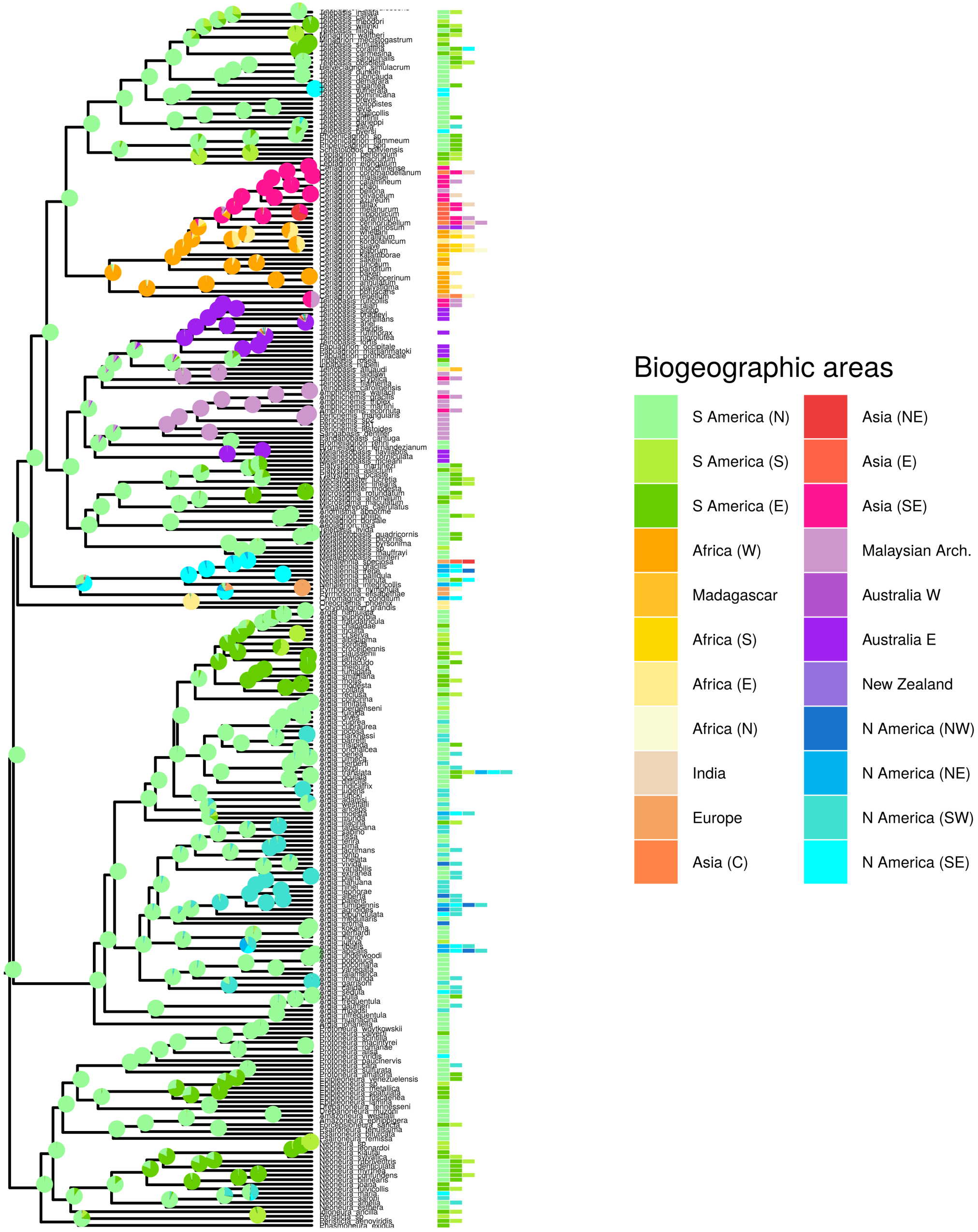

**Figure S30.** Ancestral range and time tree estimation in the damselfly superfamily Coenagrionoidea, using a data-dependent biogeographic model based on Landis (2017). Speciation times and ancestral ranges were jointly estimated using empirical paleogeography, molecular sequence data, **an informed root age prior (dnNormal(120, 20))** based on previous studies (Suvorov et al. 2021; Kohli et al. 2021), and distribution data for extant taxa. Species ranges were classified to one or more of 25 geographic areas used in the paleogeographic model. Phylogenetic inference was summarised using the maximum *a posteriori* tree. For clarity, the figure shows only the **'ridge-face' clade** (family Coenagrionidae).

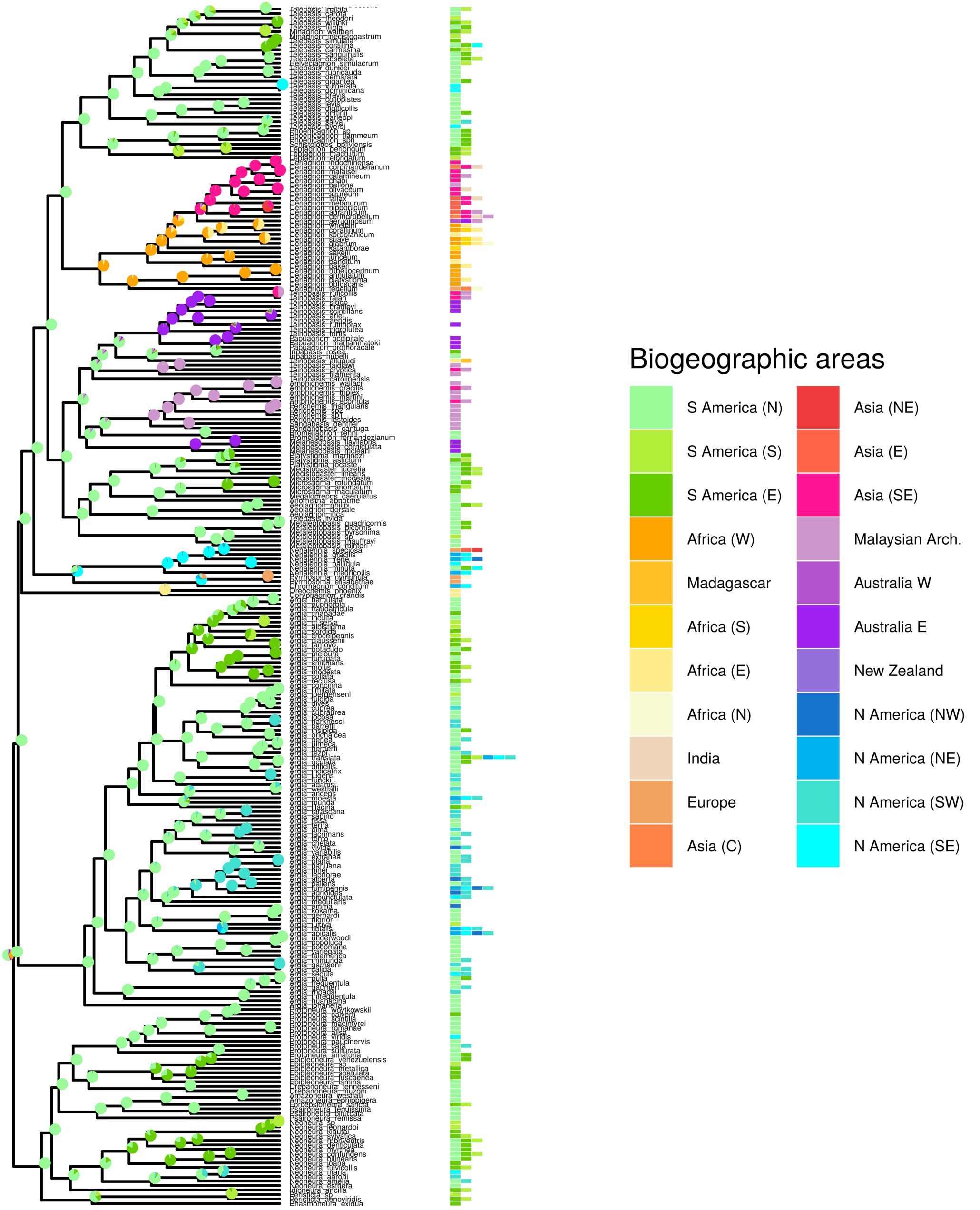

**Figure S31.** Ancestral range and time tree estimation in the damselfly superfamily Coenagrionoidea, using a data-dependent biogeographic model based on Landis (2017). Speciation times and ancestral ranges were jointly estimated using empirical paleogeography, molecular sequence data, **a broad uniform root age prior (dnUniform(40,240))**, and distribution data for extant taxa. Species ranges were classified to one or more of 25 geographic areas used in the paleogeographic model. Phylogenetic inference was summarised using the maximum *a posteriori* tree. For clarity, the figure shows only the **'ridge-face' clade**(family Coenagrionidae).

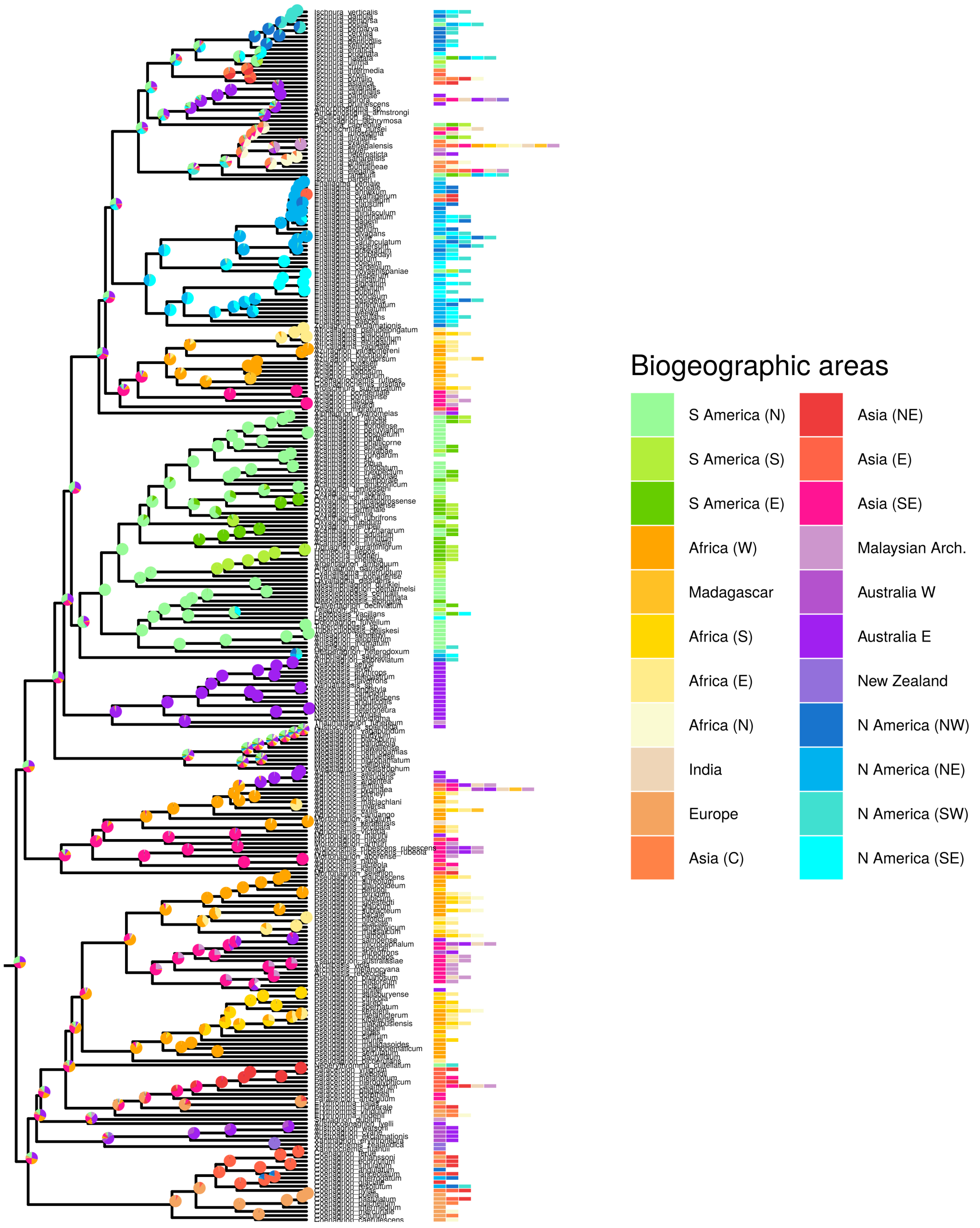

**Figure S32.** Ancestral range and time tree estimation in the damselfly superfamily Coenagrionoidea, using a data-dependent biogeographic model based on Landis (2017). Speciation times and ancestral ranges were jointly estimated using empirical paleogeography, molecular sequence data, **an informed root age prior (dnNormal(120, 20))** based on previous studies (Suvorov et al. 2021; Kohli et al. 2021), and distribution data for extant taxa. Species ranges were classified to one or more of 25 geographic areas used in the paleogeographic model. Phylogenetic inference was summarised using the maximum *a posteriori* tree. For clarity, the figure shows only the **'core' clade** (family Coenagrionidae).

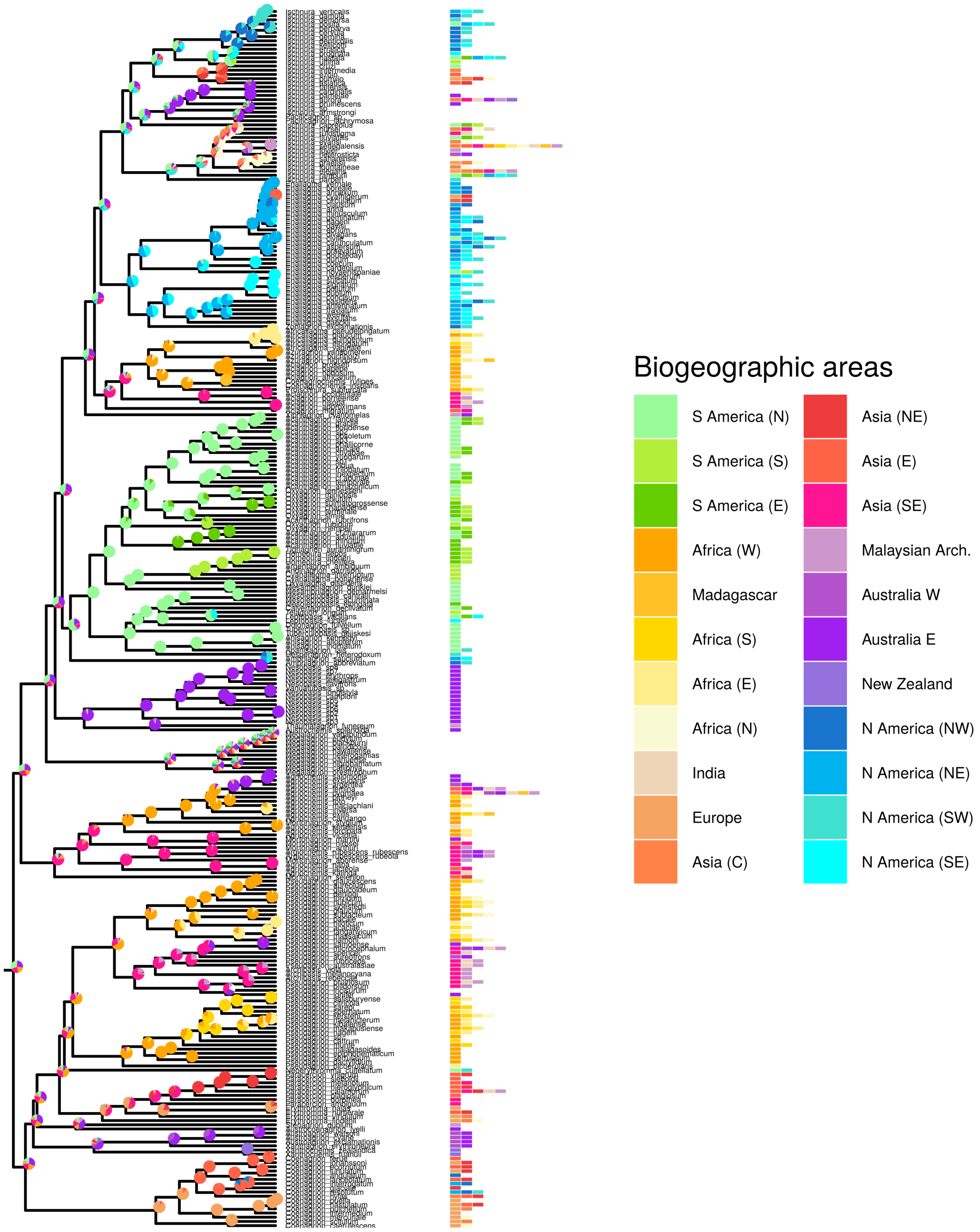

**Figure S33.** Ancestral range and time tree estimation in the damselfly superfamily Coenagrionoidea, using a data-dependent biogeographic model based on Landis (2017). Speciation times and ancestral ranges were jointly estimated using empirical paleogeography, molecular sequence data, **a broad uniform root age prior (dnUniform(40,240))**, and distribution data for extant taxa. Species ranges were classified to one or more of 25 geographic areas used in the paleogeographic model. Phylogenetic inference was summarised using the maximum *a posteriori* tree. For clarity, the figure shows only the **'core' clade** (family Coenagrionidae).

#### Diversification dynamics

##### Diversification through time

Speciation and thereby diversification have slowed down since the origin of Coenagrionoidea (Fig. 2; S34). The main text shows the results of the EBD analysis using a MAP tree informed by previous estimates for the root age of Coenagrionoidea (Suvorov et al. 2021; Kohli et al. 2021). Here, we show the results of the same analysis using the MAP tree from the biogeographic dating analysis without this informed root age prior (Fig. S34). These results contrast our main analysis in two ways. First, there is a sharper decline in the rate of speciation after the early period of constant birth (Fig. 2a, vs. S34a). Second, the increase in extinction over the last 5 My is negligible and therefore, the lower bound of 95% HPD interval for the rate of net diversification near the present remained positive.

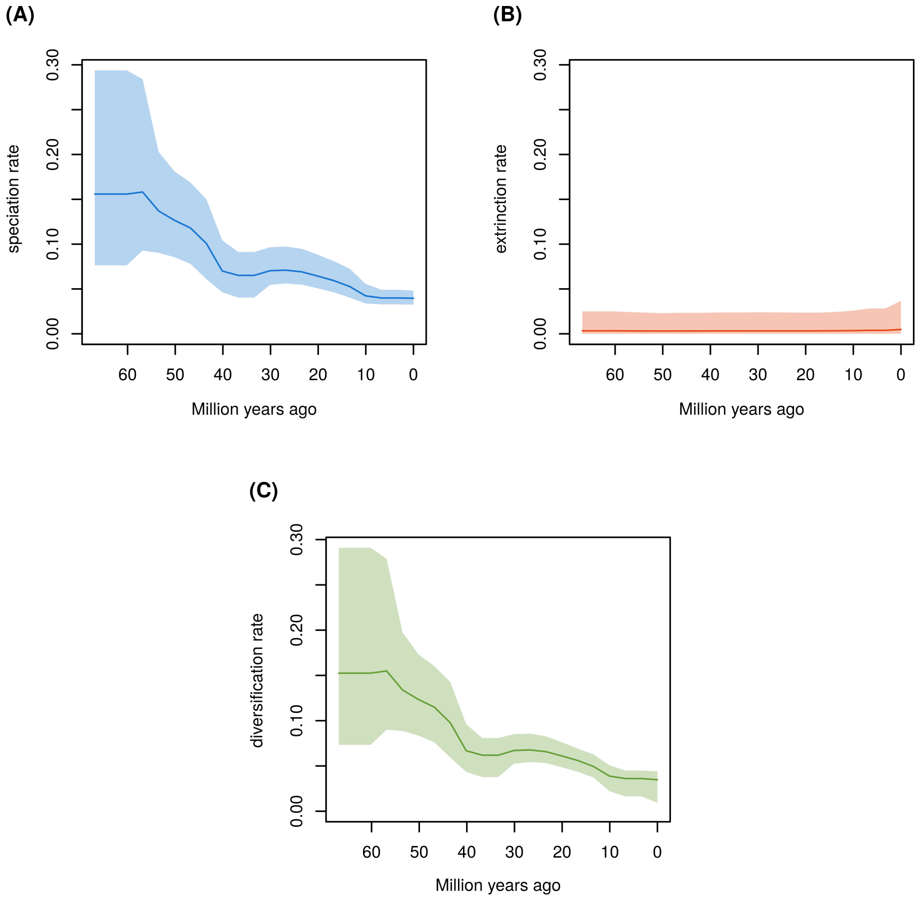

**Figure S34.** Changes in diversification rates since the MRCA of Coenagrionoidea: a) speciation, b) extinction and d) net diversification. Diversification rates were estimated on the maximum *a posteriori* (MAP) tree from the biogeographic dating analysis using a weakly informed root age prior, placing equal probability on origin times between 240 - 40 Ma. Changes in diversification rates were modelled under an episodic birth-death (EBD) process, over 20 equal-length time intervals and assuming autocorrelation among consecutive time intervals (see Extended Methods).

**Biome-dependent diversification**

After obtaining strong support for biome-dependent diversification (Table S11), we ran three HiSEE models to quantify diversification rates in tropical, warm-temperate and cold-temperate biomes, under different sets of assumptions. First, we used the MAP tree from our strongly-informed dating analysis, and character state data in which wide-ranging species were assigned to the coldest biome they inhabit. We found relatively high speciation rates in cold-temperate biomes compared to tropical and warm-temperate habitats, and relatively high extinction rates in both temperate habitats compared to the tropics (Fig. 3a-b, Table S12). Consequently, net diversification rates were similar across biomes, although with in weak slowing trend in warm-temperate habitats, and turnover rates were relatively high in both temperate biomes (Fig. 3c-d, Table S12). Ancestral state reconstructions from this modelled strongly supported a tropical ancestor of Coenagionoidea, with a relatively low background diversification rate (Fig. S35).

**Table S11.** Alternative diversification models compared using marginal likelihood (ML) approximations in RevBayes (Höhna et al. 2016). Marginal likelihoods were approximated twice using the stepping stone algorithm (Xie et al. 2010) for a hidden-state biome-dependent diversification model and a null model with state-independent diversification but background heterogeneity in diversification rates. The model with highest marginal likelihood is highlighted in **bold**. A difference in marginal likelihood $> 2$ is considered as substantial support for the biome dependent model.

| Model | ML run 1 |
|---|---|
| **biome-dependent + background** | 123.0637 |
| background only | -6121.9260 |

We then ran the HiSSE model with the same input tree but using a character matrix in which wide-ranging species were coded as ambiguous between their current states, and obtained qualitatively similar results (Table S13, Fig. S36-S37). Finally, we modelled biome-dependent diversification on the MAP tree of our weakly informed diversification analysis and again obtained qualitatively similar results (Table S14, Fig. S38-S39). In addition to testing the robustness of our results to different assumptions in the data, we asked if the model parametrisation could have been responsible for our results, by running a Markov chain of the model without data. Here we obtained, as expected, posteriors that match the prior distributions (Fig. S40-S41).

**Table S12.** Comparison of biome-dependent diversification rates in Coenagrionoidea. We report mean differences and 95% highest posterior density (HPD) intervals. PMCMC values show the fraction of posterior samples in which the difference between rates and the posterior mean difference have the opposite signs. PMCMC values $< 0.05$ are considered evidence for biome-dependent diversification. Rates were estimated under a Hidden-State Dependent Speciation and Extinction (HiSSE) model, which accommodates background heterogeneity in diversification by modelling the diversification effects of a hidden trait with two states (see Extended Methods). The HiSSE analysis was conducted on the maximum *a posteriori* (MAP) tree from a biogeographic dating analyses informed by previous estimates for the root age of Coenagrionoidea (Suvorov et al. 2021; Kohli et al. 2021). Species with distribution ranges spanning across two or three biomes were assigned to the coldest biome they inhabit.

| Rate | Biomes | Hidden state | Mean difference | Lower bound | Upper bound | PMCMC |
|---|---|---|---|---|---|---|
| Speciation | Cold-Temperate vs Warm-Temperate | 0 | 0.03 | 0.00 | 0.07 | 0.02 |
| Speciation | Cold-Temperate vs Warm-Temperate | 1 | 0.05 | 0.00 | 0.12 | 0.02 |
| Speciation | Tropical vs Cold-Temperate | 0 | -0.04 | -0.08 | -0.01 | 0.00 |
| Speciation | Tropical vs Cold-Temperate | 1 | -0.06 | -0.13 | -0.01 | 0.00 |

*(continued)*

| Rate | Biomes | Hidden state | Mean difference | Lower bound | Upper bound | PMCMC |
|---|---|---|---|---|---|---|
| Speciation | Tropical vs Warm-Temperate | 0 | -0.01 | -0.03 | 0.01 | 0.30 |
| Speciation | Tropical vs Warm-Temperate | 1 | -0.01 | -0.05 | 0.02 | 0.30 |
| Extinction | Cold-Temperate vs Warm-Temperate | 0 | 0.02 | -0.05 | 0.09 | 0.26 |
| Extinction | Cold-Temperate vs Warm-Temperate | 1 | 0.02 | -0.03 | 0.09 | 0.26 |
| Extinction | Tropical vs Cold-Temperate | 0 | -0.05 | -0.11 | 0.01 | 0.03 |
| Extinction | Tropical vs Cold-Temperate | 1 | -0.04 | -0.11 | 0.00 | 0.03 |
| Extinction | Tropical vs Warm-Temperate | 0 | -0.02 | -0.07 | 0.01 | 0.06 |
| Extinction | Tropical vs Warm-Temperate | 1 | -0.02 | -0.05 | 0.01 | 0.06 |
| Diversification | Cold-Temperate vs Warm-Temperate | 0 | 0.01 | -0.04 | 0.06 | 0.25 |
| Diversification | Cold-Temperate vs Warm-Temperate | 1 | 0.03 | -0.02 | 0.08 | 0.07 |
| Diversification | Tropical vs Cold-Temperate | 0 | 0.01 | -0.04 | 0.05 | 0.44 |
| Diversification | Tropical vs Cold-Temperate | 1 | -0.03 | -0.08 | 0.03 | 0.14 |
| Diversification | Tropical vs Warm-Temperate | 0 | 0.02 | -0.01 | 0.05 | 0.07 |
| Diversification | Tropical vs Warm-Temperate | 1 | 0.01 | -0.02 | 0.03 | 0.20 |
| Turnover | Cold-Temperate vs Warm-Temperate | 0 | 0.05 | -0.86 | 0.76 | 0.42 |
| Turnover | Cold-Temperate vs Warm-Temperate | 1 | 0.05 | -0.31 | 0.45 | 0.42 |
| Turnover | Tropical vs Cold-Temperate | 0 | -0.50 | -1.01 | 0.12 | 0.05 |
| Turnover | Tropical vs Cold-Temperate | 1 | -0.24 | -0.67 | 0.03 | 0.05 |
| Turnover | Tropical vs Warm-Temperate | 0 | -0.46 | -1.24 | 0.12 | 0.06 |
| Turnover | Tropical vs Warm-Temperate | 1 | -0.19 | -0.55 | 0.03 | 0.06 |

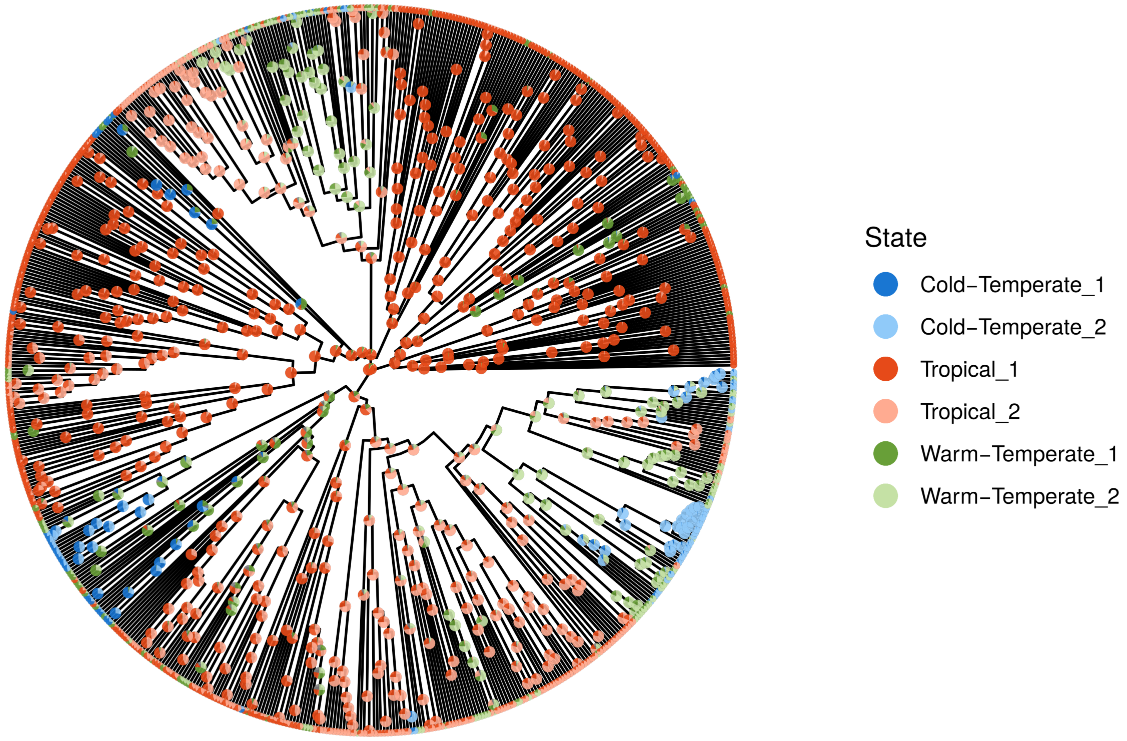

**Figure S35.** Ancestral biomes reconstructed using a Hidden-State Dependent Speciation and Extinction (HiSSE) model that accounts for background heterogeneity in diversification rates across the phylogeny, by including a hidden trait with two states. Pies show the proportion of the three most probable ancestral states in 400 posterior samples. Ancestral states are shaded to represent relatively slow (dark) and relatively fast (light) background diversification. The analysis was conducted on the the maximum *a posteriori* (MAP) tree from the biogeographic dating analysis informed by previous estimates for the root age of Coenagrionoidea (Suvorov et al. 2021; Kohli et al. 2021). Species with distribution ranges spanning across two or three biomes were assigned to the coldest biome they inhabit.

**Table S13.** Comparison of biome-dependent diversification rates in Coenagrionoidea. We report mean differences and 95% highest posterior density (HPD) intervals. PMCMC values show the fraction of posterior samples in which the difference between rates and the posterior mean difference have the opposite signs. PMCMC values < 0.05 are considered evidence for biome-dependent diversification. Rates were estimated under a Hidden-State Dependent Speciation and Extinction (HiSSE) model, which accommodates background heterogeneity in diversification by modelling the diversification effects of a hidden trait with two states (see Extended Methods). The HiSSE analysis was conducted on the maximum *a posteriori* (MAP) tree from a biogeographic dating analyses informed by previous estimates for the root age of Coenagrionoidea (Suvorov et al. 2021; Kohli et al. 2021). Species with distribution ranges spanning across two or three biomes were coded as ambiguous between their current states.

| Rate | Biomes | Hidden state | Mean difference | Lower bound | Upper bound | PMCMC |
|---|---|---|---|---|---|---|
| Speciation | Cold-Temperate vs Warm-Temperate | 0 | 0.07 | 0.01 | 0.14 | 0.00 |
| Speciation | Cold-Temperate vs Warm-Temperate | 1 | 0.11 | 0.02 | 0.22 | 0.00 |
| Speciation | Tropical vs Cold-Temperate | 0 | -0.07 | -0.14 | -0.02 | 0.00 |
| Speciation | Tropical vs Cold-Temperate | 1 | -0.12 | -0.22 | -0.03 | 0.00 |
| Speciation | Tropical vs Warm-Temperate | 0 | 0.00 | -0.02 | 0.01 | 0.42 |
| Speciation | Tropical vs Warm-Temperate | 1 | 0.00 | -0.04 | 0.03 | 0.42 |
| Extinction | Cold-Temperate vs Warm-Temperate | 0 | 0.04 | -0.03 | 0.13 | 0.19 |
| Extinction | Cold-Temperate vs Warm-Temperate | 1 | 0.04 | -0.03 | 0.16 | 0.19 |
| Extinction | Tropical vs Cold-Temperate | 0 | -0.05 | -0.14 | 0.01 | 0.06 |
| Extinction | Tropical vs Cold-Temperate | 1 | -0.05 | -0.16 | 0.02 | 0.06 |
| Extinction | Tropical vs Warm-Temperate | 0 | -0.01 | -0.04 | 0.01 | 0.17 |
| Extinction | Tropical vs Warm-Temperate | 1 | -0.01 | -0.05 | 0.03 | 0.17 |
| Diversification | Cold-Temperate vs Warm-Temperate | 0 | 0.03 | -0.02 | 0.08 | 0.09 |
| Diversification | Cold-Temperate vs Warm-Temperate | 1 | 0.07 | -0.01 | 0.15 | 0.03 |
| Diversification | Tropical vs Cold-Temperate | 0 | -0.02 | -0.07 | 0.03 | 0.15 |
| Diversification | Tropical vs Cold-Temperate | 1 | -0.07 | -0.16 | 0.01 | 0.04 |
| Diversification | Tropical vs Warm-Temperate | 0 | 0.01 | -0.01 | 0.03 | 0.10 |
| Diversification | Tropical vs Warm-Temperate | 1 | 0.01 | -0.02 | 0.03 | 0.25 |
| Turnover | Cold-Temperate vs Warm-Temperate | 0 | 0.09 | -0.55 | 0.66 | 0.38 |
| Turnover | Cold-Temperate vs Warm-Temperate | 1 | 0.07 | -0.34 | 0.54 | 0.38 |

*(continued)*

| Rate | Biomes | Hidden state | Mean difference | Lower bound | Upper bound | PMCMC |
|---|---|---|---|---|---|---|
| Turnover | Tropical vs Cold-Temperate | 0 | -0.33 | -0.82 | 0.12 | 0.12 |
| Turnover | Tropical vs Cold-Temperate | 1 | -0.18 | -0.70 | 0.12 | 0.12 |
| Turnover | Tropical vs Warm-Temperate | 0 | -0.23 | -0.66 | 0.14 | 0.16 |
| Turnover | Tropical vs Warm-Temperate | 1 | -0.11 | -0.48 | 0.25 | 0.16 |

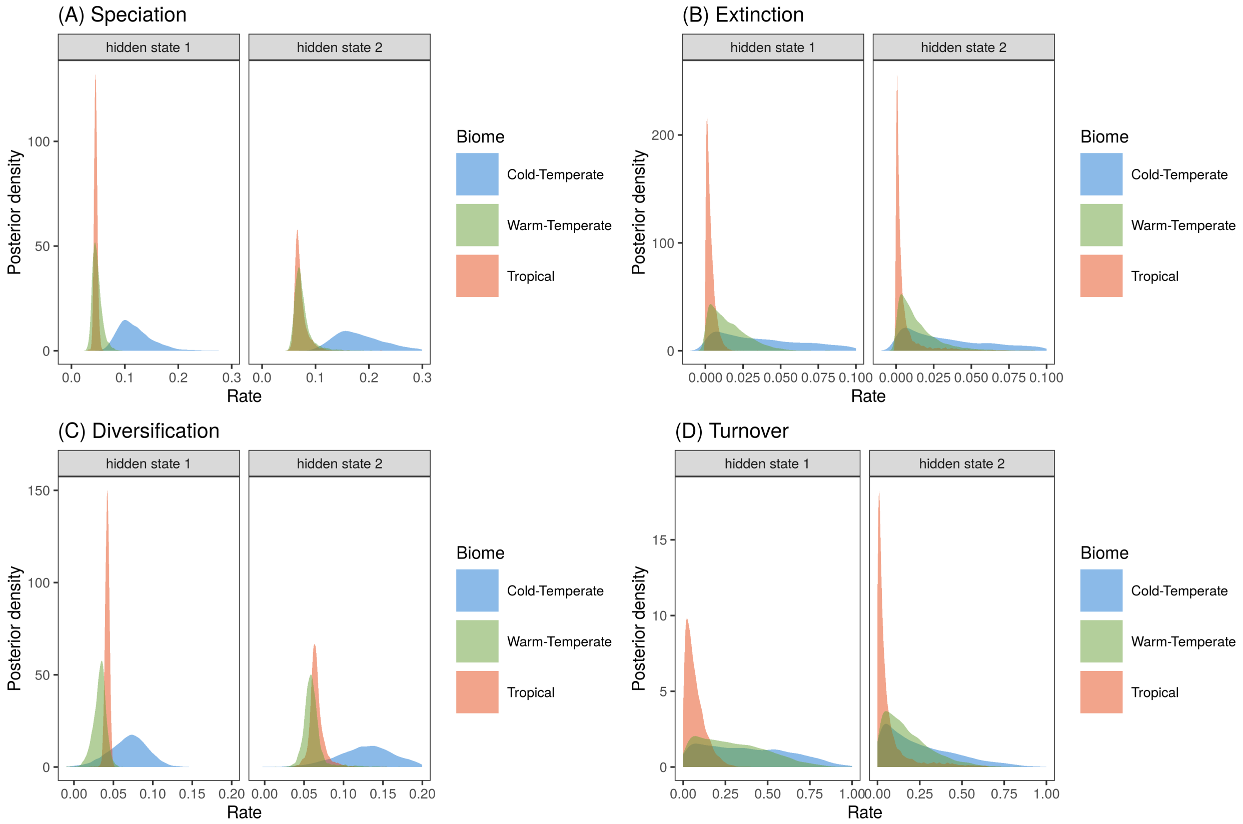

**Figure S36.** Biome effects on diversification of pond damselflies and their relatives the featherlegs (superfamily Coenagrionoidea). **(a)** Speciation and **(b)** extinction rates in tropical, warm-temperate, and cold-temperate biomes, were estimated using a Hidden-State Dependent Speciation and Extinction (HiSSE) model that accounts for background heterogeneity in diversification rates across the phylogeny, by including a hidden trait with two states. **(c)** Diversification was calculated as the net difference between speciation and extiction and **(d)** turnover was calculated as the ratio between extinction and speciation. The histograms show the posterior distribution of parameter estimates. The HiSSE analysis was conducted on the maximum *a posteriori* (MAP) tree from a biogeographic dating analyses informed by previous estimates for the root age of Coenagrionoidea (Suvorov et al. 2021; Kohli et al. 2021). Species with distribution ranges spanning across two or three biomes were coded as ambiguous between their current states.

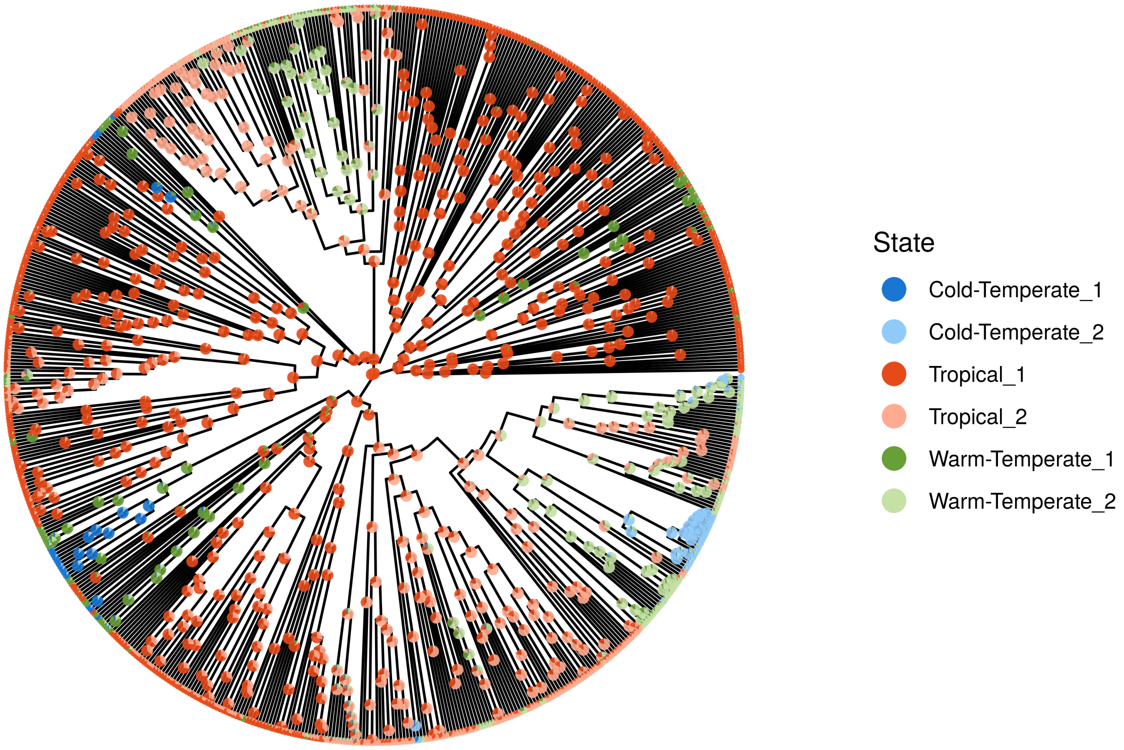

**Figure S37.** Ancestral biomes reconstructed using a Hidden-State Dependent Speciation and Extinction (HiSSE) model that accounts for background heterogeneity in diversification rates across the phylogeny, by including a hidden trait with two states. Pies show the proportion of the three most probable ancestral states in 4000 posterior samples. Ancestral states are shaded to represent relatively slow (dark) and relatively fast (light) background diversification. The analysis was conducted on the the maximum *a posteriori* (MAP) tree from the biogeographic dating analysis informed by previous estimates for the root age of Coenagrionoidea (Suvorov et al. 2021; Kohli et al. 2021). Species with distribution ranges spanning across two or three biomes were coded as ambiguous between their current states.

**Table S14.** Comparison of biome-dependent diversification rates in Coenagrionoidea. We report mean differences and 95% highest posterior density (HPD) intervals. PMCMC values show the fraction of posterior samples in which the difference between rates and the posterior mean difference have the opposite signs. PMCMC values < 0.05 are considered evidence for biome-dependent diversification. Rates were estimated under a Hidden-State Dependent Speciation and Extinction (HiSSE) model, which accommodates background heterogeneity in diversification by modelling the diversification effects of a hidden trait with two states (see Extended Methods). The HiSSE analysis was conducted on the maximum *a posteriori* (MAP) tree from a weakly-informed biogeographic dating analyses bounding the root age between 240 and 40 Ma. Species with distribution ranges spanning across two or three biomes were assigned to the coldest biome they inhabit.

| Rate | Biomes | Hidden state | Mean difference | Lower bound | Upper bound | PMCMC |
|---|---|---|---|---|---|---|
| Speciation | Cold-Temperate vs Warm-Temperate | 0 | 0.04 | 0.00 | 0.08 | 0.01 |
| Speciation | Cold-Temperate vs Warm-Temperate | 1 | 0.06 | 0.00 | 0.13 | 0.01 |
| Speciation | Tropical vs Cold-Temperate | 0 | -0.05 | -0.09 | -0.01 | 0.00 |
| Speciation | Tropical vs Cold-Temperate | 1 | -0.07 | -0.13 | -0.01 | 0.00 |
| Speciation | Tropical vs Warm-Temperate | 0 | 0.00 | -0.03 | 0.02 | 0.41 |
| Speciation | Tropical vs Warm-Temperate | 1 | -0.01 | -0.04 | 0.03 | 0.41 |
| Extinction | Cold-Temperate vs Warm-Temperate | 0 | 0.02 | -0.08 | 0.12 | 0.35 |
| Extinction | Cold-Temperate vs Warm-Temperate | 1 | 0.01 | -0.04 | 0.08 | 0.35 |
| Extinction | Tropical vs Cold-Temperate | 0 | -0.06 | -0.14 | 0.01 | 0.04 |
| Extinction | Tropical vs Cold-Temperate | 1 | -0.03 | -0.10 | 0.00 | 0.04 |
| Extinction | Tropical vs Warm-Temperate | 0 | -0.04 | -0.09 | 0.01 | 0.05 |
| Extinction | Tropical vs Warm-Temperate | 1 | -0.02 | -0.05 | 0.01 | 0.05 |
| Diversification | Cold-Temperate vs Warm-Temperate | 0 | 0.02 | -0.06 | 0.10 | 0.20 |
| Diversification | Cold-Temperate vs Warm-Temperate | 1 | 0.05 | 0.00 | 0.10 | 0.03 |
| Diversification | Tropical vs Cold-Temperate | 0 | 0.01 | -0.06 | 0.08 | 0.49 |
| Diversification | Tropical vs Cold-Temperate | 1 | -0.04 | -0.09 | 0.01 | 0.06 |
| Diversification | Tropical vs Warm-Temperate | 0 | 0.03 | -0.01 | 0.09 | 0.04 |
| Diversification | Tropical vs Warm-Temperate | 1 | 0.01 | -0.01 | 0.03 | 0.18 |
| Turnover | Cold-Temperate vs Warm-Temperate | 0 | -0.01 | -1.08 | 0.84 | 0.48 |
| Turnover | Cold-Temperate vs Warm-Temperate | 1 | 0.01 | -0.29 | 0.37 | 0.48 |
| Turnover | Tropical vs Cold-Temperate | 0 | -0.46 | -1.09 | 0.11 | 0.06 |

*(continued)*

| Rate | Biomes | Hidden state | Mean difference | Lower bound | Upper bound | PMCMC |
|---|---|---|---|---|---|---|
| Turnover | Tropical vs Cold-Temperate | 1 | -0.15 | -0.49 | 0.04 | 0.06 |
| Turnover | Tropical vs Warm-Temperate | 0 | -0.48 | -1.25 | 0.11 | 0.05 |
| Turnover | Tropical vs Warm-Temperate | 1 | -0.14 | -0.40 | 0.03 | 0.05 |

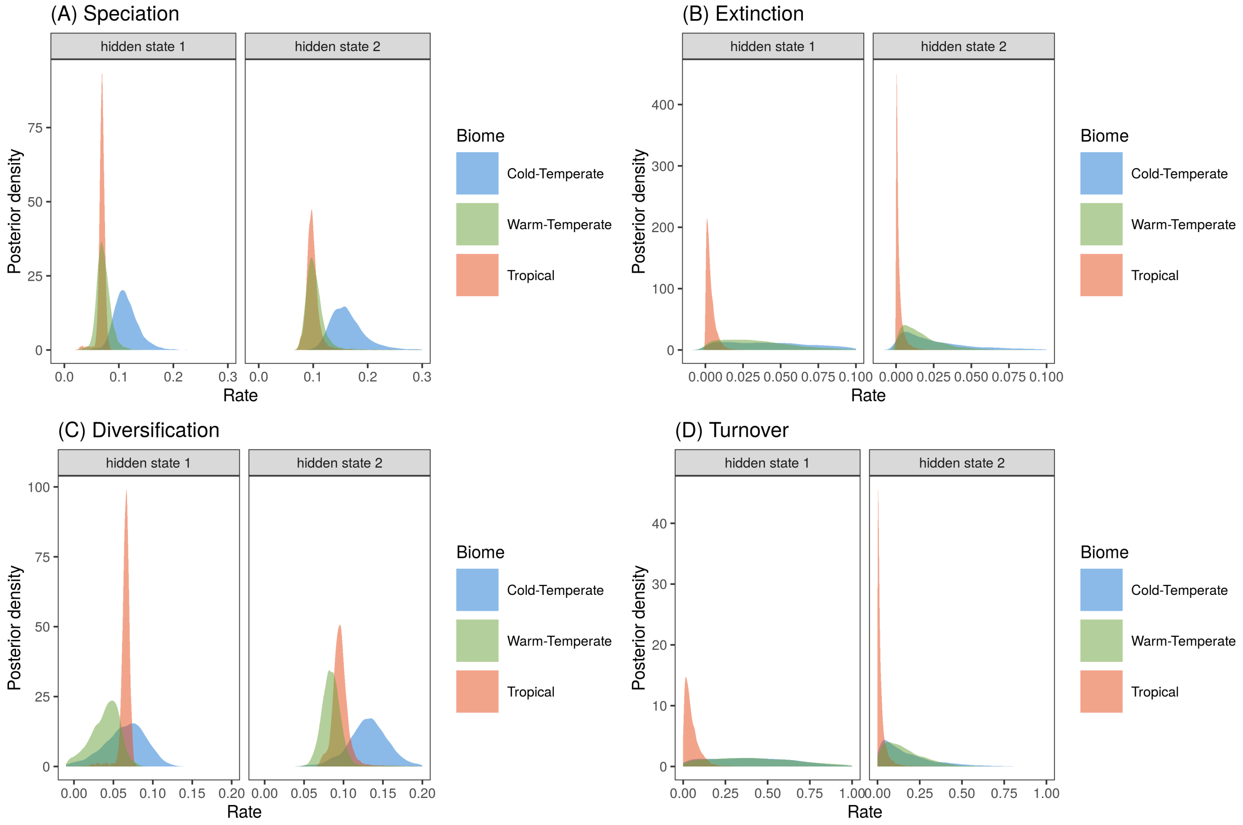

**Figure S38.** Biome effects on diversification of pond damselflies and their relatives the featherlegs (superfamily Coenagrionoidea). **(a)** Speciation and **(b)** extinction rates in tropical, warm-temperate, and cold-temperate biomes, were estimated using a Hidden-State Dependent Speciation and Extinction (HiSSE) model that accounts for background heterogeneity in diversification rates across the phylogeny, by including a hidden trait with two states. **(c)** Diversification was calculated as the net difference between speciation and extiction and **(d)** turnover was calculated as the ratio between extinction and speciation. The histograms show the posterior distribution of parameter estimates. The HiSSE analysis was conducted on the maximum *a posteriori* (MAP) tree from a weakly-informed biogeographic dating analyses bounding the root age between 240 and 40 Ma. Species with distribution ranges spanning across two or three biomes were assigned to the coldest biome they inhabit.

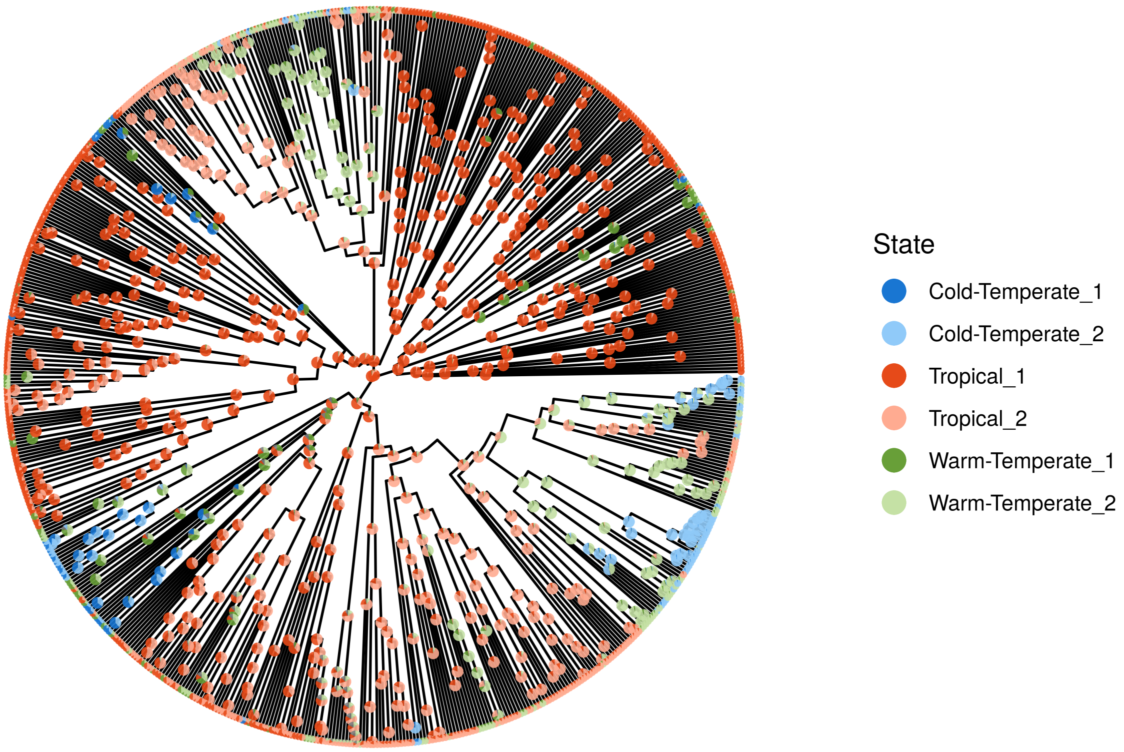

**Figure S39.** Ancestral biomes reconstructed using a Hidden-State Dependent Speciation and Extinction (HiSSE) model that accounts for background heterogeneity in diversification rates across the phylogeny, by including a hidden trait with two states. Pies show the proportion of the three most probable ancestral states in 4000 posterior samples. Ancestral states are shaded to represent relatively slow (dark) and relatively fast (light) background diversification. The analysis was conducted on the maximum *a posteriori* (MAP) tree from a weakly-informed biogeographic dating analyses bounding the root age between 240 and 40 Ma. Species with distribution ranges spanning across two or three biomes were assigned to the coldest biome they inhabit.

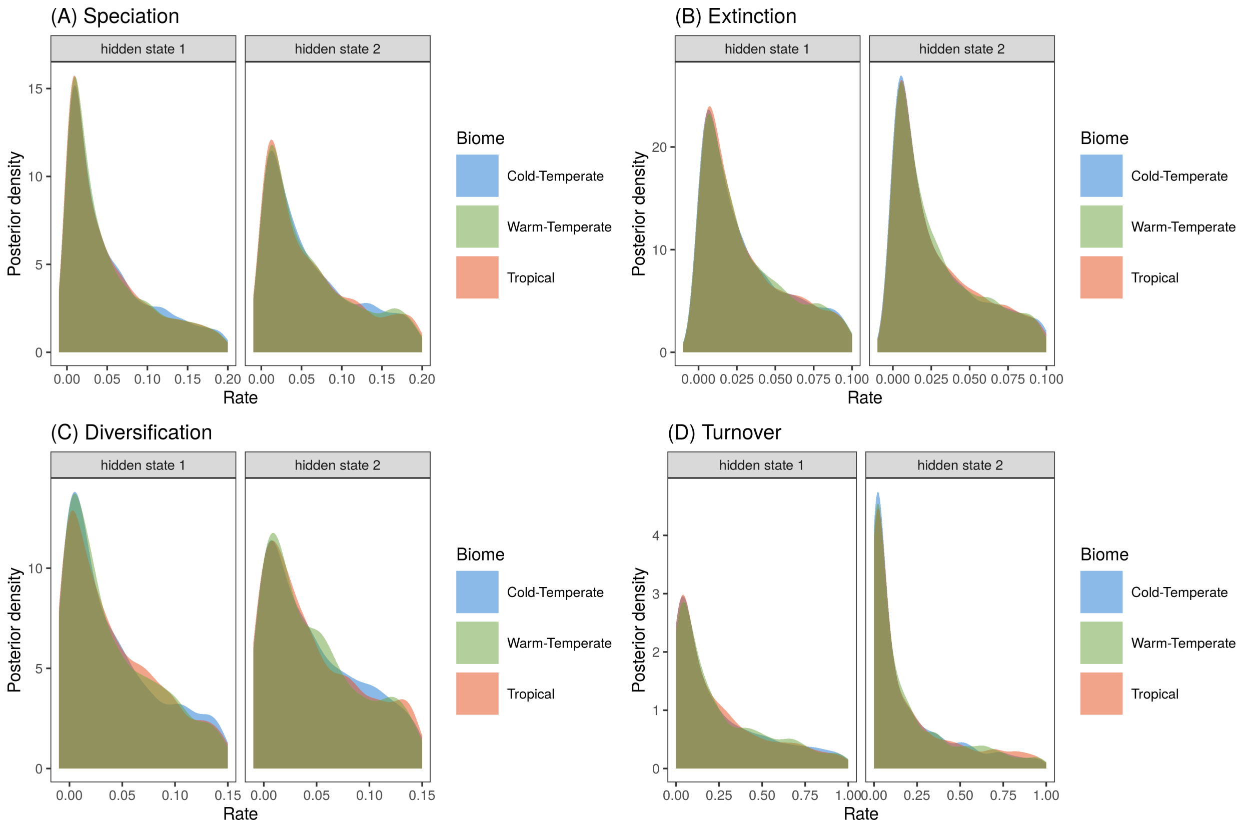

**Figure S40.** Hidden-State Dependent Speciation and Extinction (HiSSE) model of biome-dependent diversification run under the prior (i.e. without data). **(a)** Speciation and **(b)** extinction rates in tropical, warm-temperate, and cold-temperate biomes, were estimated while accounting for background heterogeneity in diversification rates, by including a hidden trait with two states. **(c)** Diversification was calculated as the net difference between speciation and extiction and **(d)** turnover was calculated as the ratio between extinction and speciation. The histograms show the posterior distribution of parameter estimates, which in this case recover the priors. The HiSSE analysis was conducted on the the maximum *a posteriori* (MAP) tree from the biogeographic dating analysis informed by previous estimates for the root age of Coenagrionoidea (Suvorov et al. 2021; Kohli et al. 2021).

#### Dispersal and biome-shift dynamics

We jointly modelled the history of dispersal events and biome shifts in Coenagrionoidea. In the main text, we show the proportions of lineages across biome-region states over time (Fig. 4), and the frequencies of different types of dispersal and biome-shift events in stochastically mapped character histories (Fig. 5). Here, we provide ancestral state reconstructions based on the joint dispersal and biome-shift model (Fig. S41), the posterior distributions of relative biome-shift rates (Fig. S42), and a visual assessment of congruence between inferred and available biomes throughout the history of the clade (Fig. S43).

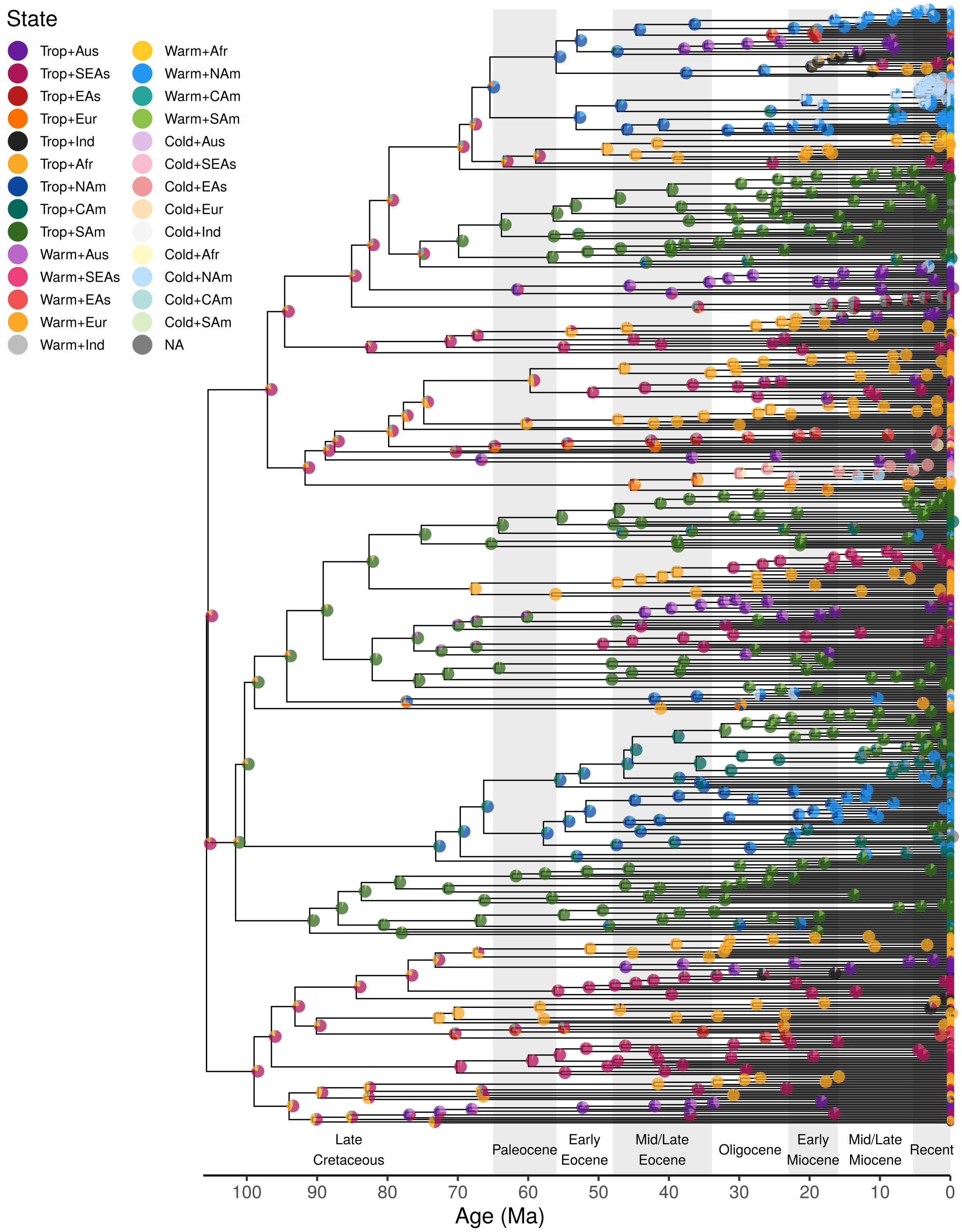

**Figure S41.** Ancestral ranges estimated using a time-heterogeneous continuous-time Markov model of biome-region shifts in Coenagrionoidea. Dispersal and biome-shift rates were estimated on the maximum *a posteriori* (MAP) tree from the biogeographic dating analysis, using an informed root age prior (dnNormal(120, 20)) based on previous studies (Suvorov et al. 2021; Kohli et al. 2021), and distribution data for extant taxa. The regions occupied by each of the extant taxa are shown at the tips of the tree. Pies show the proportion of the three most probable ancestral states in 3000 posterior samples.

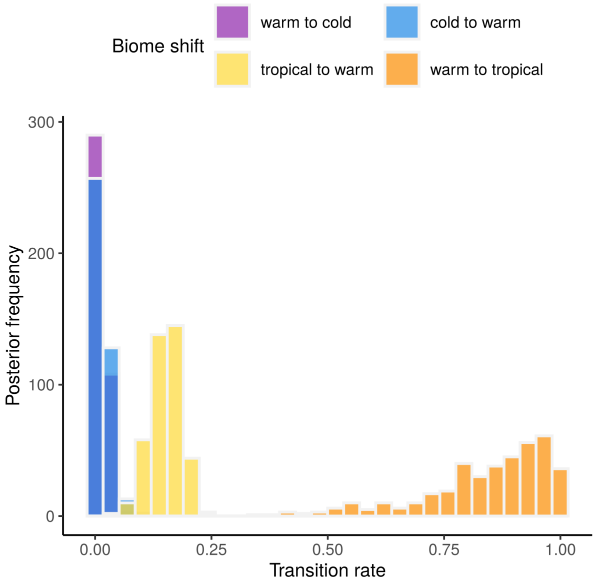

**Figure S42.** Relative transition rates between biome states in pond damselflies and their relatives the featherlegs (superfamily Coenagrionoidea). The histograms show the posterior distribution of parameter estimates. Relative transition rates were estimated using a time-heterogeneous continuous-time Markov model of biome-region shifts on the maximum *a posteriori* (MAP) tree from the biogeographic dating analysis using a an informed root age prior (dnNormal(120, 20)) based on previous studies (Suvorov et al. 2021; Kohli et al. 2021), and distribution data for extant taxa.

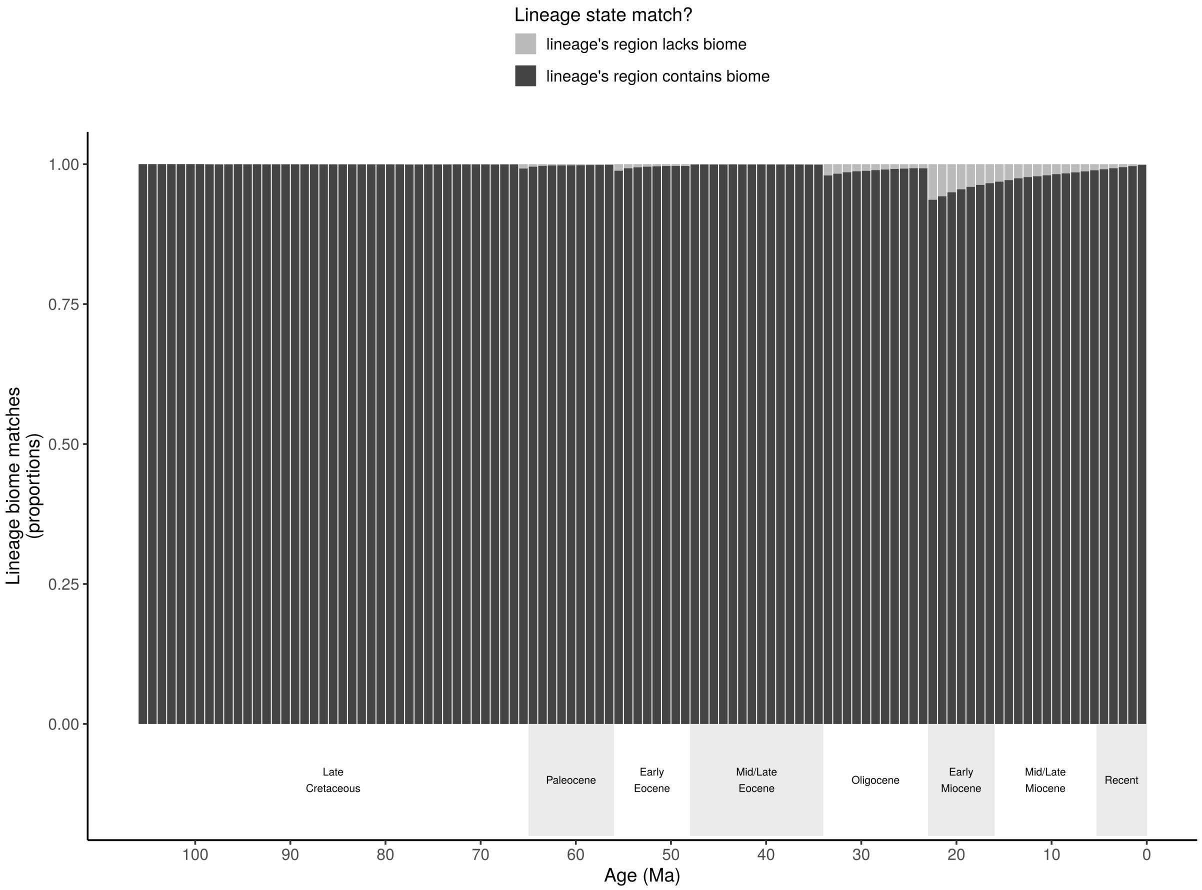

**Figure S43.** Congruence between inferred and available biomes in stochastic histories sampled in a time-heterogeneous model of biome-region shifts in Coenagrionoidea. We show the proportion of lineages with biome states that match (dark) or mismatch (light) locally accessible biomes (i.e. non-marginal biomes), at the lineage's geographic location. Accessible biomes are defined by an empirical paleobiome model (see Extended Methods for details).

### Literature cited

Barreda V., Palazzesi L. 2007. Patagonian vegetation turnovers during the Paleogene-early Neogene: Origin of arid-adapted floras. The Botanical Review. 73:31–50.

Beatty C.D., Sánchez Herrera M., Skevington J.H., Rashed A., Van Gossum H., Kelso S., Sherratt T.N. 2017. Biogeography and systematics of endemic island damselflies: The *Nesobasis* and *Melanesobasis* (Odonata: Zygoptera) of Fiji. Ecology and Evolution. 7:7117–7129.

Beaulieu J.M., O'meara B.C. 2016. Detecting hidden diversification shifts in models of trait-dependent speciation and extinction. Systematic Biology. 65:583–601.

Bechly G. 2000. A new fossil damselfly species(Insecta: Odonata: Zygoptera: Coenagrionidae:Ischnurinae) from Dominican Amber. Stuttgarter Beiträge zur Naturkunde Serie B (Geologie und Paläontologie).:1–9.

Bhatia H., Khan M.A., Srivastava G., Hazra T., Spicer R.A., Hazra M., Mehrotra R.C., Spicer T.E.V., Bera S., Roy K. 2021. Late Cretaceous–Paleogene Indian monsoon climate vis-à-vis movement of the Indian plate, and the birth of the South Asian Monsoon. Gondwana Research. 93:89–100.

Blow R., Willink B., Svensson E.I. 2021. A molecular phylogeny of forktail damselflies (genus *Ischnura*) reveals a dynamic macroevolutionary history of female colour polymorphisms. Molecular Phylogenetics and Evolution. 160:107134.

Bowman V.C., Francis J.E., Askin R.A., Riding J.B., Swindles G.T. 2014. Latest Cretaceous–earliest Paleogene vegetation and climate change at the high southern latitudes: Palynological evidence from seymour island, antarctic peninsula. Palaeogeography, Palaeoclimatology, Palaeoecology. 408:26–47.

Buerki S., Forest F., Stadler T., Alvarez N. 2013. The abrupt climate change at the Eocene–Oligocene boundary and the emergence of South-East Asia triggered the spread of sapindaceous lineages. Annals of Botany. 112:151–160.

Bybee S.M., Kalkman V.J., Erickson R.J., Frandsen P.B., Breinholt J.W., Suvorov A., Dijkstra K.-D.B., Cordero-Rivera A., Skevington J.H., Abbott J.C., others. 2021. Phylogeny and classification of Odonata using targeted genomics. Molecular Phylogenetics and Evolution. 160:107115.

Bybee S.M., Ogden T.H., Branham M.A., Whiting M.F. 2008. Molecules, morphology and fossils: A comprehensive approach to odonate phylogeny and the evolution of the odonate wing. Cladistics. 24:477–514.

Byrne M., Yeates D.K., Joseph L., Kearney M., Bowler J., Williams M.A.J., Cooper S., Donnellan S.C., Keogh J.S., Leys R., others. 2008. Birth of a biome: Insights into the assembly and maintenance of the Australian arid zone biota. Molecular Ecology. 17:4398.

Caesar R.M., Wenzel J.W. 2009. A phylogenetic test of classical species groups in *Argia* (Odonata: Coenagrionidae). Entomologica Americana. 115:97–108.

Callahan M.S., McPeek M.A. 2016. Multi-locus phylogeny and divergence time estimates of *Enallagma* damselflies (Odonata: Coenagrionidae). Molecular Phylogenetics and Evolution. 94:182–195.

Carle F.L., Kjer K.M., May M. 2008. Evolution of Odonata, with special reference to Coenagrionoidea (Zygoptera). Arthropod Systematics & Phylogeny. 66:37–44.

Carpenter R.J., Macphail M.K., Jordan G.J., Hill R.S. 2015. Fossil evidence for open, Proteaceae-dominated heathlands and fire in the Late Cretaceous of Australia. American Journal of Botany. 102:2092–2107.

Carvalho C.M., Polson N.G., Scott J.G. 2010. The horseshoe estimator for sparse signals. Biometrika. 97:465–480.

Chen J., Thomas D.C., Saunders R.M.K. 2019. Geographic range and habitat reconstructions shed light on palaeotropical intercontinental disjunction and regional diversification patterns in *Aartabotrys* (Annonaceae). Journal of Biogeography. 46:2690–2705.

Colgan D.J., McLauchlan A., Wilson G.D.F., Livingston S.P., Edgecombe G.D., Macaranas J., Cassis G., Gray M.R. 1998. Histone H3 and U2 snRNA DNA sequences and arthropod molecular evolution. Australian Journal of Zoology. 46:419–437.

De Marmels J. 1984. The genus *Nehalennia* Selys, its species and their phylogenetic relationships (Zygoptera: Coenagrionidae). Odonatologica. 13:501–527.

De Marmels J. 2002. A study of *Chromagrion* Needham, 1903, *Hesperagrion* Calvert, 1902, and *Zoniagrion* Kennedy, 1917: Three monotypic North American damselfly genera with uncertain generic relationships (Zygoptera: Coenagrionidae). Odonatologica. 31:139–150.

Dijkstra K.D.B. 2013. Three new genera of damselflies (Odonata: Chlorocyphidae, Platycnemididae). International Journal of Odonatology. 16:269–274.

Dijkstra K.D.B., Groeneveld L.F., Clausnitzer V., Hadrys H. 2007. The *Pseudagrion* split: Molecular phylogeny confirms the morphological and ecological dichotomy of Africa's most diverse genus of Odonata (Coenagrionidae). International Journal of Odonatology. 10:31–41.

Dijkstra K.D.B., Kalkman V.J., Dow R.A., Stokvis F.R., Van Tol J.A.N. 2014. Redefining the damselfly families: A comprehensive molecular phylogeny of Zygoptera (Odonata). Systematic Entomology. 39:68–96.

Dow R.A., Choong C.Y., Ng Y.F. 2010. A review of the genus *Amphicnemis* in Peninsular Malaysia and Singapore, with descriptions of two new species (Odonata: Zygoptera: Coenagrionidae). Zootaxa. 2605:45–55.

Dzombak R.M., Sheldon N.D., Mohabey D.M., Samant B. 2020. Stable climate in India during Deccan volcanism suggests limited influence on K–Pg extinction. Gondwana Research. 85:19–31.

Edgar R.C. 2004. MUSCLE: Multiple sequence alignment with high accuracy and high throughput. Nucleic Acids Research. 32:1792–1797.

Ferguson D.G., Marinov M., Saxton N.A., Rashni B., Bybee S.M. 2023. Phylogeny and classification of *Nesobasis* Selys, 1891 and *Vanuatubasis* Ober & Staniczek, 2009 (Odonata: Coenagrionidae). Insect Systematics & Evolution. 1:1–18.

Ferreira S., Lorenzo-Carballa M.O., Torres-Cambas Y., Cordero-Rivera A., Thompson D.J., Watts P.C. 2014. New EPIC nuclear DNA sequence markers to improve the resolution of phylogeographic studies of coenagrionids and other odonates. International Journal of Odonatology. 17:135–147.

FitzJohn R.G. 2012. Diversitree: Comparative phylogenetic analyses of diversification in R. Methods in Ecology and Evolution. 3:1084–1092.

Freyman W.A., Höhna S. 2019. Stochastic character mapping of state-dependent diversification reveals the tempo of evolutionary decline in self-compatible Onagraceae lineages. Systematic Biology. 68:505–519.

Garrison R.W. 2009. A synopsis of the genus *Telebasis* (Odonata: Coenagrionidae). International Journal of Odonatology. 12:1–121.

Garrison R.W., von Ellenrieder N. 2008. *Dolonagrion* nov. Gen. For *Telagrion fulvellum* from South America (Odonata: Coenagrionidae). International Journal of Odonatology. 11:173–183.

Garrison R.W., von Ellenrieder N. 2009. Redefinition of *Mesoleptobasis* Sjöstedt 1918 with the inclusion of *Metaleptobasis cyanolineata* (Wasscher 1998) comb. Nov. And description of a new species, *Mesoleptobasis elongata* (Odonata: Coenagrionidae). Zootaxa. 2145:47–68.

Garrison R.W., von Ellenrieder N. 2010. Redefinition of *Leptobasis* Selys with the synonymy of *Chrysobasis* Rácenis and description of *L. Mauffrayi* sp. Nov. From Peru (Odonata: Coenagrionidae). Zootaxa. 2438:1–36.

Garrison R.W., von Ellenrieder N., Louton J.A. 2010. Damselfly genera of the New World. Johns Hopkins University Press.

Gassmann D. 2005. The phylogeny of southeast Asian and indo-Pacific Calicnemiinae (Odonata, Platycnemididae). Bonner Zoologische Beiträge. 53:37–80.

Greenwood D.R., Moss P.T., Rowett A.I., Vadala A.J., Keefe R.L. 2003. Plant communities and climate change in southeastern australia during the early paleogene. Causes and consequences of globally warm climates in the Early Paleogene. 369:365.

Guan Z., Dumont H.J., Yu X., Han B.-P., Vierstraete A. 2013. *Pyrrhosoma* and its relatives: A phylogenetic study (Odonata: Zygoptera). International Journal of Odonatology. 16:247–257.

Guo Z.T., Sun B., Zhang Z.S., Peng S.Z., Xiao G.Q., Ge J.Y., Hao Q.Z., Qiao Y.S., Liang M.Y., Liu J.F., others. 2008. A major reorganization of Asian climate by the early Miocene. Climate of the Past. 4:153–174.

Herold N., Buzan J., Seton M., Goldner A., Green J.A.M., Müller R.D., Markwick P., Huber M. 2014. A suite of early Eocene (~ 55 ma) climate model boundary conditions. Geoscientific Model Development. 7:2077–2090.

Herold N., Huber M., Greenwood D., Müller R., Seton M. 2011. Early to middle Miocene monsoon climate in Australia. Geology. 39:3–6.

Höhna S. 2014. Likelihood inference of non-constant diversification rates with incomplete taxon sampling. PLoS ONE. 9:e84184.

Höhna S. 2015. The time-dependent reconstructed evolutionary process with a key-role for mass-extinction events. Journal of Theoretical Biology. 380:321–331.

Höhna S., Landis M.J., Heath T.A., Boussau B., Lartillot N., Moore B.R., Huelsenbeck J.P., Ronquist F. 2016. RevBayes: Bayesian phylogenetic inference using graphical models and an interactive model-specification language. Systematic Biology. 65:726–736.

Höhna S., Stadler T., Ronquist F., Britton T. 2011. Inferring speciation and extinction rates under different sampling schemes. Molecular Biology and Evolution. 28:2577–2589.

Ingley S.J., Bybee S.M., Tennessen K.J., Whiting M.F., Branham M.A. 2012. Life on the fly: Phylogenetics and evolution of the helicopter damselflies (Odonata, Pseudostigmatidae). Zoologica Scripta. 41:637–650.

Jacobs B.F. 2004. Palaeobotanical studies from tropical Africa: Relevance to the evolution of forest, woodland and savannah biomes. Philosophical Transactions of the Royal Society of London. Series B: Biological Sciences. 359:1573–1583.

Jacobs B.F., Pan A.D., Scotese Christopher.R., Werdelin L., Sanders W.J. 2010. A review of the Cenozoic vegetation history of Africa. University of California Press Berkeley, CA.

Jolly D., Harrison S.P., Damnati B., Bonnefille R. 1998. Simulated climate and biomes of Africa during the Late Quaternary: Comparison with pollen and lake status data. Quaternary Science Reviews. 17:629–657.

Jones H.B.C., Lim K.S., Bell J.R., Hill J.K., Chapman J.W. 2016. Quantifying interspecific variation in dispersal ability of noctuid moths using an advanced tethered flight technique. Ecology and Evolution. 6:181–190.

Jordan S., Simon C., Polhemus D. 2003. Molecular systematics and adaptive radiation of Hawaii's endemic damselfly genus *Megalagrion* (Odonata: Coenagrionidae). Systematic Biology. 52:89–109.

Kambhampati S. 1995. A phylogeny of cockroaches and related insects based on DNA sequence of mitochondrial ribosomal RNA genes. Proceedings of the National Academy of Sciences. 92:2017–2020.

Karube H., Futahashi R., Sasamoto A., Kawashima I. 2012. Taxonomic revision of Japanese odonate species, based on nuclear and mitochondrial gene genealogies and morphological comparison with allied species. Part I. Tombo. 54:75–106.

Kent D.V., Muttoni G. 2008. Equatorial convergence of India and early Cenozoic climate trends. Proceedings of the National Academy of Sciences. 105:16065–16070.

Kim M.J., Jung K.S., Park N.S., Wan X., Kim K.-G., Jun J., Yoon T.J., Bae Y.J., Lee S.M., Kim I. 2014. Molecular phylogeny of the higher taxa of Odonata (Insecta) inferred from COI, 16S rRNA, 28S rRNA, and EF1-a sequences. Entomological Research. 44:65–79.

Kjer K.M. 1995. Use of rRNA secondary structure in phylogenetic studies to identify homologous positions: An example of alignment and data presentation from the frogs. Molecular Phylogenetics and Evolution. 4:314–330.

Kjer K.M., Blahnik R.J., Holzenthal R.W. 2001. Phylogeny of Trichoptera (caddisflies): Characterization of signal and noise within multiple datasets. Systematic Biology. 50:781–816.

Klaus S., Morley R.J., Plath M., Zhang Y.-P., Li J.-T. 2016. Biotic interchange between the Indian subcontinent and mainland Asia through time. Nature Communications. 7:1–6.

Kohli M., Letsch H., Greve C., Béthoux O., Deregnaucourt I., Liu S., Zhou X., Donath A., Mayer C., Podsiadlowski L., others. 2021. Evolutionary history and divergence times of Odonata (dragonflies and damselflies) revealed through transcriptomics. Iscience. 24:103324.

Korasidis V.A., Wallace M.W., Wagstaff B.E., Hill R.S. 2019. Terrestrial cooling record through the Eocene-Oligocene transition of Australia. Global and Planetary Change. 173:61–72.

Landis M., Edwards E.J., Donoghue M.J. 2021. Modeling phylogenetic biome shifts on a planet with a past. Systematic Biology. 70:86–107.

Landis M.J. 2017. Biogeographic dating of speciation times using paleogeographically informed processes. Systematic Biology. 66:128–144.

Lencioni F.A.A. 1999. The genus *Phasmoneura*, with description of *Forcepsioneura* gen. nov. And two new species (Zygoptera: Protoneuridae). Odonatologica. 28:127–137.

Lim P.-E., Tan J., Eamsobhana P., Yong H.S. 2013. Distinct genetic clades of Malaysian *Copera* damselflies and the phylogeny of platycnemine subfamilies. Scientific Reports. 3:2977.

Löytynoja A., Goldman N. 2005. An algorithm for progressive multiple alignment of sequences with insertions. Proceedings of the National Academy of Sciences. 102:10557–10562.

Löytynoja A., Goldman N. 2008. Phylogeny-aware gap placement prevents errors in sequence alignment and evolutionary analysis. Science. 320:1632–1635.

Machado A. 2004. Studies on neotropical Protoneuridae. 15. *Amazoneura* gen. nov. With description of *A. Juruaensis* sp. Nov.(Odonata, Zygoptera). Revista Brasileira de Zoologia. 21:333–336.

Machado A.B.M. 2009. *Denticulobasis* and *Tuberculobasis*, new genera close to *Leptobasis*, with description of ten new species (Odonata: Coenagrionidae). Zootaxa. 2108:1–36.

Machado A.B.M., Lacerda D.S.S. 2017. Revalidation of *Platystigma* Kennedy, 1920, with a synopsis of the quadratum species group and the description of three new species (Odonata: Pseudostigmatidae). Zootaxa. 4242:493–516.

Maddison W.P. 2006. Confounding asymmetries in evolutionary diversification and character change. Evolution. 60:1743–1746.

Maddison W.P., FitzJohn R.G. 2014. The unsolved challenge to phylogenetic correlation tests for categorical characters. Systematic Biology. 64:127–136.

Maddison W.P., Midford P.E., Otto S.P. 2007. Estimating a binary character's effect on speciation and extinction. Systematic Biology. 56:701–710.

Magee A.F., Höhna S., Vasylyeva T.I., Leaché A.D., Minin V.N. 2020. Locally adaptive Bayesian birth-death model successfully detects slow and rapid rate shifts. PLoS Computational Biology. 16:e1007999.

Maley J. 1996. The African rain forest–main characteristics of changes in vegetation and climate from the Upper Cretaceous to the Quaternary. Proceedings of the Royal Society of Edinburgh, Section B: Biological Sciences. 104:31–73.

Marinov M., Amaya-Perilla C., Holwell G.I., Varsani A., Bysterveldt K., Kraberger S., Stainton D., Dayaram A., Curtis N., Cruickshank R.H., others. 2016. Geometric morphometrics and molecular systematics of *Xanthocnemis sobrina* (McLachlan, 1873)(Odonata: Coenagrionidae) and comparison to its congeners. Zootaxa. 4078:84–120.

Misof B., Liu S., Meusemann K., Peters R.S., Donath A., Mayer C., Frandsen P.B., Ware J., Flouri T., Beutel R.G., others. 2014. Phylogenomics resolves the timing and pattern of insect evolution. Science. 346:763–767.

Neumann F.H., Bamford M.K. 2015. Shaping of modern southern African biomes: Neogene vegetation and climate changes. Transactions of the Royal Society of South Africa. 70:195–212.

O'Brien C.L., Robinson S.A., Pancost R.D., Damsté J.S.S., Schouten S., Lunt D.J., Alsenz H., Bornemann A., Bottini C., Brassell S.C., others. 2017. Cretaceous sea-surface temperature evolution: Constraints from TEX86 and planktonic foraminiferal oxygen isotopes. Earth-Science Reviews. 172:224–247.

Ohba M., Ueda H. 2010. A GCM study on effects of continental drift on tropical climate at the early and late Cretaceous. Journal of the Meteorological Society of Japan. Ser. II. 88:869–881.

Orr A.G., Kalkman V.J., Richards S.J. 2012. A review of the New Guinean genus '*Paramecocnemis*' Lieftinck (Odonata: Platycnemididae), with the description of three new species. The Australian Entomologist. 39:161–177.

Otto-Bliesner B.L., Brady E.C., Tomas R.A., Albani S., Bartlein P.J., Mahowald N.M., Shafer S.L., Kluzek E., Lawrence P.J., Leguy G., others. 2020. A comparison of the CMIP6 midHolocene and lig127k simulations in CESM2. Paleoceanography and Paleoclimatology. 35:e2020PA003957.

Otto-Bliesner B.L., Upchurch G.R. 1997. Vegetation-induced warming of high-latitude regions during the Late Cretaceous period. Nature. 385:804–807.

Paradis E., Schliep K. 2019. *ape* 5.0: An environment for modern phylogenetics and evolutionary analyses in R. Bioinformatics. 35:526–528.

Parham J.F., Donoghue P.C.J., Bell C.J., Calway T.D., Head J.J., Holroyd P.A., Inoue J.G., Irmis R.B., Joyce W.G., Ksepka D.T., others. 2012. Best practices for justifying fossil calibrations. Systematic Biology. 61:346–359.

Paulson D., Schorr M. 2021. World Odonata List. Available from: https://www.pugetsound.edu/academics/academic-resources/slater-museum/biodiversity-resources/dragonflies/world-odonata-list2/.

Pessacq P. 2008. Phylogeny of Neotropical Protoneuridae (Odonata: Zygoptera) and a preliminary study of their relationship with related families. Systematic Entomology. 33:511–528.

Pessacq P. 2014. Synopsis of *Epipleoneura* (Zygoptera, Coenagrionidae,"Protoneuridae"), with emphasis on its Brazilian morphospecies. Zootaxa. 3872:201–234.

Pimenta A.L.A., Pinto Â., Takiya D. 2019. Integrative taxonomy and phylogeny of the damselfly genus *Forcepsioneura* Lencioni, 1999 (Odonata: Coenagrionidae: Protoneurinae) with description of two new species from the Brazilian Atlantic Forest. Arthropod Systematics & Phylogeny. 77:379–415.

Plummer M., Best N., Cowles K., Vines K. 2006. CODA: Convergence Diagnosis and Output Analysis for MCMC. R News. 6:7–11.

Poinar Jr G., Bechly G., Buckley R. 2010. First record of Odonata and a new subfamily of damselflies from Early Cretaceous Burmese amber. Palaeodiversity. 3:e22.

Potter P.E., Szatmari P. 2009. Global Miocene tectonics and the modern world. Earth-Science Reviews. 96:279–295.

Pound M.J., Haywood A.M., Salzmann U., Riding J.B. 2012. Global vegetation dynamics and latitudinal temperature gradients during the Mid to Late Miocene (15.97–5.33 Ma). Earth-Science Reviews. 112:1–22.

Pound M.J., Haywood A.M., Salzmann U., Riding J.B., Lunt D.J., Hunter S.J. 2011. A Tortonian (late Miocene, 11.61–7.25 Ma) global vegetation reconstruction. Palaeogeography, Palaeoclimatology, Palaeoecology. 300:29–45.

Pound M.J., Salzmann U. 2017. Heterogeneity in global vegetation and terrestrial climate change during the late Eocene to early Oligocene transition. Scientific Reports. 7:1–12.

Prasad V., Utescher T., Sharma A., Singh I., Garg R., Gogoi B., Srivastava J., Uddandam P., Joachimski M. 2018. Low-latitude vegetation and climate dynamics at the Paleocene-Eocene transition – A study based on multiple proxies from the Jathang section in northeastern India. Palaeogeography, Palaeoclimatology, Palaeoecology. 497:139–156.

Pross J., Contreras L., Bijl P.K., Greenwood D.R., Bohaty S.M., Schouten S., Bendle J.A., Röhl U., Tauxe L., Raine J.I., others. 2012. Persistent near-tropical warmth on the Antarctic continent during the early Eocene epoch. Nature. 488:73–77.

R Core Team. 2021. R: A language and environment for statistical computing. Vienna, Austria: R Foundation for Statistical Computing.

Rabosky D.L., Goldberg E.E. 2015. Model inadequacy and mistaken inferences of trait-dependent speciation. Systematic Biology. 64:340–355.

Roberts D.L., Neumann F.H., Cawthra H.C., Carr A.S., Scott L., Durugbo E.U., Humphries M.S., Cowling R.M., Bamford M.K., Musekiwa C., others. 2017. Palaeoenvironments during a terminal Oligocene or early Miocene transgression in a fluvial system at the southwestern tip of Africa. Global and Planetary Change. 150:1–23.

Ross A.J., Coutiño José M.A., Nel A. 2016. The first records of coenagrionid damselflies (Odonata: Zygoptera: Coenagrionidae: *Neoerythromma* sp. and *Nehalennia* sp.) From Mexican Amber (Miocene). Boletín de la Sociedad Geológica Mexicana. 68:81–86.

Rundel P.W., Arroyo M.T., Cowling R.M., Keeley J.E., Lamont B.B., Vargas P., others. 2016. Mediterranean biomes: Evolution of their vegetation, floras, and climate. Annual Review of Ecology, Evolution, and Systematics. 47:383–407.

Salzmann U., Haywood A.M., Lunt D.J., Valdes P.J., Hill D.J. 2008. A new global biome reconstruction and data-model comparison for the Middle Pliocene. Global Ecology and Biogeography. 17:432–447.

Saxton N.A., Marinov M.G., Bybee S.M. 2022. Revision of *Vanuatubasis* Ober & Staniczek, 2009 (Odonata, Coenagrionidae), with description of seven new morphospecies. ZooKeys.

Selys-Longchamps E. de. 1876. Synopsis des Agrionines, 5me légion: *Agrion* (suite). Le genre *Agrion*. Bulletins de l'Académie royale des sciences, des lettres et des beaux-arts de Belgique. 41:247–322.

Singh H., Prasad M., Kumar K., Singh S.K. 2011. Paleobotanical remains from the Paleocene–lower Eocene Vagadkhol Formation, western India and their paleoclimatic and phytogeographic implications. Palaeoworld. 20:332–356.

Specht R.L., Dettmann M.E., Jarzen D.M. 1992. Community associations and structure in the Late Cretaceous vegetation of southeast Australasia and Antarctica. Palaeogeography, Palaeoclimatology, Palaeoecology. 94:283–309.

Srivastava G., Spicer R.A., Spicer T.E.V., Yang J., Kumar M., Mehrotra R., Mehrotra N. 2012. Megaflora and palaeoclimate of a Late Oligocene tropical delta, Makum Coalfield, Assam: Evidence for the early development of the South Asia Monsoon. Palaeogeography, Palaeoclimatology, Palaeoecology. 342:130–142.

Stadler T. 2011. Mammalian phylogeny reveals recent diversification rate shifts. Proceedings of the National Academy of Sciences. 108:6187–6192.

Su T., Spicer R.A., Li S.-H., Xu H., Huang J., Sherlock S., Huang Y.-J., Li S.-F., Wang L., Jia L.-B., others. 2019. Uplift, climate and biotic changes at the Eocene–Oligocene transition in south-eastern Tibet. National Science Review. 6:495–504.

Suvorov A., Scornavacca C., Fujimoto M.S., Bodily P., Clement M., Crandall K.A., Whiting M.F., Schrider D.R., Bybee S.M. 2021. Deep ancestral introgression shapes evolutionary history of dragonflies and damselflies. Systematic Biology.:syab063.

Swaegers J., Janssens S.B., Ferreira S., Watts P.C., Mergeay J., McPeek M.A., Stoks R. 2014. Ecological and evolutionary drivers of range size in *Coenagrion* damselflies. Journal of Evolutionary Biology. 27:2386–2395.

Tennessen K.J. 2009. *Aeolagrion philipi* sp. nov. from Bolivia, and a review of the genus *Aeolagrion* (Odonata: Coenagrionidae). International Journal of Odonatology. 12:309–411.

Tennessen K.J. 2015. Four new species of *Calvertagrion* St. Quentin from South America (Odonata: Coenagrionidae). Odonatologica. 44:397–430.

Torres-Pachón M., Novelo-Gutiérrez R., Espinosa de los Monteros A. 2017. Phylogenetic analysis of the genus *Argia* Rambur, 1842 (Odonata: Coenagrionidae), based on morphological characters of larvae and mitochondrial DNA sequences. Organisms Diversity & Evolution. 17:409–420.

Toussaint E.F.A., Bybee S.M., Erickson R.J., Condamine F.L. 2019. Forest giants on different evolutionary branches: Ecomorphological convergence in helicopter damselflies. Evolution. 73:1045–1054.

Turgeon J., Stoks R., Thum R.A., Brown J.M., McPeek M.A. 2005. Simultaneous Quaternary radiations of three damselfly clades across the Holarctic. The American Naturalist. 165:E78–E107.

Utescher T., Mosbrugger V. 2007. Eocene vegetation patterns reconstructed from plant diversity—a global perspective. Palaeogeography, Palaeoclimatology, Palaeoecology. 247:243–271.

Vilela D.S., Anjos-Santos D., Koroiva R., Cordero-Rivera A., Guillermo-Ferreira R. 2020. Revision of the genus *Minagrion* Santos, 1965 (Odonata: Coenagrionidae). Zootaxa. 4786:zootaxa–4786.

Villanueva R.J.T. 2012. Review of the Philippine taxa formerly assigned to the genus *Amphicnemis* Selys. Part I: Overview and descriptions of three new genera (Odonata: Coenagrionidae). Zoologische Mededelingen. 86.

Vincens A., Tiercelin J.-J., Buchet G. 2006. New Oligocene–early Miocene microflora from the southwestern Turkana Basin: Palaeoenvironmental implications in the northern kenya rift. Palaeogeography, Palaeoclimatology, Palaeoecology. 239:470–486.

von Ellenrieder N. 2008a. Revalidación de *Argentagrion* y redefinición de *Homeoura*, con la descripción de *H. Obrieni* n. sp.(Odonata: Coenagrionidae). Revista de la Sociedad entomológica Argentina. 67:81–106.

von Ellenrieder N. 2008b. *Phoenicagrion* gen. nov. for *Leptagrion flammeum*, with description of a new species, *P. Paulsoni*, from Peru (Odonata: Coenagrionidae). International Journal of Odonatology. 11:81–93.

von Ellenrieder N. 2013. A revision of *Metaleptobasis* Calvert (Odonata: Coenagrionidae) with seven synonymies and the description of eighteen new species from South America. Zootaxa. 3738:1–155.

von Ellenrieder N., Garrison R.W. 2008a. *Oreiallagma* gen. nov. with a redefinition of *Cyanallagma* Kennedy 1920 and *Mesamphiagrion* Kennedy 1920, and the description of *M. Dunklei* sp. nov. and *M. Ecuatoriale* sp. nov. from Ecuador (Odonata: Coenagrionidae). Zootaxa.

von Ellenrieder N., Garrison R.W. 2008b. *Drepanoneura* gen. Nov. For *Epipleoneura letitia* and *Protoneura peruviensis*, with descriptions of eight new Protoneuridae from South America (Odonata: Protoneuridae). Zootaxa. 1842:1–34.

von Ellenrieder N., Garrison R.W. 2008c. A redefinition of *Telagrion* Selys and *Aceratobasis* Kennedy stat. rev. and the description of *Schistolobos* gen. nov. for *Telagrion boliviense* Daigle (Odonata: Coenagrionidae).

Transactions of the American Entomological Society. 134:1–22.

von Ellenrieder N., Lozano F. 2008. Blues for the red *Oxyagrion*: A redefinition of the genera *Acanthagrion* and *Oxyagrion* (Odonata: Coenagrionidae). International Journal of Odonatology. 11:95–113.

Waller J.T., Svensson E.I. 2017. Body size evolution in an old insect order: No evidence for Cope's Rule in spite of fitness benefits of large size. Evolution. 71:2178–2193.

Waller J.T., Willink B., Tschol M., Svensson E.I. 2019. The odonate phenotypic database, a new open data resource for comparative studies of an old insect order. Scientific Data. 6:1–6.

Ware J., May M., Kjer K. 2007. Phylogeny of the higher Libelluloidea (Anisoptera: Odonata): An exploration of the most speciose superfamily of dragonflies. Molecular Phylogenetics and Evolution. 45:289–310.

Warny S., Jarzen D.M., Haynes S.J., MacLeod K.G., Huber B.T. 2019. Late Cretaceous (Turonian) angiosperm pollen from Tanzania: A glimpse of past vegetation from a warmer climate. Palynology. 43:608–620.

Weekers P.H.H., Dumont H.J. 2004. A molecular study of the relationship between the coenagrionid genera *Erythromma* and *Cercion*, with the creation of *Paracercion* gen. nov. for the East Asiatic "*Cercion*" (Zygoptera: Coenagrionidae). Odonatologica. 33:181–188.

Westerhold T., Marwan N., Drury A.J., Liebrand D., Agnini C., Anagnostou E., Barnet J.S.K., Bohaty S.M., De Vleeschouwer D., Florindo F., others. 2020. An astronomically dated record of Earth's climate and its predictability over the last 66 million years. Science. 369:1383–1387.

Willink B., Duryea M.C., Svensson E.I. 2019. Macroevolutionary origin and adaptive function of a polymorphic female signal involved in sexual conflict. The American Naturalist. 194:707–724.

Xia X., Lemey P. 2009. The phylogenetic handbook: A practical approach to DNA and protein phylogeny. In: Lemey P., Salemi M., Vandamme A.-M., editors. The phylogenetic handbook: a practical approach to DNA and protein phylogeny. New York: Cambridge University Press Cambridge. p. 615–630.

Xia X., Xie Z. 2001. DAMBE: Software package for data analysis in molecular biology and evolution. Journal of Heredity. 92:371–373.

Xie W., Lewis P.O., Fan Y., Kuo L., Chen M.-H. 2010. Improving marginal likelihood estimation for Bayesian phylogenetic model selection. Systematic Biology. 60:150–160.

Xiong B., Kocher T.D. 1991. Comparison of mitochondrial DNA sequences of seven morphospecies of black flies (Diptera: Simuliidae). Genome. 34:306–311.

Yang Z. 2006. Computational molecular evolution. Oxford, UK: Oxford University Press.

Zheng D., Nel A., Jarzembowski E.A., Chang S.-C., Zhang H., Wang B. 2018. Exceptionally well-preserved dragonflies (Insecta: Odonata) in Mexican amber. Alcheringa: An Australasian Journal of Palaeontology. 43:157–164.
